# A new label-free optical imaging method for the lymphatic system enhanced by deep learning

**DOI:** 10.1101/2023.01.13.523938

**Authors:** Zhongming Li, Shengnan Huang, Yanpu He, Jan Willem van Wijnbergen, Yizhe Zhang, Rob D. Cottrell, Sean G. Smith, Paula T. Hammond, Danny Z. Chen, Timothy P. Padera, Angela M. Belcher

## Abstract

Our understanding of the lymphatic vascular system lags far behind that of the blood vascular system, limited by available imaging technologies. We present a label-free optical imaging method that visualizes the lymphatic system with high contrast. We developed an orthogonal polarization imaging (OPI) in the shortwave infrared range (SWIR) and imaged both lymph nodes and lymphatic vessels of mice and rats *in vivo* through intact skin, as well as human mesenteric lymph nodes in colectomy specimens. By integrating SWIR-OPI with U-Net, a deep learning image segmentation algorithm, we automated the lymph node size measurement process. Changes in lymph nodes in response to cancer progression were monitored in two separate mouse cancer models, through which we obtained insights into pre-metastatic niches and correlation between lymph node masses and many important biomarkers. In a human pilot study, we demonstrated the effectiveness of SWIR-OPI to detect human lymph nodes in real time with clinical colectomy specimens.

**One Sentence Summary:** We develop a real-time high contrast optical technique for imaging the lymphatic system, and apply it to anatomical pathology gross examination in a clinical setting, as well as real-time monitoring of tumor microenvironment in animal studies.

## INTRODUCTION

Cancer cells metastasize through the vascular or lymphatic system to colonize distant organs. Tumor draining lymph nodes are often the first sites to be colonized. In multiple solid tumors, the presence of cancer cells in nearby lymph nodes is a strong predictor of poor outcome.(*1-6*) Additionally, cancer cells in metastatic lymph nodes are able to escape the node and seed distant metastatic sites.(*7, 8*) However, given the critical clinical significance of the lymphatic system, our understanding of it lags far behind that of the vascular system, mainly limited by the capabilities of available technologies to study the lymphatic system *in vivo*.(*9*) Common imaging modalities, such as scintography,(*10*) computed tomography (CT),(*11*) magnetic resonance imaging (MRI),(*12*) ultrasound (US),(*13*) positron emission tomography (PET),(*14*) or a combination of these,(*15, 16*) generally lack satisfactory tissue contrast and spatial resolution.(*17*) Contrast agents are often injected to mitigate the contrast limitation and provide important tumor draining information. However, sufficiently high spatial resolution still remains a challenge, and contrast agents pose potential health hazards to patients and may also alter the function of the lymphatic system.(*17, 18*)

Optical imaging, on the other hand, offers superior spatial resolution. Blue dyes(*19*) and near-infrared fluorescent dyes(*20*) have been widely used for lymphography in the clinical setting,(*21*) especially during lymphatic-venous bypass surgeries, when spatial precision is of critical importance.(*22, 23*) These optical imaging modalities, though free from radioactive reagents or processes, still require a laborious dye injection procedure, and the injected dyes can still pose potential health hazards to patients.(*24, 25*) The applications of such methods are constrained by tissue depth, limiting their use to superficial tissues or those that can be made accessible. Practically, dye-injection based lymphatic imaging techniques are limited to the evaluation of superficial lymphatic drainage in lymphedema patients(*21*) or in breast cancer, melanoma, and head and neck cancer patients, because cancers in other parts of the body are either too hard to access or may have lymphatic networks that are too interconnected for dyes to delineate the features of interest.(*26, 27*) Optical coherence tomography (OCT),(*28*) as a label-free optical imaging method commonly used for vasculature imaging, has been also been used for imaging the lymphatic system.(*29, 30*) However, OCT has a limited field-of-view and involves a sophisticated laser scanning technique that makes this method slow and inconvenient to use for clinical lymphatic imaging.

Biomedical imaging in the SWIR range has been demonstrated to be a powerful tool for deep tissue imaging.(*31*) However, most SWIR biomedical imaging methods are based on fluorescence(*32-34*) or photoluminescence(*35*) Label-free SWIR imaging based on light absorption and scattering is an under-explored area with very limited previous research.(*36*) Bawendi and coworkers previously demonstrated that label-free SWIR imaging can be used for middle ear inspection.(*37*) Recently, Roblyer and coworkers demonstrated SWIR meso-patterned imaging (SWIR-MPI), which provides a label-free, non-contact solution for spatial mapping of water and lipid concentrations in tissue.(*38*) Compared to how the natural optical absorption of hemoglobin in the visible range gave rise to a number of widely used clinical imaging modalities for visualizing blood vasculature,(*39-41*) the natural optical absorptions of water, lipid, collagen, and other biologically important chemicals in the SWIR range are vastly under-explored.(*42*) These previous forays into label-free SWIR imaging together with our work are likely to mark the inception of a new field of label-free SWIR imaging for a range of biomedical applications.

Here, we present a label-free, high-contrast optical imaging method that visualizes lymph nodes and lymphatic vessels in real-time with high spatial resolution. We use orthogonal polarization imaging (OPI) in the shortwave infrared (SWIR) range, which exploits the natural chemical composition contrast between lymph nodes and lymphatic vessels and the surrounding fat tissue. OPI, also known as cross-polarization imaging, can increase the imaging penetration depth,(*39*) because conventionally it prevents photons that are reflected directly off the surface of imaged objects from reaching the camera sensor. It is particularly effective for improving the imaging penetration depth in the SWIR range, because the scattering effect in the SWIR domain is intrinsically greatly reduced compared to that of the visible light. OPI can thereby not only remove photons that reflect directly off the surface of biological tissue, but also the photons that back-scatter from shallower layers of tissue.

## RESULTS

### SWIR-OPI imaging system

We first demonstrate the effectiveness of a SWIR-OPI imaging system with commonly used rodent models – BALB/c mice and Wistar IGS rats. We prepare the BALB/c mice for imaging sessions by removing the hair from the groin and hind limbs (see Method). The mice are anesthetized with isoflurane to reduce motion artifacts during the imaging sessions. We use a linear polarizer in front of a collimated 1550 nm LED to create a linearly polarized illumination (Figure 1(a)). We then use another linear polarizer in front of the imaging lens, which is adjusted to be orthogonal to the direction of the linearly polarized illumination. We also developed a home-built SWIR object-space telecentric lens to improve the accuracy of image-based lymph node size measurements. We used a liquid nitrogen-cooled, scientific-grade InGaAs camera (Princeton Instruments, 2D-OMA V:320), as well as an affordable, commercial-grade InGaAs camera (SU320M-1.7RT, Sensors Unlimited), for detection.

**Figure 1.**
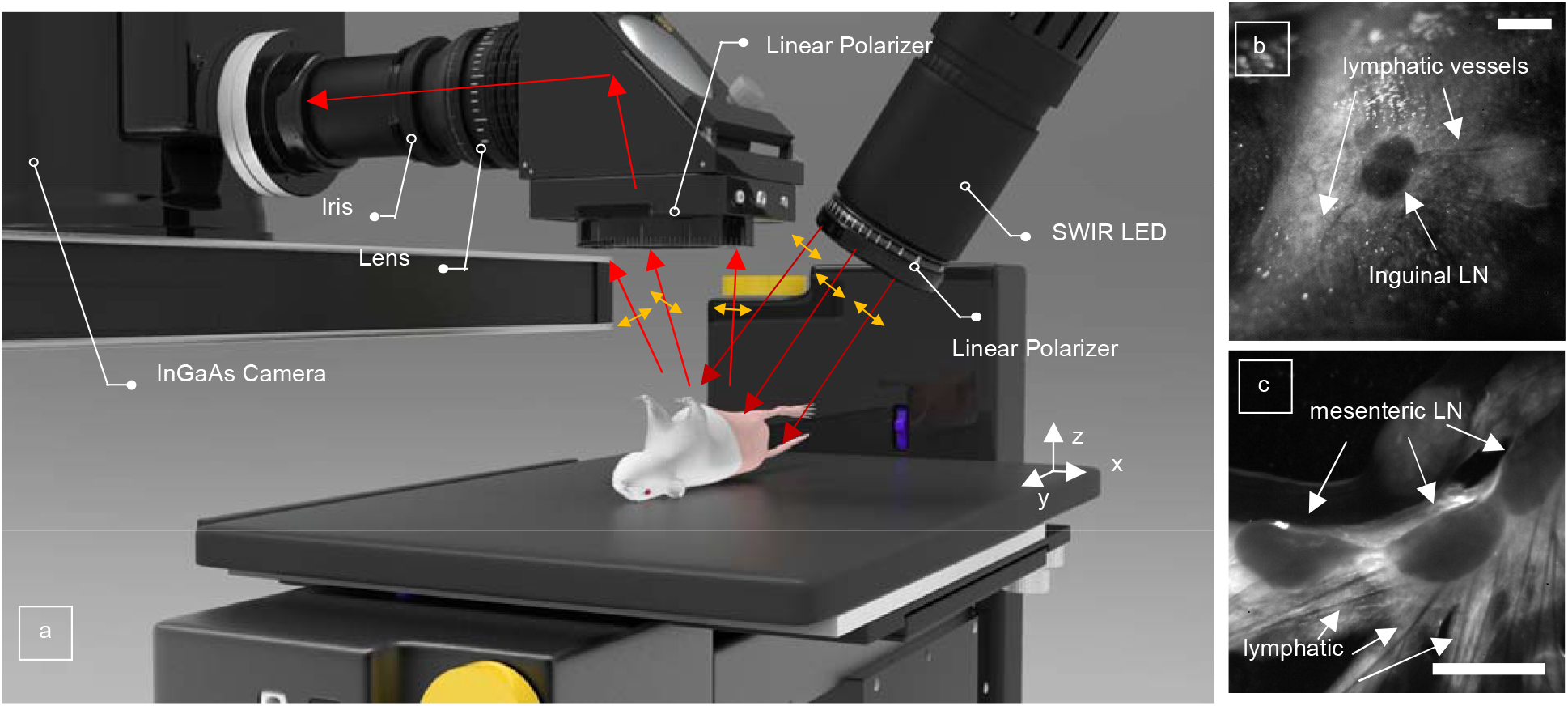
(a) Schematic diagram of Shortwave Infrared Orthogonal Polarization Imaging (SWIR-OPI); (b) mouse inguinal LN through intact mouse skin; (c) mouse mesenteric LN. Lymphatic vessels can also be visualized clearly both *ex-vivo* near the mesenteric LNs and *in-vivo* near the inguinal LN noninvasively. Scale bar 2 mm.

With this imaging setup, we visualized the inguinal lymph nodes of BALB/c mice through intact skin, as shown in Figure 1(b). The inguinal lymph node and surrounding lymphatic vessels can be seen clearly with high contrast. The spatial resolution was measured to be 65 μm (see Supplementary Information). This resolution can be further improved by using imaging lenses with higher magnifications. The spatial resolution that we obtained was a product of optimizing different optical imaging parameters, such as field-of-view, depth-of-field, and telecentricity, for the specific use case of imaging mouse inguinal lymph nodes *in vivo* (see Supplementary Information). We also imaged murine mesenteric lymph nodes surrounded by fat tissue *ex vivo*, as shown in Figure 1(c). The spatial resolution was measured to be approximately 80 μm. It is worth noting that SWIR-OPI works with live animals *in vivo*, as well as with *ex vivo* specimens, whereas dye injection-based imaging methods require a living, functional lymphatic system to actively drain the tracer molecules in order for the lymphatic system to be visualized.

There are three key parts of the SWIR-OPI system that provide the unique capability of visualizing lymph nodes noninvasively: wavelength, polarization, and an object-space telecentric SWIR lens.

As shown in Figure 2, imaging at different wavelengths results in significantly different imaging contrast values. We measured the Weber contrasts of lymph nodes against surrounding tissue at 1000 nm, 1175 nm, 1250 nm, 1400 nm, and 1550 nm (see Supplementary Information). The corresponding Weber contrasts are 0.24, 0.40, 0.36, 1.19, and 1.68, respectively. Contrast values obtained near the water absorption peak around 1450 nm are clearly superior to contrast values obtained at shorter wavelengths.

**Figure 2.**
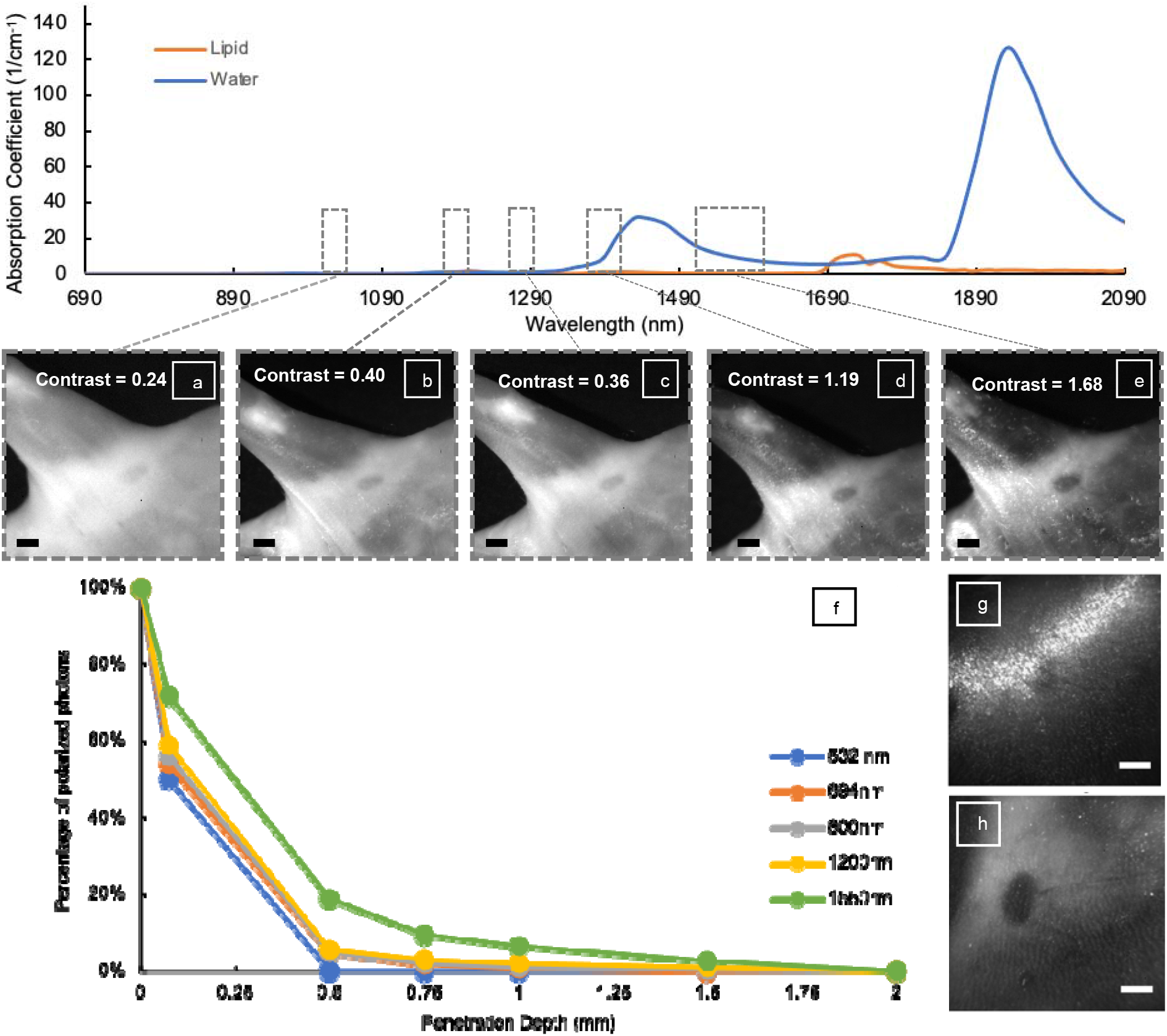
The SWIR-OPI images of mouse inguinal lymph nodes at (a) 1000 nm (b) 1175 nm (c) 1250 nm (d) 1400 nm (e) 1550 nm. The contrasts are calculated as Weber Contrast. Panel (f) shows the Monte Carlo simulation results of how linearly polarized light with different wavelengths becomes unpolarized after interacting with biological tissue at different depths. The longer the wavelength, the deeper the linearly polarized light penetrates. This corroborates that OPI is particularly effective in the SWIR range. At 1550 nm, imaging contrast with OPI, shown in panel (h), is much improved compared to that of without OPI, shown in panel (g). All scale bars are 2 mm.

As for the source of the imaging contrast, water content is definitely a major contributor. Tissue that is high in water content is generally dark under SWIR-OPI, such as lymph nodes, muscle tissue, and mammary ducts, which are shown in Figure 2(d), Figure S18 and Figure S19, respectively. Tissue that is low in water content is generally light or bright under SWIR-OPI, such as fat, shown in Figure 1(c). Interestingly, there are some special cases that demonstrate that the source of contrast is more nuanced than simply considering water content. For example, blood vessels do not appear to be nearly as dark as lymphatic vessels under SWIR-OPI, despite containing blood that is clearly high in water content, as shown in Figure S19(B). The low contrast of blood vessels can be attributed to the strong scattering effects from blood cells, which occupy up to 40% of blood volume. The absorption and scattering effects are competing factors when it comes to imaging blood under SWIR-OPI: water absorption results in low photon counts in the imaged area, and strong scattering effects from blood cells result in high photon counts in the imaged area. A detailed analysis of blood visualization under SWIR-OPI was performed in the Supplementary Information. This unique observation potentially makes SWIR-OPI even more attractive as a lymphatic imaging technique as it provides orthogonal information to blood flow imaging techniques that are well-established. Other interesting findings using SWIR-OPI include high contrast from mammary ducts and skeletal muscle tissue (see Supplementary Information). As a label-free imaging technology based on the natural chemical composition contrast in biological tissues, SWIR-OPI can be potentially used for a wide range of biomedical imaging applications yet to be explored.

The second key part of the imaging system is the orthogonal polarization setup. Although it is a commonly used strategy to reduce surface glare in scientific and medical imaging as well as everyday macrophotography, the benefit is significantly enhanced when imaging in the SWIR range. As shown in Figure 2(g) and (h), even when at the optimal imaging wavelength, the lymph node is not clearly visible without orthogonal polarization, whereas the lymph node can be clearly seen with orthogonal polarization. To demonstrate this effect quantitatively, we performed Monte Carlo simulation to calculate the penetration depths in tissue at different wavelengths. 100% linearly polarized light with wavelengths of 532 nm, 690 nm, 800 nm, 1200 nm, and 1550 nm interacts with biological tissues with known optical properties. With tissue sample thicknesses of 0.075 nm, 0.5 mm, 0.75 nm, 1 mm, 1.5 mm, and 2 mm, we calculated the percentage of photons detected that are aligned with the original direction of polarization. As shown in Figure 2(f) and Table 1 in the Method section, light at shorter wavelengths loses its polarization at much shallower penetration depths than light at longer wavelengths. At a typical near-infrared wavelength of 800 nm, only 2.0% of detected light is linearly polarized after interacting with a 0.75-mm thick specimen of biological tissue, whereas at the SWIR wavelength used here, 1550 nm, 2.7% of detected light is linearly polarized after interacting with a twice-as-thick 1.5-mm tissue specimen. This calculation corroborates the well-known knowledge that SWIR light scatters much less than near-infrared light, which makes SWIR particularly suitable for deep tissue imaging.

**Table 1.** Percentage of light with the same polarization as the illumination light at different tissue depths.

| Wavelength<br>(nm) | Tissue depth (mm) |  |  |  |  |  |
| --- | --- | --- | --- | --- | --- | --- |
|  | 0.075 | 0.5 | 0.75 | 1 | 1.5 | 2 |
| **532 nm** | 50.00% | 0.00% | 0.00% | 0.00% | 0.00% | 0.00% |
| **694 nm** | 54.00% | 5.00% | 1.00% | 0.75% | 0.00% | 0.00% |
| **800 nm** | 56.00% | 5.00% | 2.00% | 1.00% | 0.20% | 0.10% |
| **1200 nm** | 59.00% | 5.40% | 3.00% | 1.80% | 1.00% | 0.10% |
| **1550 nm** | 72.00% | 19.00% | 9.50% | 6.40% | 2.70% | 0.15% |

The third key part of the imaging system is the home-built object-space telecentric SWIR lens. Telecentric lenses are commonly used in machine vision where the sizes of imaged objects need to be measured precisely.(*43*) Images collected by such lenses have magnifications that are independent of an object’s distance from the imaging lens. This feature becomes extremely useful when we collect intravital measurements over time to monitor lymph node size longitudinally within the same animal. For example, the specific position, angle, and posture of an imaged animal could vary slightly from day to day, which would change the distance between the imaging lens and the imaged animal. An object-space telecentric imaging lens can prevent such slight variations from producing significant measurement errors in the size of features of interest. However, because SWIR imaging for biomedical applications is still an emerging field, there are no commercially available telecentric lenses to the best of our knowledge, and other custom-built SWIR imaging setups reported by peer researchers(*44*) are not telecentric. We built a telecentric lens by putting an iris (SM1D12C, aperture size: 5 mm, Thorlabs) at the focal plane of a commercially available SWIR lens (50 mm focal length, Kowa American Corporation, LM50HC-SW), and an extension tube (0.3 inch, Thorlabs) was attached behind the iris. The size of the pinhole was optimized, as the telecentricity improves with smaller pinhole sizes, but the imaging resolution worsens (see Supplementary Information).

### Label-free Lymph Node Mapping with SWIR-OPI

Indocyanine green (ICG) has been demonstrated to be a useful tool for lymph node mapping in animal experiments as well as in the clinical setting.(*21, 45, 46*) We used ICG injection as a benchmark for SWIR-OPI. (808 nm illumination, 900 nm long-pass detection, irradiance 20 mW/cm^2^) In Figure 3(a) and (c), ICG was used to visualize inguinal lymph nodes of BALB/c mice and Wistar IGS rats, respectively. The lymph nodes can be clearly seen; however, liver uptake of ICG is a source of interference in the imaging results, as shown in Figure 3(a). It is noteworthy that mouse skin is approximately 0.7 mm in thickness, rat skin is approximately 2 mm thick, and human skin ranges from 0.5 to 5 mm in thickness.(*47*) Being able to visualize lymph nodes noninvasively through intact skin in both mice and rats proves that SWIR-OPI is capable of visualizing features at more than 2 mm depth, which is consistent with the Monte Carlo simulation results mentioned above.

**Figure 3.**
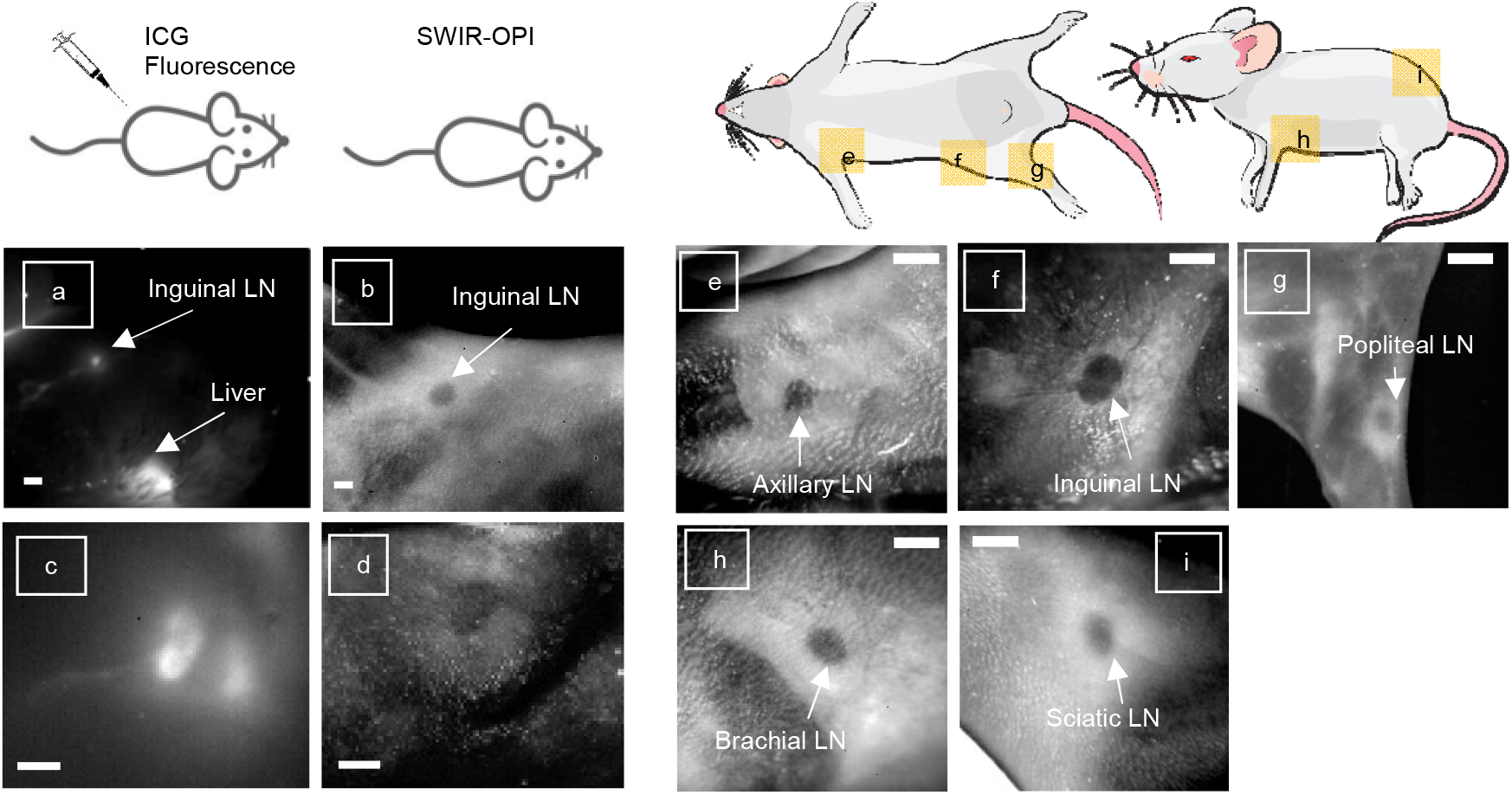
(a) and (c) show the fluorescence images of inguinal lymph nodes of BALB/c mouse (a) and rat (c) with ICG injections. (b) and (d) demonstrate the corresponding SWIR-OPI images respectively. (e) – (f): SWIR-OPI lymph node mapping of (e) axillary lymph node; (f) inguinal lymph node; (g) popliteal lymph node; (h) brachial lymph node; (i) sciatic lymph node; All scale bars are 2 mm.

Besides imaging the inguinal and mesenteric lymph nodes mentioned above, SWIR-OPI can also be used to map almost all of the superficial lymph nodes in mice noninvasively. As shown in Figure 3(e) and (g) – (i), the axillary, popliteal, brachial, and sciatic lymph nodes, respectively, were visualized using the same imaging setup.

### Automated Lymph Node Size Measurements Empowered by Deep Learning

In addition to visualizing lymph nodes, with the help of a home-built telecentric SWIR lens, we use SWIR-OPI as a monitoring tool to track daily changes in lymph node size quantitatively. A series of images is taken by recording a video of at least 300 frames with a 0.01 second exposure time. The size of lymph nodes can be extracted by manually tracing the perimeter of lymph nodes and filling the shape with a solid color using commercially available computer graphics tools, such as Microsoft Paint or Adobe Photoshop. The number of pixels that a lymph node occupies is a proxy for the size of a lymph node. Similar practices are routinely performed clinically for ultrasound, CT, or MRI images of lymph nodes of cancer patients.(*48*) Over the course of our studies, we accumulating over 500,000 images – making manually tracing and processing these images was practically impossible. Therefore, we decide to automate the image tracing process using a deep learning-based model.

We utilized U-Net as our main segmentation network for automatic lymph node segmentation of raw image data. U-Net has been widely used in many biomedical image segmentation problems and has been proven to be quite successful.(*49*) Following the original U-Net design, we further increase the level of encoding to 6 levels. This level of encoding enables the network to analyze a larger field of view than the original design, allowing for handling of larger objects per image. Following modern deep learning convention, batch normalization layers are added after each convolutional layer. More in-depth description of the U-Net model in this work is demonstrated in Supplementary Materials. We manually labeled 364 images for network training; each image contains one lymph node. Adam optimizer(*50*) is used for model training, where the learning rate is set at 5e^-4^ for the first 30000 iterations and decays to 5e^-5^ for the remaining 20000 iterations. During model testing, the trained network is directly applied on top of an image and produces a probability map with the same size as the input image. The final binary map (for lymph node detection) is generated by applying an argmax operation along the channel dimension of the probability map.

### Monitoring Lymph Node Responses to Cancer with SWIR-OPI

Using the aforementioned imaging hardware combined with U-Net for rapid image processing, we monitored lymph node sizes daily in BALB/c mice after the orthoptic implantation of 4T1 triple-negative breast cancer cells (100,000 cells in 30 μL of PBS) in the 4^th^ mammary fatpad on the right side of the body. Mice in the control group were injected with 30 μL of PBS. For 14 consecutive days, we recorded daily images of the inguinal lymph node draining the site of injection and the contralateral lymph nodes as a control. On the 14^th^ day, primary tumors were resected from tumor-bearing mice (study group) in order to reduce disease burden, prevent direct local tumor invasion of the draining lymph node, and allow for lymph node metastasis to develop after resection. After surgeries, mice were kept alive for another two weeks before sacrifice for histology studies. Imaging with SWIR-OPI of the lymph nodes became infeasible after the surgeries, because lymph nodes were obstructed by sutures and post-surgical scar tissue. For this reason, no images after Day 14 were included in this study. On Day 30, mice were sacrificed, and inguinal TDLNs and contralateral lymph nodes were collected for histology studies. H&E histology slides were analyzed for the presence of metastasis in lymph nodes.

The raw recorded SWIR-OPI images were processed by U-Net to measure lymph node size. After examining the histology results, mice with metastasis in their TDLNs were grouped to into one cohort, and plotted in Figure 5 in red. Mice in the study group without lymph node metastasis were grouped into another cohort and plotted in Figure 5 in blue. Figure 4 shows that metastatic TDLNs increased in size by as much as 130%, whereas non-metastatic TDLNs grew in size by at most 65%. Also, differences in size between metastatic lymph nodes and non-metastatic lymph nodes can be observed as early as Day 4. This observation is consistent with previous reports of pre-metastatic lymph node expansion in studies of 4T1 cells in mouse models.(*51, 52*) The cohort with metastatic TDLNs has 7 subjects, the cohort with non-metastatic TDLNs has 12 subjects, and the control group has 2 subjects. The downward trend in lymph node sizes around Day 11 – 13 was largely attributed to a measurement artifact due to the large size of primary tumors and their close proximity to the TDLNs. More details about this artifact are discussed in the Supplementary Information.

**Figure 4.**
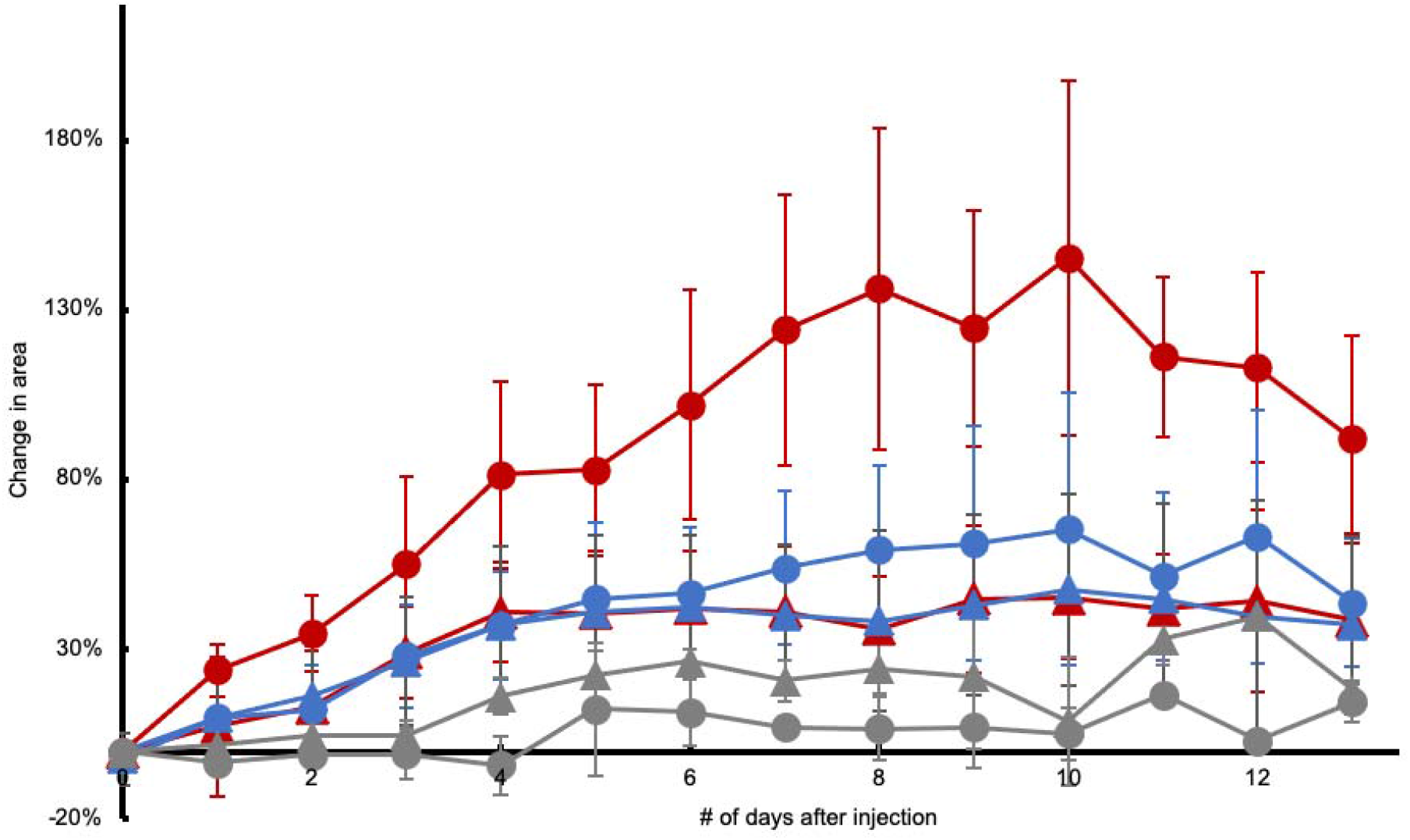
Red symbols correspond to mice injected with 4T1 cells with metastatic lymph nodes. Circles represent measurements on tumor-draining lymph nodes, and triangles represent measurements on contralateral lymph nodes. Blue symbols correspond to mice injected with 4T1 cells with non-metastatic lymph nodes. Gray symbols correspond to mice injected with PBS. All injections are done on the right side.

**Figure 5.**
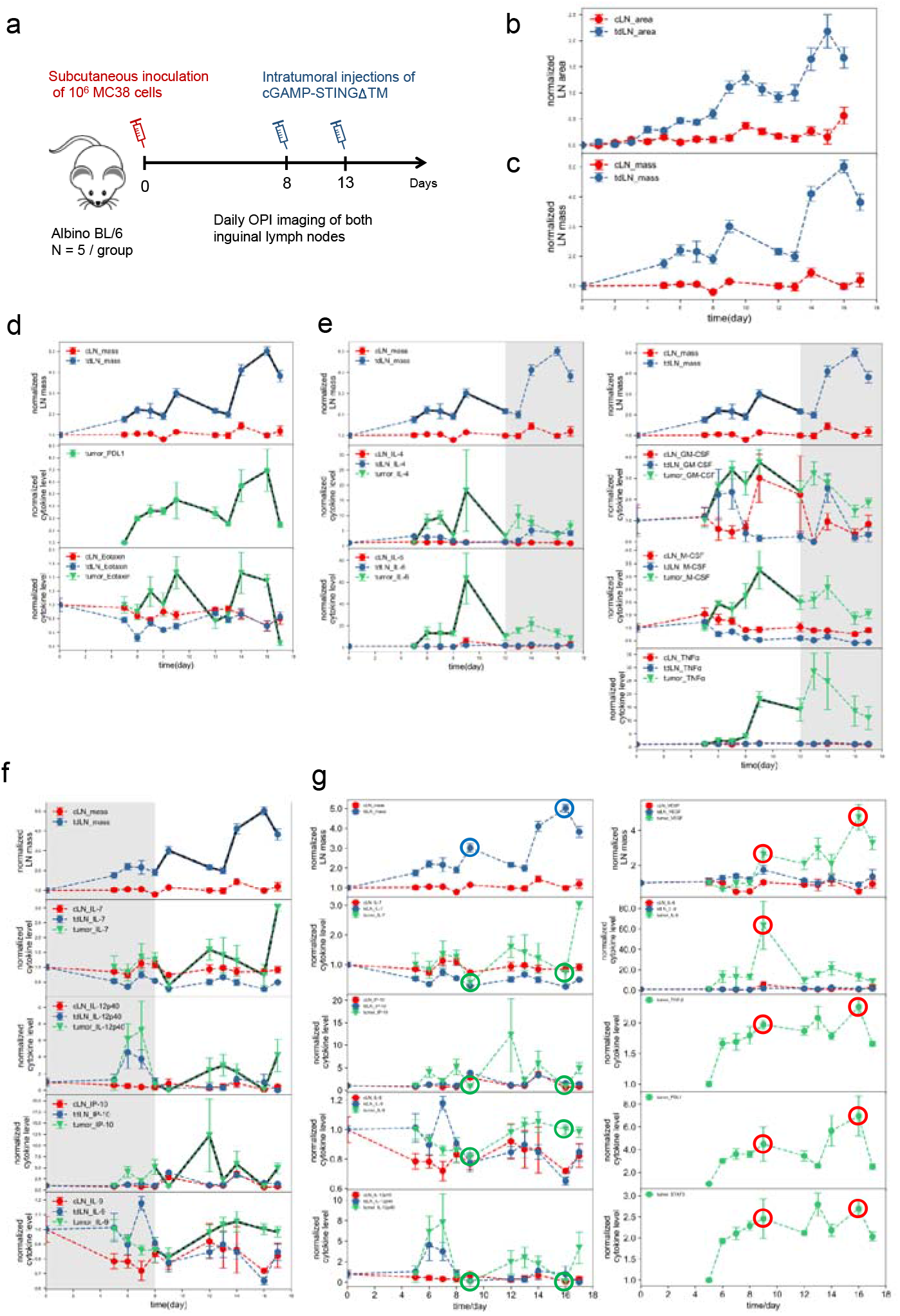
Correlation of tdLN sizes to cancer biomarkers in a subcutaneous MC38 model. (a) Groups of Albino BL/6 mice were inoculated with one million MC38 cells subcutaneous, followed by two intratumoral treatments at day 8 and 13 post inoculation. Area of both inguinal lymph nodes were measured via OPI imaging daily throughout the course of treatment (b) and compared with weighed mass of lymph nodes collected from sacrificed mice at corresponding time points (c). (d) Comparison of tdLN mass change with levels of PD-L1 and Eotaxin in the tumor. (e) Levels of IL-4, IL-6, GM-CSF, M-CSF, and TNFα in the tumor showed similar trend with tdLN mass change at the initial growth stage of tumor plus the early stage of treatment (first 12 days). (f) Levels of IL-7, IL-12p40, IP-10, and IL-9 in the tumor showed inverse trend with the tdLN mass change during the treatment (day 8-17). (g)Tumor microenvironment appears most immunosuppressive and pro-tumor when tdLN mass peaks (day 9 and 16 as marked by blue circles), at these two time points, anti-tumor cytokines are at lowest level while pro-tumor ones are highest. Anti-tumor biomarkers are circled in green while pro-tumor ones are circled in red. Values are plotted as mean ± SEM.

### Correlating tdLN Sizes to Relevant Biomarkers of Immuno-Oncology

We next sought to develop this technique as a diagnostic tool for monitoring the tumor microenvironment by correlating the variation in tdLN size with changes in cancer biomarkers over time. For this study, we chose a tumor model with lower metastatic potential – subcutaneously inoculated colon cancer MC38 – to reduce the contribution of tdLN enlargement from harboring metastatic tumor tissues. Groups of albino BL/6 Mice (N=5) were inoculated subcutaneously with 1 million MC38 cells in the right hind flank. At day 8 and 13 post-inoculation, we intratumorally injected the tumors with STINGΔTM + cGAMP, a therapeutic combination that has been shown to boost anti-cancer immunity(*53*). Sizes of both inguinal lymph nodes were measured daily to monitor their change in response to tumor growth and treatments [Figure 5 (a)-(c)]. Throughout the course of treatment, we sacrificed 11 groups of mice at each representative time point of tdLN size changes and harvested their tumors and inguinal lymph nodes to quantify the levels of 38 cancer relevant cytokines, chemokines, growth factors, and other biomarkers. We also recorded the mass of lymph nodes and found them in good agreement with the trend of lymph node area change measured via OPI imaging.

We then compared the trend of lymph node mass change with the biomarker levels and made note of two key observations. Firstly, the levels of many important biomarkers in the tumor microenvironment exhibited a strong correlation with tdLN size, such as those of PD-L1 and eotaxin. Other markers such as IL-4, IL-6, M-CSF, and TNF-alpha had good agreement with tdLN size during the first 12 days post-tumor inoculation, while IL-7, IL-12p40, and IP-10 followed an inverse trend during day 6-18 post-tumor inoculation [Figure 5 (d)-(f)]. Additionally, we found that the tumor microenvironment appeared to be the most immunosuppressive and pro-tumorigenic at the local maxima of tdLN size - day 9 and day 16. At these timepoints, many anti-tumoral biomarkers such as IL-7, IL-9, IL-12p40, and IP-10 reached a local minimum, whereas pro-tumoral factors such as VEGF, TGF-beta, PD-L1, IL-6 and STAT3 reached a local maximum [Figure 5(g)]. To explore the translational potential of this technique, we next evaluated its efficacy in human clinical samples.

### Human Pilot Study of SWIR-OPI in Human Mesentery Specimens

In anatomical pathology, identifying and extracting lymph nodes is a critically important process, because lymph node count has been shown to be correlated with cancer patients’ survival rate.(*54, 55*) However, this process is labor-intensive and time-consuming, and it leads to large differences in inter- and intra-observer variability.(*56-58*) SWIR-OPI can potentially solve this problem by visually assisting pathologists and Pathologists’ Assistants in identifying lymph nodes in *ex vivo* anatomical specimens.

SWIR-OPI is compatible with *in vivo, ex vivo*, and also formalin-fixed tissues. Human mesenteric lymph nodes shown in Figure 6 are fixed by formalin. As shown in Figure 6(a) and (c), human mesenteric lymph nodes are buried in fat tissue that is extremely similar in color to lymph nodes to the naked eye. With SWIR-OPI, shown in Figure 6(b) and (d) and similar to the *in vivo* data shown in Figure 1, human mesenteric lymph nodes are visible with high contrast. Additionally, this experiment was performed with an affordable, commercially available uncooled InGaAs camera (SU320M-1.7RT, Sensors Unlimited). Translation of this technology can provide a powerful imaging guidance tool to assist pathologists and pathologist’s assistants with extracting lymph nodes from anatomical pathology specimens. Since this technology has been demonstrated to work with *in vivo*, freshly excised, and formalin-fixed tissues, it is also promising as an intra-operative imaging guidance tool for surgeons to find more lymph nodes faster, especially for thoracic, gynecological, and urological oncology surgeons for whom sentinel lymph node procedures could be difficult.

**Figure 6.**
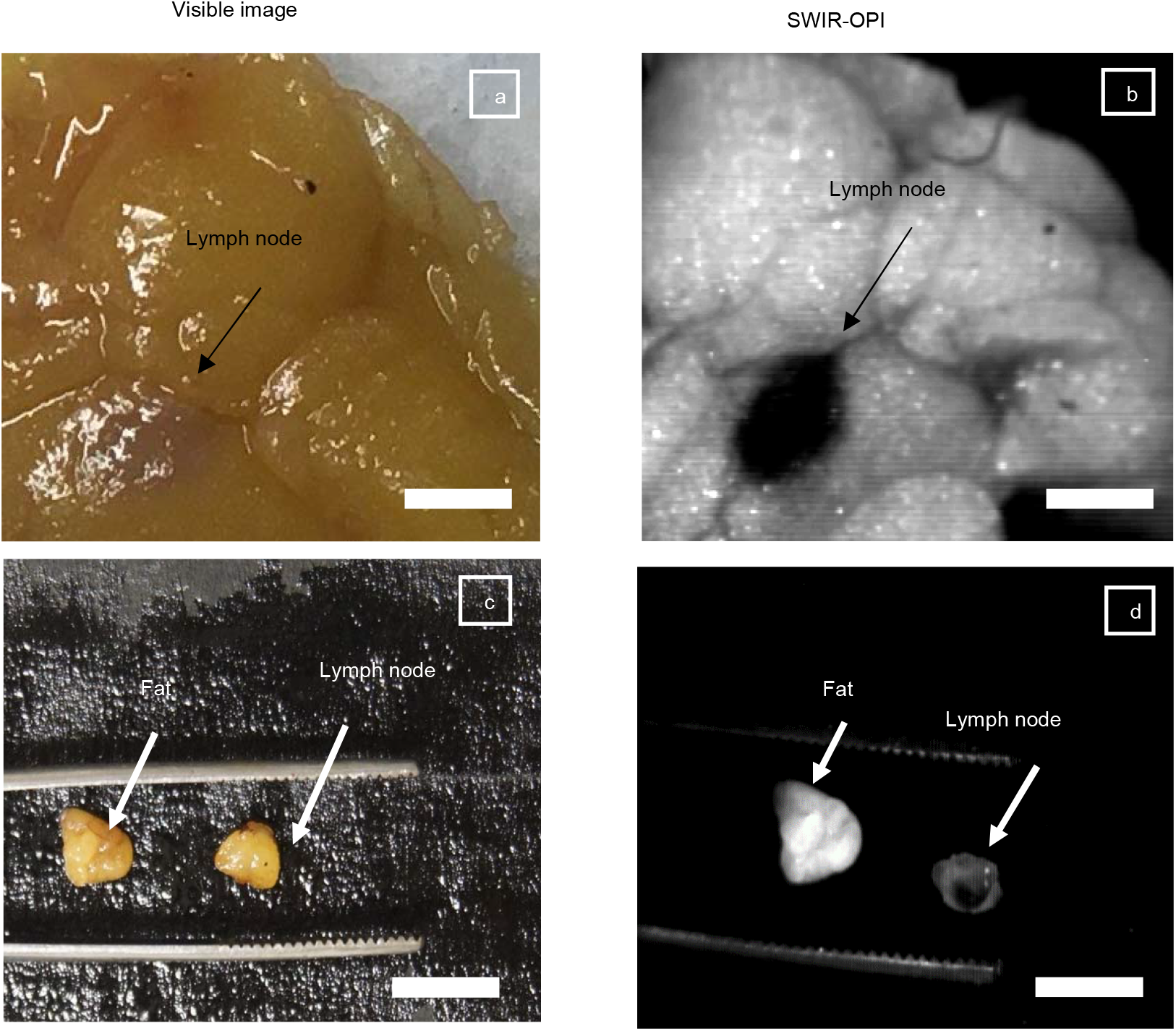
Panel (a) and (c) depict regular visible light images of human mesenteric lymph nodes surrounded by mesenteric fat. Lymph nodes are hard to see. (b) and (d) are images taken on the same mesentery tissue with SWIR-OPI. Lymph nodes are clearly visible. Scale bars are 5 mm.

## DISCUSSION

Figure 4 provides some interesting biological insights. We observed lymph node expansion as early as Day 4 after tumor inoculation, and this result is consistent with previous literature reports on the 4T1 cell line.(*51, 52*) Detmar and colleagues reported enlargement of the lymphatic vascular network in TDLNs that precedes lymph node metastasis in a mouse model inoculated with 4T1 breast cancer cells. Lymphatic expansion in TDLNs is mediated by sprouting and proliferation of lymphatic endothelial cells (LECs) as early as 4 days after tumor implantation.(*51*) Cao and colleagues reported that 4T1 primary tumors induced B cell accumulation in draining lymph nodes *in vivo*.(*52*) These B cells selectively promoted lymph node metastasis by producing pathogenic IgG that targeted glycosylated membrane protein HSPA4, and activated both the HSPA4-binding protein ITGB5 and the downstream Src/NF-κB pathway in tumor cells for CXCR4/SDF1α-axis-mediated metastasis. High HSPA4 expression was shown to be correlated with a high degree of lymph node metastasis. They also observed significant lymph node expansion within one week, while the onset of metastasis to the lymph node(s) did not occur until two weeks after inoculation.(*52*)

While the findings of previous researchers could explain the biological mechanism of our results, our results actually provide additional information about pre-metastatic lymph nodes. We identified a quantitative relationship between the size of pre-metastatic lymph nodes and the likelihood of eventual metastasis. The lymph nodes that grow larger in the early days following cancer cell injections are more likely to develop metastasis later on. As shown in Figure 4, we observed that TDLNs of mice in the metastasis group became significantly larger than the contralateral lymph nodes as early as Day 4, which is consistent with previous reports. A detailed ANOVA analysis of the animal data is provided in the Supplementary Information. Notably, within the same timeframe, the TDLNs that showed metastasis upon histological examination were significantly larger than the non-metastatic TDLNs. This study showed that SWIR-OPI can be used as a convenient and powerful tool to monitor changes to the lymph nodes in response to disease progression. SWIR-OPI can also be potentially used to monitor and investigate the effectiveness of treatment intervention *in vivo* with respect to lymph node responses to different treatment regimens over the course of disease progression.

Monitoring cancer biomarkers in real time have important clinical values in predicting treatment outcome and monitoring disease stages(*59, 60*). Most notably, PD-L1 expression in tumors is one of the best predictive markers for patient outcome after treatment with anti-PD-(L)1 immune checkpoint blockade(*61*). Clinical trials of non-small-cell lung cancer (NSCLC) and melanoma treated with anti-PD-1 have reported that patients with high PD-L1 expression (proportion score > 50% and 33%, respectively) exhibit a response rate of ∼50%, whereas patients with low PD-L1 expression (<1% for both cancers) showed response rates below 10%(*62, 63*).

Precisely identifying immunosuppressive time points during cancer progression can inform the design of dosing schedules for combination immunotherapies that engage various facets of the immune response (for instance clinical trial NCT02983045 anti-PD1/CTLA4 and IL-2). At present, combination reagents are most often dosed simultaneously or according to a schedule determined by FDA approved usage of the individual components(*64*). The diagnostic imaging proposed herein may provide important insight in optimizing the timing of cancer immunotherapy interventions, potentially enhancing treatment efficacy.

Moreover, the safe and non-invasive prediction of tumor biomarkers has important clinical value for routine cancer screening, cancer disease staging, and monitoring the mechanism-of-action of anti-cancer therapeutics during treatment. Existing invasive approaches such as tumor biopsies have been reported to increase the risk of cancer metastasis, whereas low-invasive techniques such as serum and urine sampling are less correlated to the tumor microenvironment and are hence limited by the amount of diagnostic information and accuracy(*65, 66*). Our OPI imaging technique of the lymph nodes thus provides an ideal approach to monitor the tumor microenvironment in real-time, providing valuable information for the design of more effective, personalized treatment strategies. Beyond clinical applications, these results also expand our fundamental knowledge regarding the longitudinal role of tdLNs in an evolving immunosuppressive tumor microenvironment - a need that has been highlighted in recent literature as most existing studies on tdLN-tumor crosstalk have been restricted to single timepoints(*67*).

One might argue that ultrasound, CT, and MRI could achieve similar functionality and provide similar insights as SWIR-OPI if daily monitoring were performed. However, the likely reason why such studies have not been done before is that these existing technologies are far more cumbersome than SWIR-OPI. Compared to ultrasound, CT, and MRI, SWIR-OPI is contactless and convenient to operate. Additionally, SWIR-OPI also offers promise in an area that is completely incompatible with ultrasound, CT, and MRI – lymphatic reconstruction surgeries.(*68-70*)CT, MRI, and ultrasound cannot visualize lymphatic vessels, whereas SWIR-OPI can visualize lymphatic vessels at high resolution and with high contrast, as shown in Figure 1.

SWIR-OPI offers exciting promises in clinical surgery procedures. The current standard of care for lymphatic reconstruction surgeries relies on ICG fluorescence imaging, which has drawbacks of increased cost, inconvenience, and time pressure. Compared to dye -based lymphatic imaging procedures, which rely on injections of dye molecules or radioactive tracers that can alter the native physiology, SWIR-OPI shows natural unperturbed lymph nodes and lymphatic vessels. Similarly, the standard of care for identifying tumor draining lymph nodes, i.e. sentinel lymph nodes, also relies on injections of dye or radioactive tracers. The foundational hypothesis of the sentinel lymph node procedure is that dye or radioactive tracers drain following the same lymphatic routes as metastatic cancer cells. Though demonstrated widely to be an successful clinical procedure, sentinel lymph node detection can be confounded when metastatic cancer cells in lymph nodes and lymphatic vessels block lymphatic drainage from the primary tumor site to the “true” sentinel lymph node(s) and cause rerouting of lymph to different lymph nodes.(*71, 72*) Injected markers thus can direct surgeons to a “false” sentinel lymph node, causing the “true” metastatic sentinel lymph node to be missed. This procedure carries the risk of producing false negative results in staging reports for cancer patients.(*71*) SWIR-OPI can potentially complement the sentinel lymph node procedure by overcoming this shortcoming, as SWIR-OPI visualizes the lymphatic vessels and lymph nodes independent of their ability to drain injected markers. Also, as a label-free technique, SWIR-OPI would add no additional risk to the sentinel lymph node procedure, as it avoids allergic or immunogenic reactions to dye molecules,(*73, 74*) prolonged skin staining,(*75*) or exposure to radiation.(*76*) As the field of surgical oncology is moving towards less aggressive, more accurate lymph node dissection procedures in order to improve the quality of life for cancer patients,(*77, 78*) SWIR-OPI offers great promise in becoming part of the standard of care for future oncology surgeries as a complement to current sentinel lymph node procedures.

## CONCLUSION

We developed a new label-free optical imaging method that visualizes the lymphatic system in mice, rats, and human mesentery specimens. It utilizes shortwave infrared (SWIR) imaging with an orthogonal polarization imaging (OPI) configuration. We integrated U-Net deep learning algorithms to automatically segment out lymph nodes in SWIR-OPI images and measure the sizes of the lymph nodes longitudinally in a mouse model of triple-negative breast cancer. With this deep learning enhanced imaging method, we identified pre-metastatic lymph node expansion as early as four days after tumor inoculation. We also discovered that, as early as Day 4, lymph nodes that will be metastatic on Day 30 have already grown significantly larger than the other TDLNs that will not develop metastasis by Day 30. In addition, the tdLN sizes were found to closely correlate with many important biomarkers in the tumor such as PD-L1, based on which we could monitor in real-time how the tumor microenvironment evolves and provide valuable insight for dosing schedule design. We also demonstrated that SWIR-OPI can also be used to help identify human lymph nodes in clinical procedures, as human lymph nodes can also be visualized with high contrast in real time without any injection or radiation. SWIR-OPI can be a powerful tool for animal research as well as clinical imaging, especially anatomical pathology and surgical oncology.

## MATERIALS AND METHODS

### Study design

This study is to develop a non-invasive lymphatic system imaging technique and apply it for monitoring cancer disease stage in real-time hence informing the dosing schedule design of immunotherapy. All animal studies were performed according to approved protocols by MIT Division of Comparative Medicine. Human samples were obtained with informed consent. Sample sizes were determined from pilot studies and previous work, not from power calculations. Same purchase cohort of female mice were used for each experiment and allocated into groups randomly. Lymph node imaging were performed by a blinded researcher while tumor treatments were administrated by a researcher not blinded.

### Animal study preparation

Animal Care and Use Committee protocols. Animal experiment procedures were preapproved (Protocol #1215–112–18) by the Division of Comparative Medicine and the Committee on Animal Care, Massachusetts Institute of Technology, and in compliance with the Principles of Laboratory Animal Care of the National Institutes of Health, USA.

Hair removal is critical to the success of shortwave infrared orthogonal polarization imaging (SWIR-OPI). Compared to skin and other soft tissues, hair is extremely low in water content. Since water content contributes significantly to the imaging contrast, as discussed in the Supplementary Information, hair appears extremely bright under shortwave infrared (SWIR) illumination compared to the skin tissue around it. If hair is not completely removed, it not only will obstruct the view of the lymph nodes by covering them, but also will overshadow the imaging contrast between the lymph nodes and the surrounding tissue. The hair removal procedure is performed as follows:

To remove mouse hair for the first time, 10-week-old BALB/c mice (BALB/cAnNCrl, Charles River Laboratories) are first administered 1.5% isoflurane for anesthesia. To remove hair cleanly around the inguinal lymph node regions, mouse hair is first shaved with an electric razor. After shaving the hair, a hair removal product (Nair™, Church & Dwight Co.) is applied thoroughly to the entire lower body of the mouse and is left on for approximately 1 minute.

Nair™ is then wiped off cleanly with baby wipes and the mouse skin is rinsed with warm water. The mouse is then dried with paper towels and placed back in its cage. Heating pads are placed underneath the cages in order to keep the body temperature of mice stable. Note that this hair removal process is only compatible with mice with full-grown hair — repeating this process within less than 2 weeks causes severe chemical burns to mouse skin, which can be fatal or at least damage the skin significantly. Chemical burns caused by Nair™ can also cause substantial scarring, which will obstruct the view of the lymph nodes.

In order to perform daily imaging sessions on the same mice, mouse skin has to be kept clean of hair. When left without maintenance, the depilated mice can grow a significant amount of hair within 24 hours, preventing daily *in vivo* monitoring of lymph nodes. In order to prevent new hair from growing, a hair growth inhibitor product (Hair Growth Inhibitor, Completely Bare) is applied to the bare mouse skin twice a day. By performing this hair maintenance routine, mouse skin can be kept bare for the imaging period of two weeks without repeating any hair removal procedures.

### 4T1 cancer study

10-wk-old female BALB/c mice (Charles Rivers: BALB/cAnNCrl) are first depilated and stapled with ear tags for tracking purposes. Images are saved and named after the ear tag numbers and saved in the folder Day 0. Images in the Day 0 folder represent the condition of the lymph nodes in their unperturbed conditions before any injection. It is noteworthy to mention that ear tags frequently fall off over time as mouse ears heal from stapling. Daily monitoring of the ear tags is necessary to ensure successful tracking of individual mice over the course of long-term imaging experiments.

The experimental group of mice is injected orthotopically with 100,000 4T1 cancer cells suspended in 30 μL PBS solution into the fourth nipple on the right side. The control group of mice is injected with 30 μL PBS solution at the same location. In this study, there were 19 mice in the experimental group and 2 mice in the control group. After injections, mice are treated with the hair growth inhibition products mentioned above.

At the 24-hour time point, all mice are imaged again, and the images are saved in the Day 1 folder. The same process is repeated again on a daily basis from Day 2 to Day 13. The sizes of tumors are also recorded for reference.

Because we implanted the tumor cells orthotopically, the primary tumor site is close to the tumor-draining lymph nodes. On Day 13, the sizes of tumors are so large (3 cm x 3 cm) that the primary tumors are typically extremely close to the tumor draining lymph nodes. It is impossible to image lymph nodes consistently after Day 13 without tumor resection due to direct tumor infiltration of the lymph nodes, and it is also likely inhumane given the heavy tumor burden.

A primary tumor excision surgery is performed on Day 13 to resect the primary tumor. The specific procedure has been well-reported.(*8*) Mice are kept alive for another 14 days after the primary tumor resection surgery before sacrifice. Daily imaging sessions were not performed after the surgery, because the sutures and scars were extremely close to the lymph nodes. Sutures and scars obstruct the view of lymph nodes, and the presence of sutures and scars makes hair removal or hair growth inhibition methods impossible to perform. Hair growth near the suture areas makes *in vivo* SWIR-OPI of the lymph nodes impossible.

It is well-understood that for orthotopic 4T1 implantation, metastasis does not occur during the first two weeks after implantation.(*52*) It is also well reported that under this 4T1 orthotopic implantation protocol, only 40% of injected mice would develop lymph node metastasis.(*79*) Since we can consistently measure lymph node sizes for all mice while it is known that only some mice will develop metastasis, orthotopic injection of 4T1 tumor cells is a suitable model for investigating how changes in lymph node size correlate to the prediction, onset, and prevalence of metastasis.

### Inoculation and intratumoral injections of MC38 tumors

Groups of Albino BL/6 mice were subcutaneously injected with 10^6^ MC38 cells in 100 μL sterile PBS in their right hind flank. At day 8 and day 13 post inoculation, the mice were intratumorally injected with 40 μg of STINGΔTM protein mixed with 1 μg of 2’3’-cGAMP (InvivoGen, Cat # tlrl-nacga23-02) in 25 μL sterile PBS. STINGΔTM protein was produced in house from *E. coli* expression according to the literature.(*80*)

### Analysis of tumor and lymph node biomarkers

At each representative time points, a group of 5 mice were sacrificed and with their tumors and both inguinal lymph nodes collected and frozen in −80 C. At the end of the study, all frozen tumors and lymph node samples were processed for biomarker analysis via Luminex and qPCR.

For Luminex analysis, around 50 mg tumors and whole lymph nodes were ground with tube pestles in 100 μL ice-cold T-PER tissue protein extraction buffer (ThermoFisher, Cat # 78510) with HALT protease and phosphatase inhibitor (ThermoFisher, Cat # 78442). The lysate was then incubated for 30 min at 4 C under gentle vortex, followed by centrifugation. The total protein concentration in the isolated supernatant was quantified by Nanodrop spectrophotometer with absorption at 280 nm. Finally, the samples were all diluted to a total protein concentration of 5 mg/mL and shipped to Eve Technology for Luminex analysis. The full list of biomarkers measured by Luminex is: Eotaxin, G-CSF, GM-CSF, IFNγ, IL-1α, IL-1β, IL-2, IL-3, IL-4, IL-5, IL-6, IL-7, IL-9, IL-10, IL-12p40, IL-12p70, IL-13, IL-15, IL-17A, IP-10, KC, LIF, LIX, MCP-1, M-CSF, MIG, MIP-1α, MIP-1β, MIP-2, RANTES, TNFα, VEGF-A For qPCR analysis, only tumors were analyzed as there weren’t enough lymph node tissues after Luminex. Total RNA was extracted from 100 mg tumor tissue using the Direct-zol RNA Miniprep kits (Zymo Research, Cat # R2053) then reverse transcribed with cDNA Reverse Transcription Kit (ThermoFisher, Cat # 4374966), followed by amplification with SYBR Green PCR Master Mix (ThermoFisher, Cat # 4309155) with mouse actin as housekeeping gene. qPCR primers used is listed below, and amplification products were confirmed to have the expected length with agarose gel electrophoresis:

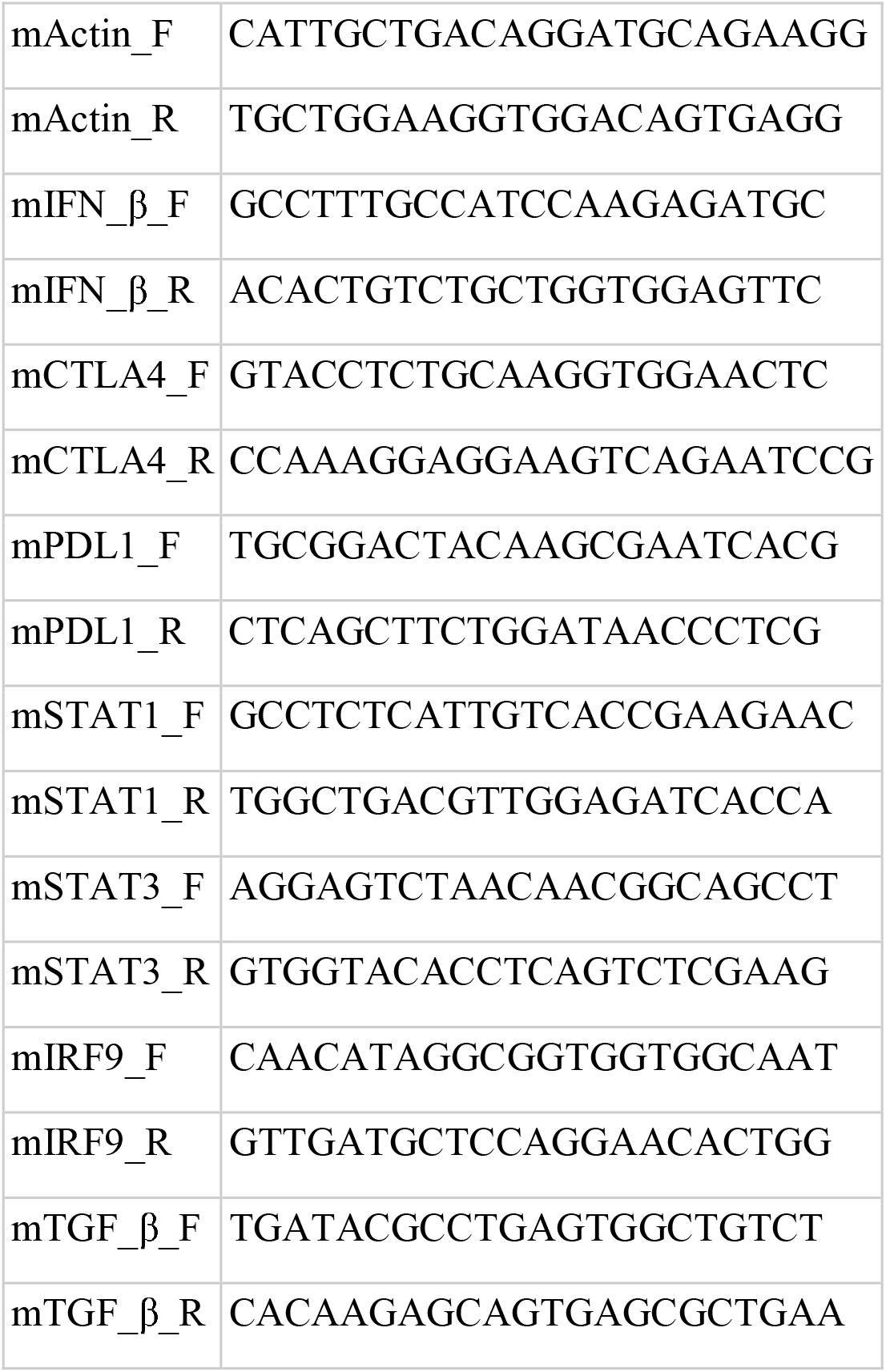

### Human specimen pilot study

This human pilot study was carried out at the School of Health Sciences of Quinnipiac University. The study protocol was reviewed by the Institutional Review Boards (IRB) of the Quinnipiac University and was determined to be IRB exempt and compliant with all Health Insurance Portability and Accountability Act (HIPAA) requirements.

Colectomy specimens are procured by American Society for Clinical Pathology (ASCP) certified Pathologist’s Assistants. Prior to experiments, similar to clinical standard-of-care, colectomy specimens are fixed in formalin solution for at least 2 hours. Mesenteric fat tissue is dissected away from the colectomy specimens, and subsequently cut into sliced at around 2 – 3 mm thickness. The mesenteric fat slices are then placed on to the SWIR-OPI imaging device for examinations. Based on the SWIR-OPI images, dark structures are extracted and placed into pathology cassettes. Extracted specimens then undergo standard histology processes and are confirmed by Board Certified pathologists to be human mesenteric lymph nodes.

### Imaging procedure

The SWIR-OPI instrumentation setup consists of an illumination path and a detection path. The illumination path is constructed by using a SWIR LED at 1550 nm (M1550D2, Thorlabs) that is collimated with a condenser lens (ACL25416U-B, Thorlabs). A glass diffuser (DG10-600-B, Thorlabs) is placed in front of the condenser lens in order to create a homogeneous illumination. A linear polarizer (LPNIRC100-MP2, Thorlabs) is placed in front of the glass diffuser in order to create linearly polarized SWIR light. Because both the linear polarizer and the glass diffuser are mounted inside optical tubes that have standard SM threading, the mounted linear polarizer can be rotated to change the polarization direction of the illumination. The SWIR LED is driven by an LED driver (LEDD1B, Thorlabs) operated at different current levels for different cameras in order to achieve optimal brightness in the images. For example, the LED driver was set to approximately 300 mA when using a cooled InGaAs camera and to approximately 1200 mA when using an uncooled InGaAs camera.

The detection path is constructed by placing another linear polarizer (LPNIRC100-MP2, Thorlabs) in front of a custom-built object space telecentric lens. The specification of the object space telecentric lens can be found in the Supplementary Document. Similar to the illumination path, the linear polarizer in front of the object space telecentric lens is also mounted in an optical tube with standard SM threading. Both a scientific-grade, liquid nitrogen-cooled InGaAs camera (2D-OMA V:320, Princeton Instruments) and a consumer-grade, uncooled InGaAs camera (SU320M-1.7RT, Sensors Unlimited) are used for detection. Both cameras have 320 × 256 resolution, and the exposure time was set to be 5 ms. While the performance of each camera under low light conditions is significantly different, the image quality produced from each camera for SWIR-OPI is almost identical. This is a key advantage of SWIR-OPI compared to other fluorescence/photoluminescence techniques in the SWIR range – fluorescence/photoluminescence requires high sensitivity/low noise SWIR cameras, which are extremely expensive, because fluorescence/photoluminescence signals are relatively weak, and are limited by the maximum illumination intensity that animals or humans are allowed to be exposed to (around 100 mW/cm^2^). SWIR-OPI, on the other hand, is based on reflection of light, which is a far more efficient process than fluorescence/photoluminescence. Quantitatively, the SWIR-OPI images that appear in this article are illuminated at approximately 1 mW/cm^2^, which is only 1% of the maximum allowed amount of light intensity exposure. The cancer study and the human mesenteric lymph node imaging studies were carried out using the consumer-grade, uncooled InGaAs camera (SU320M-1.7RT, Sensors Unlimited).

The illumination path and the detection path are set up based on Figure 1 in the main text. In practice, the relative angle between the illumination and detection paths does not significantly change the imaging results.

During live animal imaging sessions, the animals (i.e., mice and rats) are kept under anesthesia the entire time. Given that inguinal lymph nodes are anatomically located in the crease between the thigh and the torso, the human operator needs to be able to use a finger to carefully position the animal to make sure that the inguinal lymph nodes are in the field-of-view. 300 images are taken by recording a continuous video. During the video recording process, the imaged animal was slightly tilted back and forth in order to obtain multiple angles of lymph node views. Imaging two inguinal lymph nodes on both sides of an animal takes approximately 2 minutes. A daily cancer study of 30 mice takes approximately 2 hours to image all animals.

### Monte Carlo Simulation of SWIR-OPI

A simulation of linearly polarized light propagation in human skin tissue is analyzed using ray-tracing software and the Monte Carlo method. Additionally, the same technique is used to describe light distribution in biological tissues. The linearly polarized 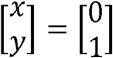 light travelling in the +z direction in skin tissue is investigated using ray-tracing software TracePro. The layers of skin tissue are produced by the software. In the simulation, the light source is defined as a point source emitting 10^7^ photons per second. The distance between the light source and the tissue is 1.9625 mm. The transverse area of the tissue is 1 mm × 1 mm, while the radius of the beam spot hitting the tissue surface is 10 μm. The polarization state of the light is investigated for various wavelengths at different skin tissue depths of 0.075 mm, 0.5 mm, 0.75 mm, 1 mm, 1.5 mm, and 2 mm. Rays emitted from the light source are incident on the skin tissue, and they are parallel to the surface normal and to the optical axis. These rays propagate within the skin tissue, and only light remaining inside the tissue is investigated for its polarization state. We use a receiver with a surface area of 1 mm × 1 mm to collect the light propagating within the tissue and neglect the light that leaves the tissue due to scattering.

As shown in Table 1 below, linearly polarized light with longer wavelengths maintains its linear polarization after interacting with a thicker section of tissue. This result is consistent with previous reports that SWIR light scatters less when interacting with biological tissue, compared to visible and near-infrared light.

### Deep learning-based image segmentation

The source codes of the deep learning algorithms that were used to train and segment lymph nodes in SWIR-OPI images are included in the Appendix section of the Supplementary Information.

Using a U-Net based deep learning algorithm, we measure the sizes of lymph nodes over the course of cancer progression. We show an example in Table 2 of the sizes of both the tumor draining lymph node and the contralateral lymph nodes after orthotopic implantation of 4T1 tumor cells, segmented and measured by the deep learning algorithm. As expected, the tumor draining lymph node grew significantly larger in size than the non-tumor draining lymph node. This process of segmentation and measurement is repeated for all mice in different study groups, and the results are discussed in detail in the following section.

**Table 2.**
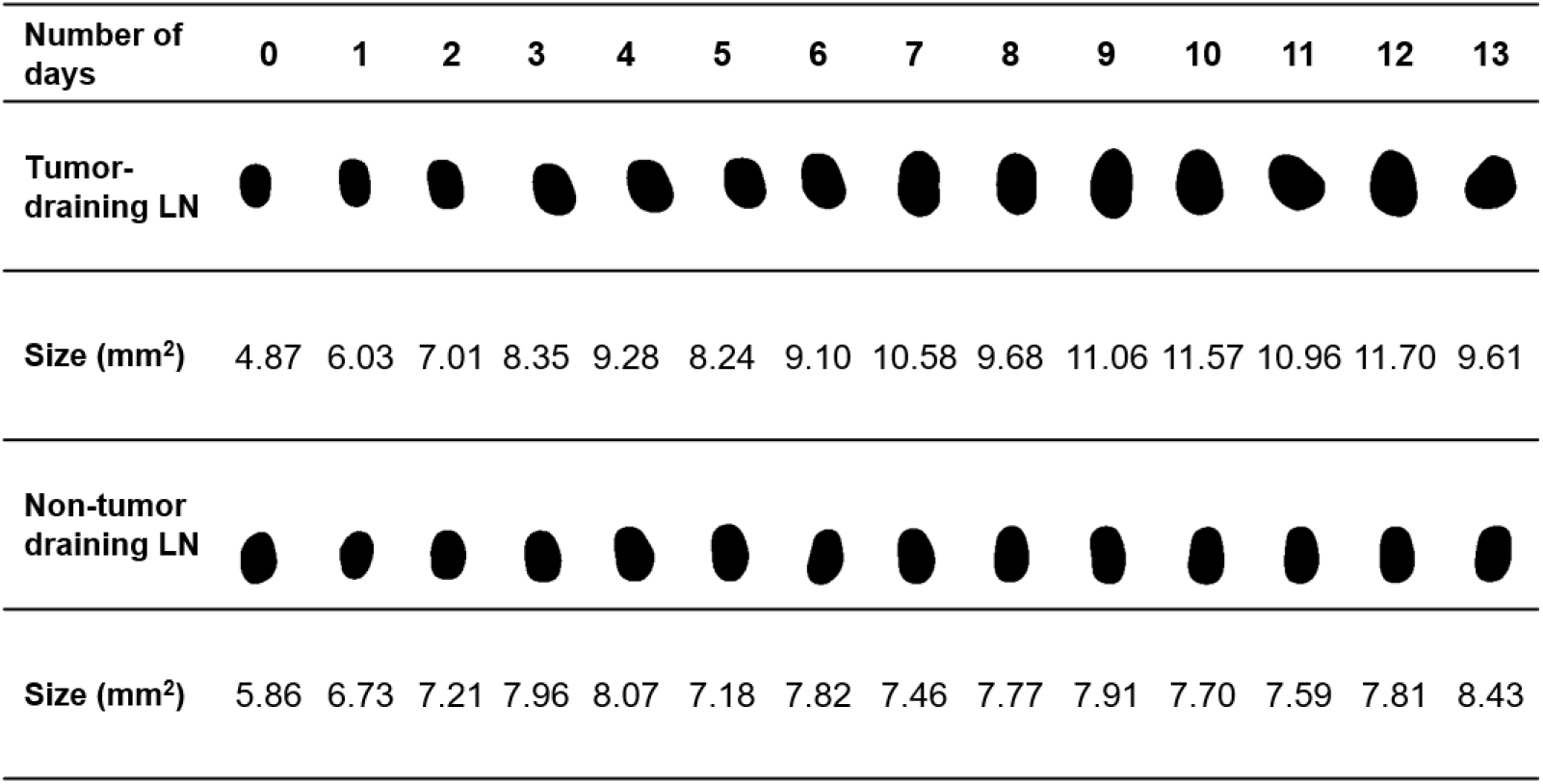
Exemplary lymph node images as segmented by U-Net and the corresponding lymph node size measurements.

| Number of days | 0 | 1 | 2 | 3 | 4 | 5 | 6 | 7 | 8 | 9 | 10 | 11 | 12 | 13 |
| --- | --- | --- | --- | --- | --- | --- | --- | --- | --- | --- | --- | --- | --- | --- |
| Tumor-draining LN |  |  |  |  |  |  |  |  |  |  |  |  |  |  |
| Size (mm<sup>2</sup>) | 4.87 | 6.03 | 7.01 | 8.35 | 9.28 | 8.24 | 9.10 | 10.58 | 9.68 | 11.06 | 11.57 | 10.96 | 11.70 | 9.61 |
| Non-tumor draining LN |  |  |  |  |  |  |  |  |  |  |  |  |  |  |
| Size (mm<sup>2</sup>) | 5.86 | 6.73 | 7.21 | 7.96 | 8.07 | 7.18 | 7.82 | 7.46 | 7.77 | 7.91 | 7.70 | 7.59 | 7.81 | 8.43 |

## Acknowledgments

We appreciate Dr. Ngozi Eze for her help in editing the manuscript. We are also grateful for Quinnipiac University’s support for providing the facility for human specimen studies.

## Funding

MIT Koch Institute Marble Center for Cancer Nanomedicine (AMB)

MIT Koch Institute Cancer Center Support (core) Grant #P30-CA14051 (AMB)

MIT Koch Institute Frontier Research Award (AMB)

National Institute of Health (AMB, R01CA214913)

Rullo Family MGH Research Scholar Award (TPP)

Natural Science Fundation (DZC, CCF-1617735)

## Author contributions

Conceptualization: ZL, AMB

Methodology: ZL

Investigation: ZL, JWW, SH, YH, SGS, YZ

Visualization: ZL

Funding acquisition: AMB

Project administration: AMB

Supervision: AMB, TPP, PTH

Writing – original draft: ZL, AMB

Writing – review & editing: ZL, AMB

## Competing Interests

Patents are assigned for the development of SWIR-OPI with inventors: Z.L. and A.M.B. All other authors declare no conflict of interest.

## Data Availability

The authors declare that the data supporting the findings of this study are available within the paper and its Supplementary Information. All other relevant data that support the findings of this study are available from the corresponding author upon request.

