## Supplementary Information for "A new label-free optical imaging method for the lymphatic system enhanced by deep learning"

### 1. Optical performance calibration

#### 1.1 Imaging lens setup:

We built an object space telecentric SWIR imaging lens by combining a commercially available Navitar 50 mm F/1.4 SWIR lens with an extension tube and an iris. In order to achieve object space telecentricity, the pinhole needs to be located at the back focal plane of the imaging lens. We achieve this by carefully selecting the correct extension tube and adapter and by fine-tuning the distance by placing a retainer ring inside the extension tube.

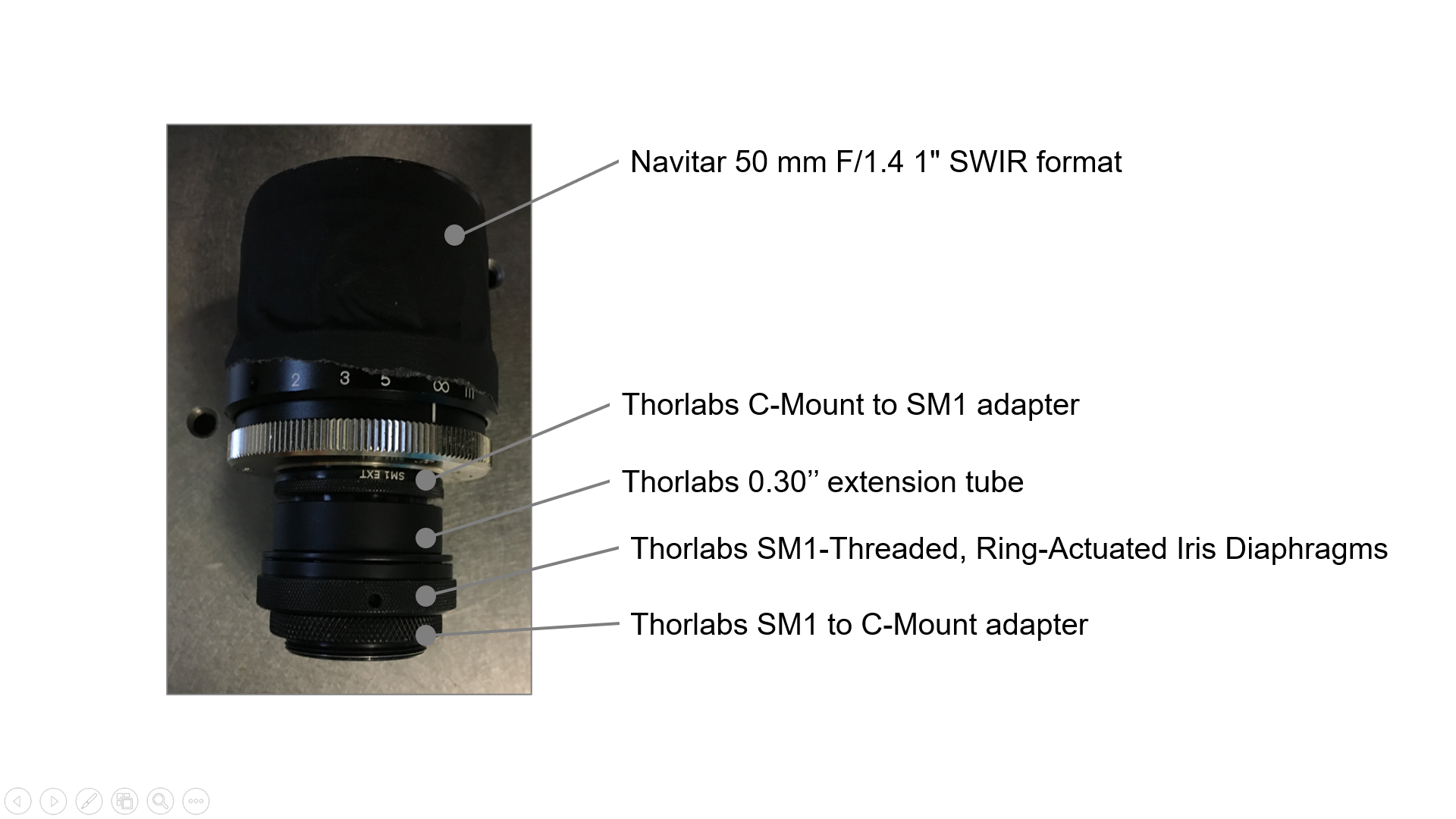

Figure S1. Home-built object space telecentric SWIR imaging lens

We measured the basic optical performance of this imaging lens as follows:

- Field-of-view: 15.7 mm x 12.8 mm
- Working Distance: ~ 60 mm

The field-of-view (magnification) is optimized to be high enough to offer enough image resolution of lymph nodes, and low enough to offer enough depth-of-field to ensure that the entire lymph node can be in focus at the same time.

The working distance is optimized to offer enough clearance between the front surface of the imaging optics, the illuminator, and the imaged animals.

Other optical parameters, including imaging resolution, telecentricity, and depth-of-field, can also be optimized by changing the size of the iris, as shown in Figure S1. The ring-actuated iris diaphragm (SM1D12C, aperture size: 5 mm, Thorlabs) has 12 pinhole stops. We measured the performance of these optical parameters at each of the 12 stops, and determined that Stop 5 corresponds to an optimal iris size. The specific measurements are shown below in Sections 2.2 and 2.3 of this document.

#### 1.2 Benchtop imaging characterization:

##### 1.2.1 Resolution

We imaged a Negative 1951 USAF test target (R1DS1N, Thorlabs) and measured the imaging resolution at each pinhole stop, shown in Figure S2 as follows:

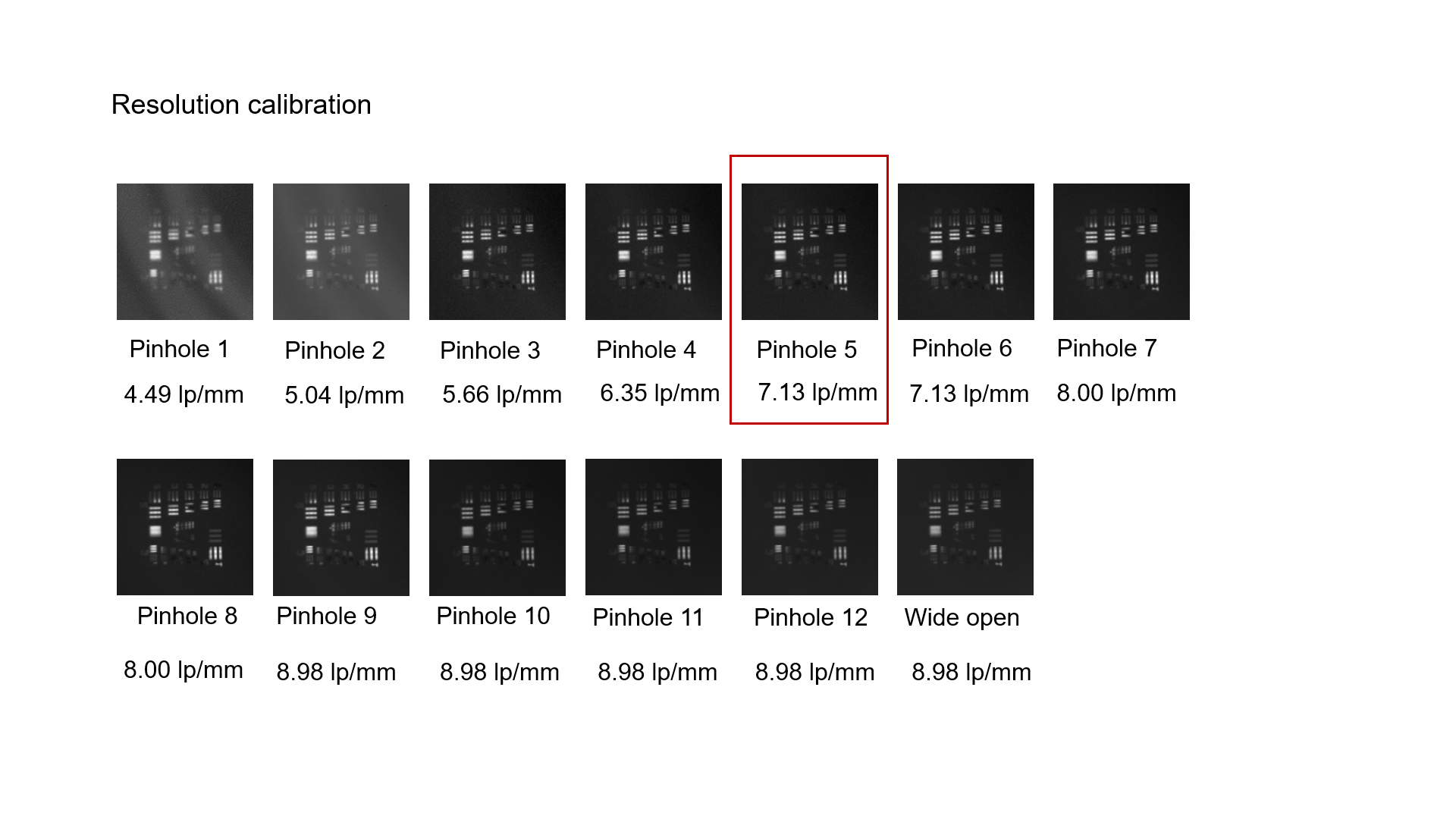

Figure S2. Resolution measurements at different pinhole sizes

##### 1.2.2 Telecentricity

In order to measure the telecentricity and depth-of-field of our imaging system, we assembled a custom imaging target (Figure S4) by attaching a standard imaging paper target, DSLRKIT Lens Focus Calibration Tool, to a 45° mounting plate (AP45, Thorlabs). The imaged lines are aligned to be 45° to the horizontal plane, i.e., parallel to the slope. This custom part creates an evenly spaced imaging target at different heights. With telecentric imaging systems, objects with the same size but at different distances to the lens would appear as the same size in pictures, whereas with regular imaging systems, objects closer to the lens would appear larger and the objects farther from the lens would appear smaller. Therefore, with our custom-built system, a comparison between the lines at the top of the image and the interval sizes at the bottom of the image can serve as a metric of the extent of telecentricity of the imaging system.

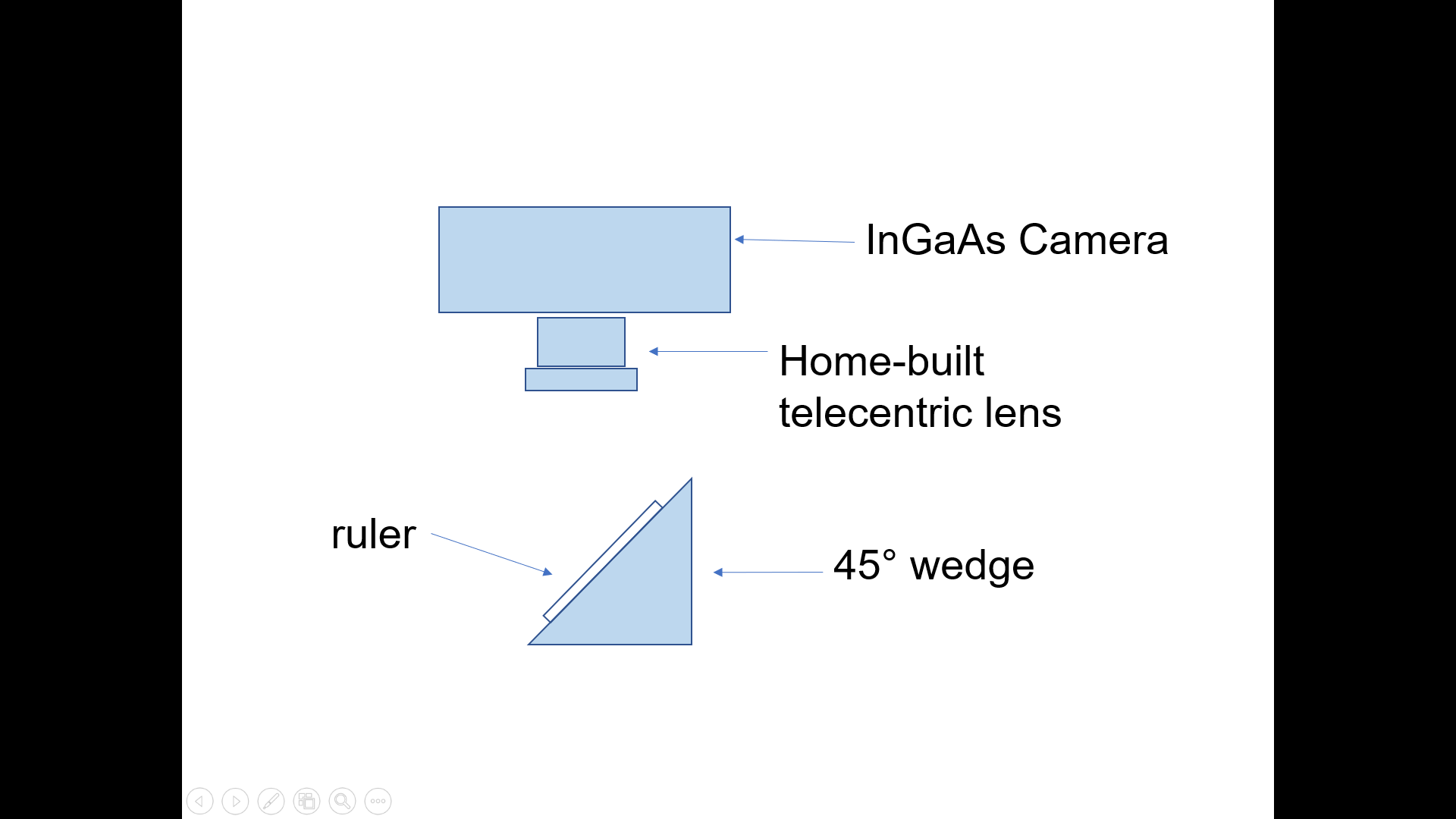

Figure S3. Telecentricity and depth-of-field measurement setup

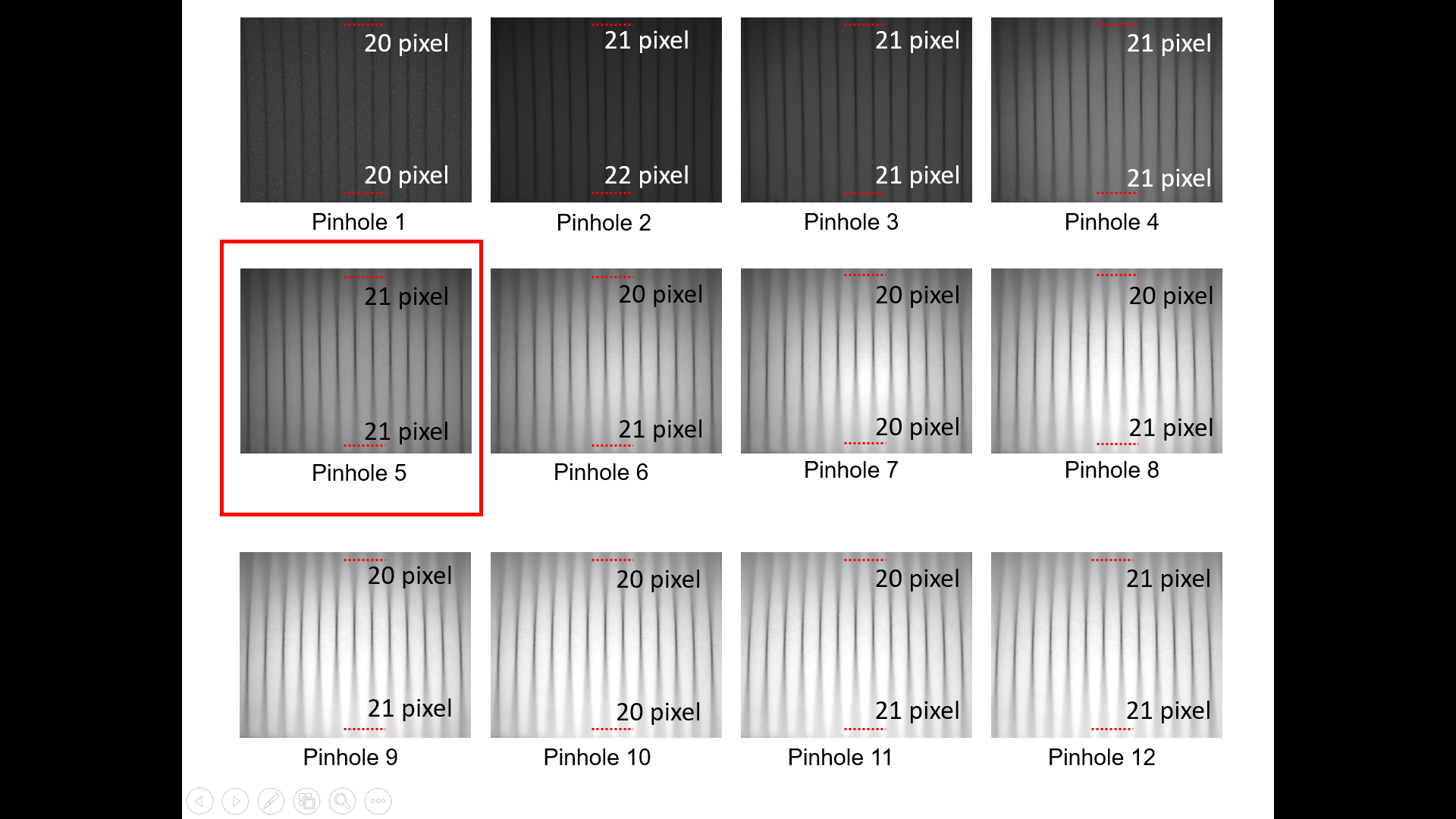

Figure S4. Telecentricity measurement of the home-built object-space telecentric lens. The numbers of pixels between two lines are compared at different apertures. The distances between all lines are 1 mm.

As shown in Figure S4, our custom-built lens shown in Figure S1 demonstrates good telecentricity at several different aperture sizes. This measurement demonstrates that our home-built imaging lens has sufficient telecentricity to allow us to track and measure the sizes of mouse lymph nodes in a consistent and reliable manner over time.

##### 1.2.3 Depth-of-field:

We also used the same imaging setup shown in Figure S4 to measure the depth-of-field of the imaging system. In order for the entire lymph node to be in focus, the depth-of-field of the imaging system needs to be larger than the size of lymph nodes, which is approximately 2 mm. At the pinhole stop 5 position, the depth of field is determined to be 6.8 mm (Figure S5), which meets this requirement.

`
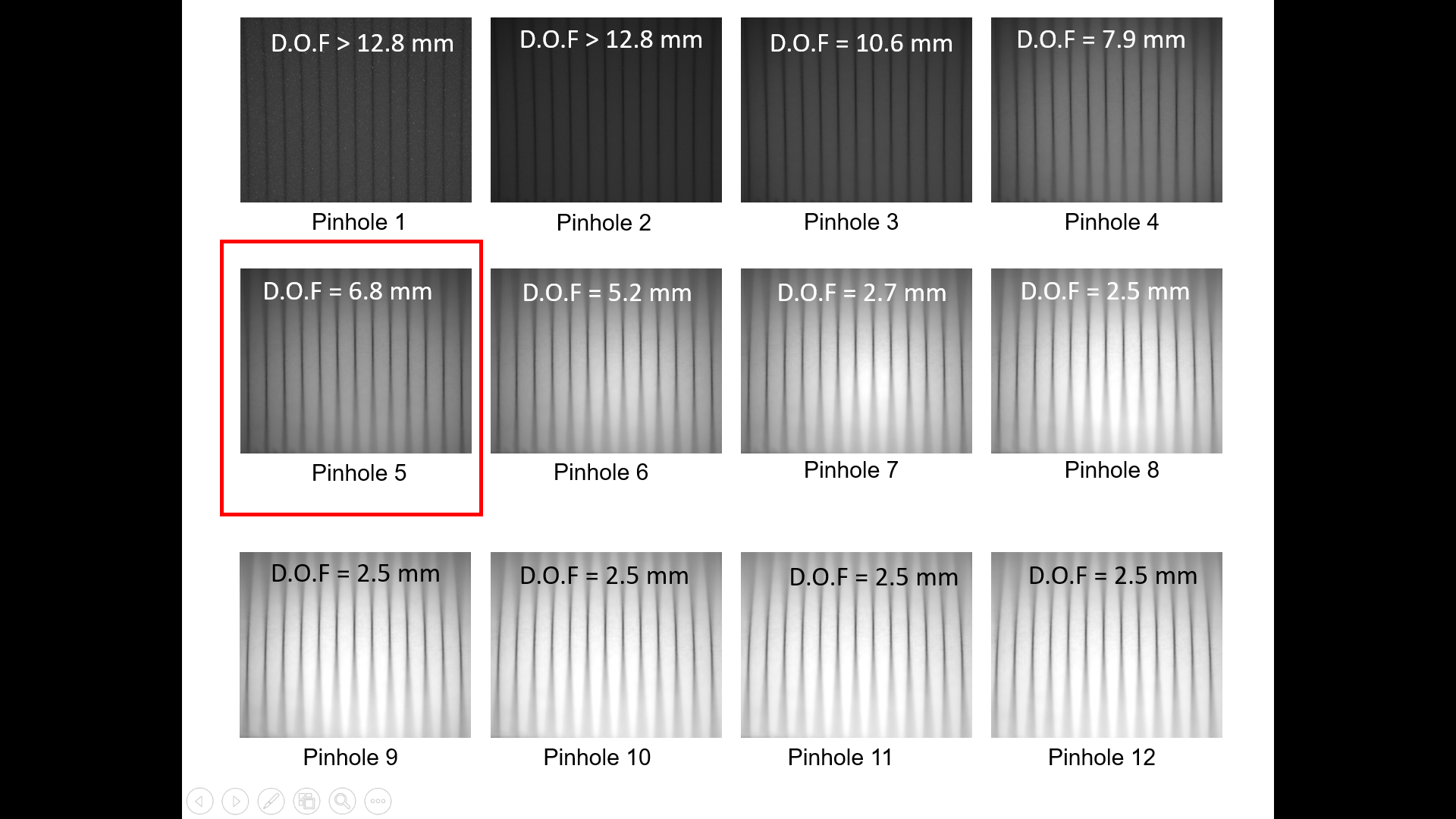

Figure S5. Depth-of-field measurements at different pinhole openings

#### 1.3 *In vivo* imaging characterization

##### 1.3.1 *In vivo* imaging resolution

We also measured the imaging resolution *in vivo*. The imaging resolution was measured by taking line profiles across the thinnest features in the image and fitting the line profiles into a Gaussian curve. The full-width half-maximum (FWHM) is calculated to determine the imaging resolution. The resolution is 65 µm for *in vivo* through-skin measurement of inguinal lymph nodes (Figure S7), and 80 µm for *ex vivo* measurement of mesenteric lymph nodes. This FWHM measurement is consistent with the results obtained from measurements on a 1951 USAF imaging target at 7.1 lp/mm, leading to a resolution of ~70 µm. This result is notable because it demonstrates the feasibility of SWIR for deep-tissue imaging. Even with intact mouse skin on top of the lymph nodes in a noninvasive *in vivo* setting, we achieved the same imaging resolution through the skin as imaging standard targets on the surface of the skin. The layers of mouse skin do not significantly affect our SWIR-OPI imaging resolution.

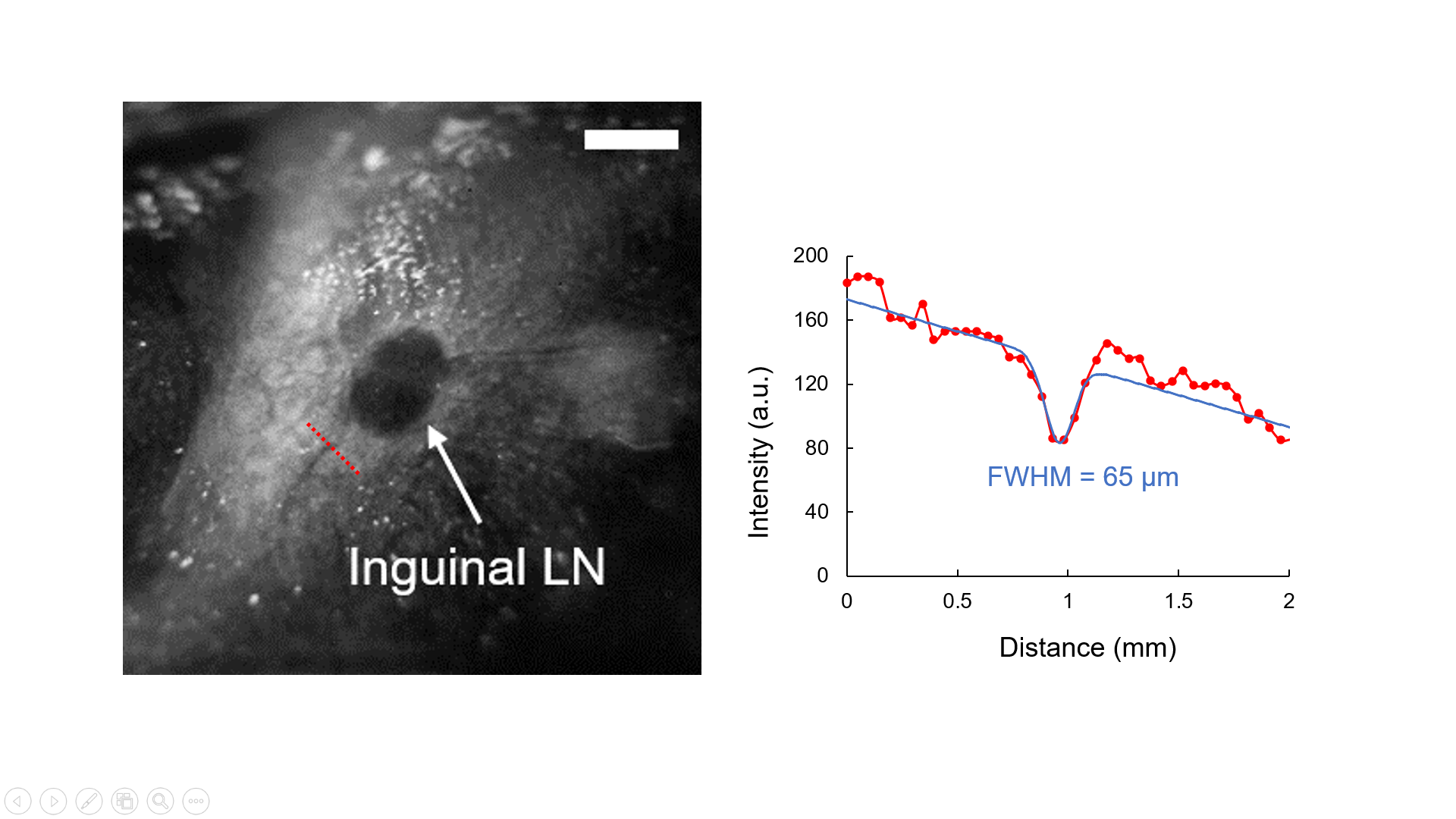

Figure S6. Imaging resolution during animal imaging sessions (Scale bar = 2 mm)

When comparing SWIR-OPI to other imaging techniques, the most common modality for imaging mouse lymph nodes is currently ultrasound imaging. In Figure S8, we conducted mouse inguinal lymph node imaging with ultrasound. When comparing to Figure S8 to S7, a clear distinction can be drawn between SWIR-OPI and ultrasound imaging, with SWIR-OPI producing images of superior contrast based on subjective visual perception.

In both imaging modalities, the contrast of the lymph node is visibly clear. A common problem in ultrasound imaging for lymph nodes is that other features can be easily mistaken as lymph nodes, such as blood vessels, bones, etc.^1^ The appearance of these extraneous features is an inherent limitation of ultrasound imaging as a cross-sectional imaging modality. Ultrasound, however, has greater imaging depth. The limited penetration depth of SWIR-OPI compared to ultrasound is not limiting when features of interest are close to the imaging surface.

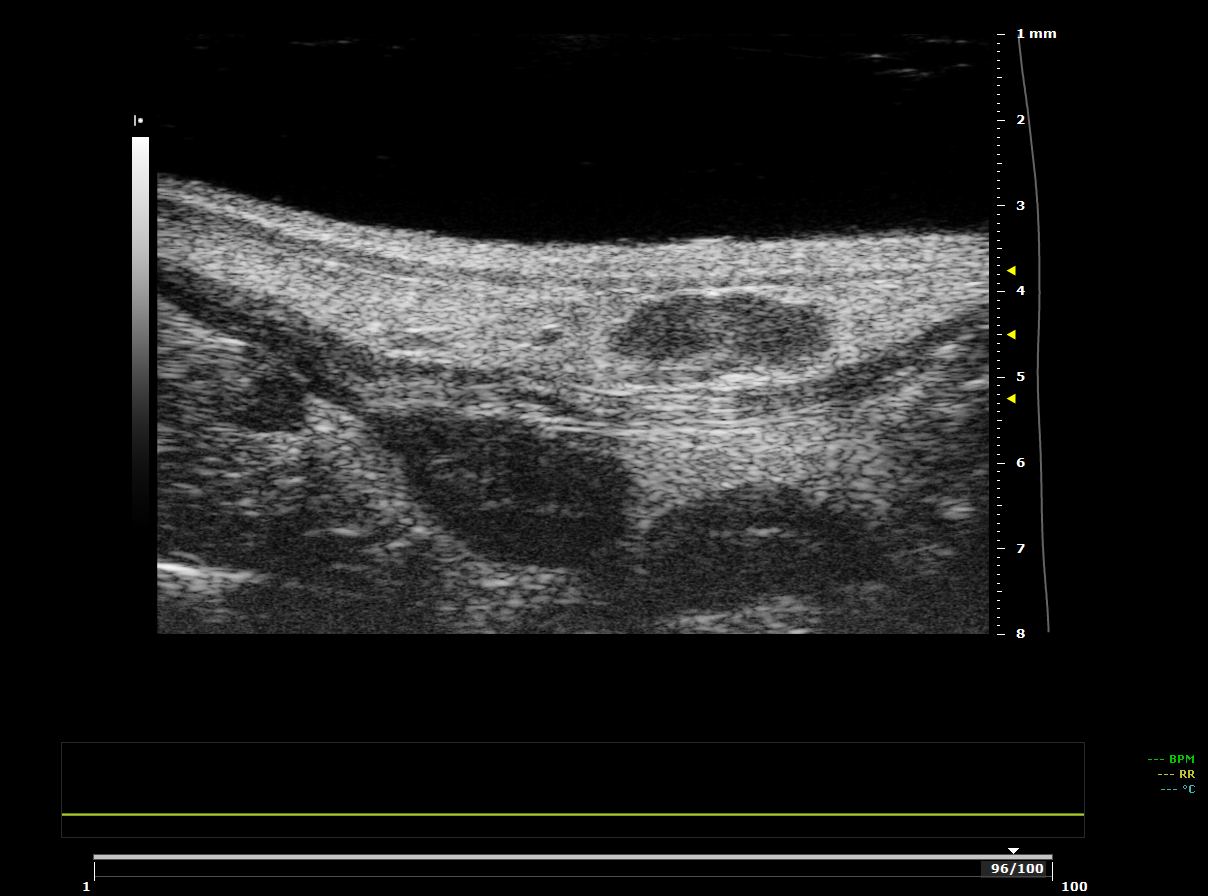

Other hypoechoic features that could be mistaken as lymph nodes

Inguinal lymph node node

Figure S7. Ultrasound image of mouse inguinal lymph node

The en-face view from SWIR-OPI is also more intuitive than ultrasound and the results are easier to interpret, because it is not a cross-sectional imaging modality. The resemblance of SWIR-OPI to visual perception makes it an ideal tool for image-guided procedures on tissues close to the surface. To demonstrate this advantage, we also performed *in vivo* SWIR-OPI-guided injection, which is commonly performed using ultrasound guidance in mouse models. As shown in Movie S1, lymph node injection can be easily performed when assisted by SWIR-OPI. Unlike ultrasound imaging, SWIR-OPI is a non-contact and intuitive modality, requiring significantly less training compared to ultrasound imaging.

##### 1.3.2 Wavelength dependency

The most unique feature of SWIR-OPI is that it offers strong natural contrast between lymph nodes and surrounding fat tissue. To quantify this contrast performance at different wavelengths, we measured the Weber contrast by applying Equation 1:

$Weber Contrast= \frac{Luminance difference}{Average luminance}$ ………………………….. Equation 1

The difference in luminance is taken by subtracting the average intensity of pixel readings in the regions marked with a blue box in the images below by the average intensity of pixel readings in the regions marked with a red and blue box in the images. The average luminance is the mean of the average intensity in the red box and the blue box regions. All images are processed using ImageJ software.

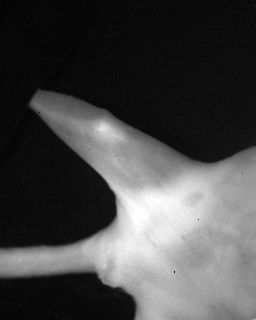

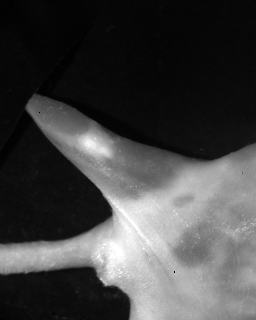

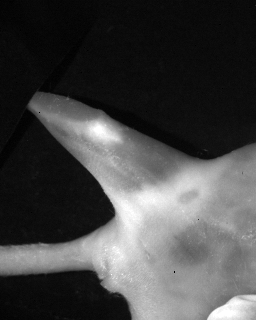

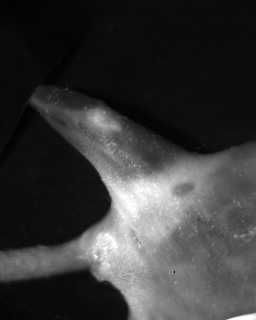

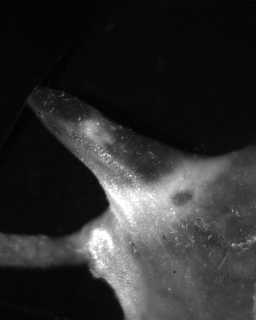

A

B

C

D

E

**1000 – 1050 nm**

**1150 – 1200 nm**

**1250 – 1300 nm**

**1375 – 1425 nm**

**1525 – 1575 nm**

**Contrast = 0.24**

**Contrast = 0.40**

**Contrast = 0.36**

**Contrast = 1.19**

**Contrast = 1.68**

Figure S9. SWIR-OPI images obtained at different wavelength ranges. Images are taken with an incandescent light source and different bandpass filters from Thorlabs and Edmund Optics.

##### 1.3.3 Polarization dependency

Orthogonal polarization imaging is particularly useful in the SWIR range, due to the reduced scattering effect of SWIR light. A detailed simulation is provided in Section 3 of this document. Based on the results that we obtained from the wavelength dependency study, here we present a comparison between regular SWIR imaging at 1550 nm using unpolarized light and SWIR-OPI at 1550 nm. As shown in Figure S10, the lymph nodes are not visible through intact mouse skin even though imaging was performed at 1550 nm, whereas lymph nodes are clearly visible through intact mouse skin using SWIR-OPI.

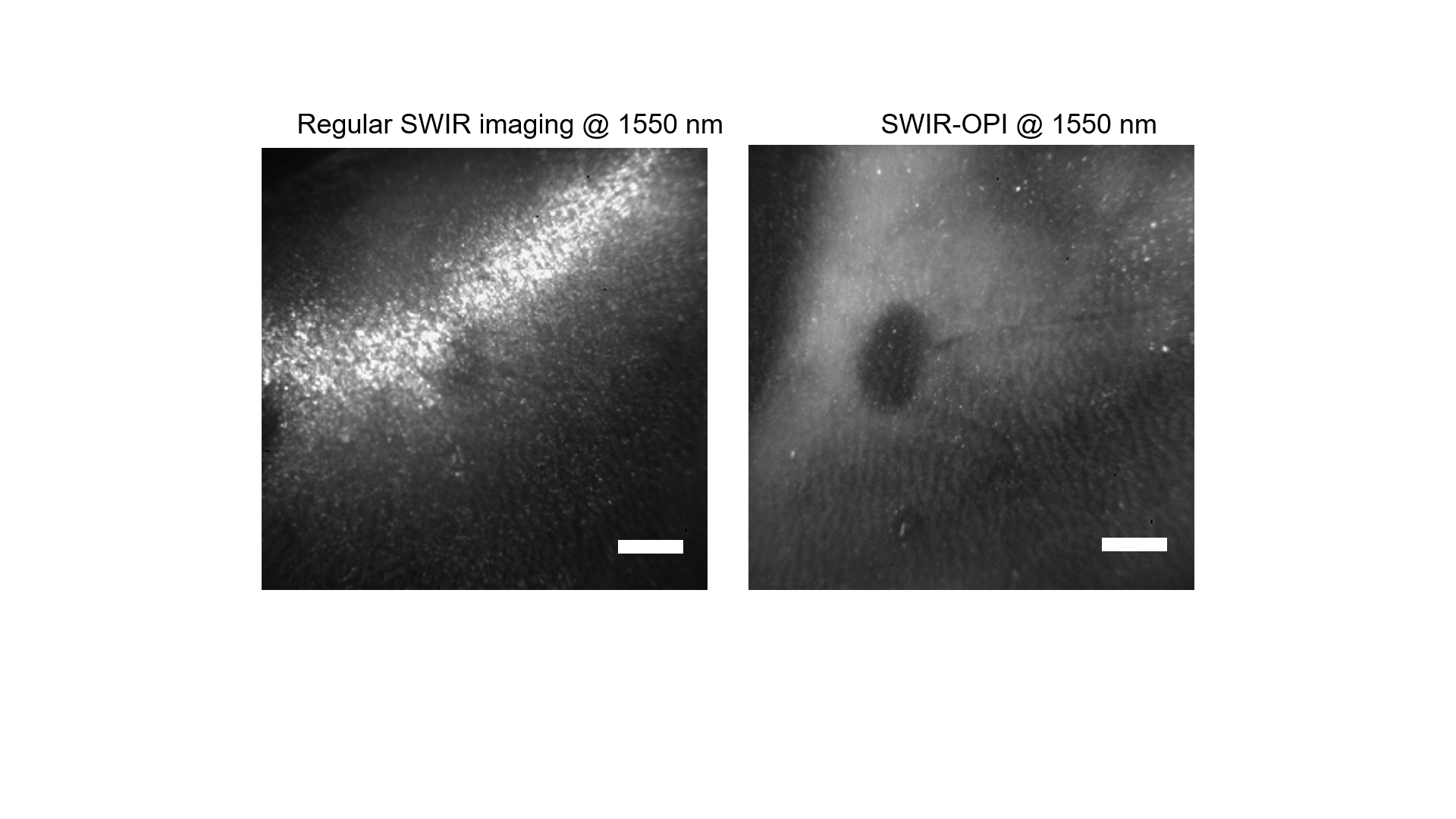

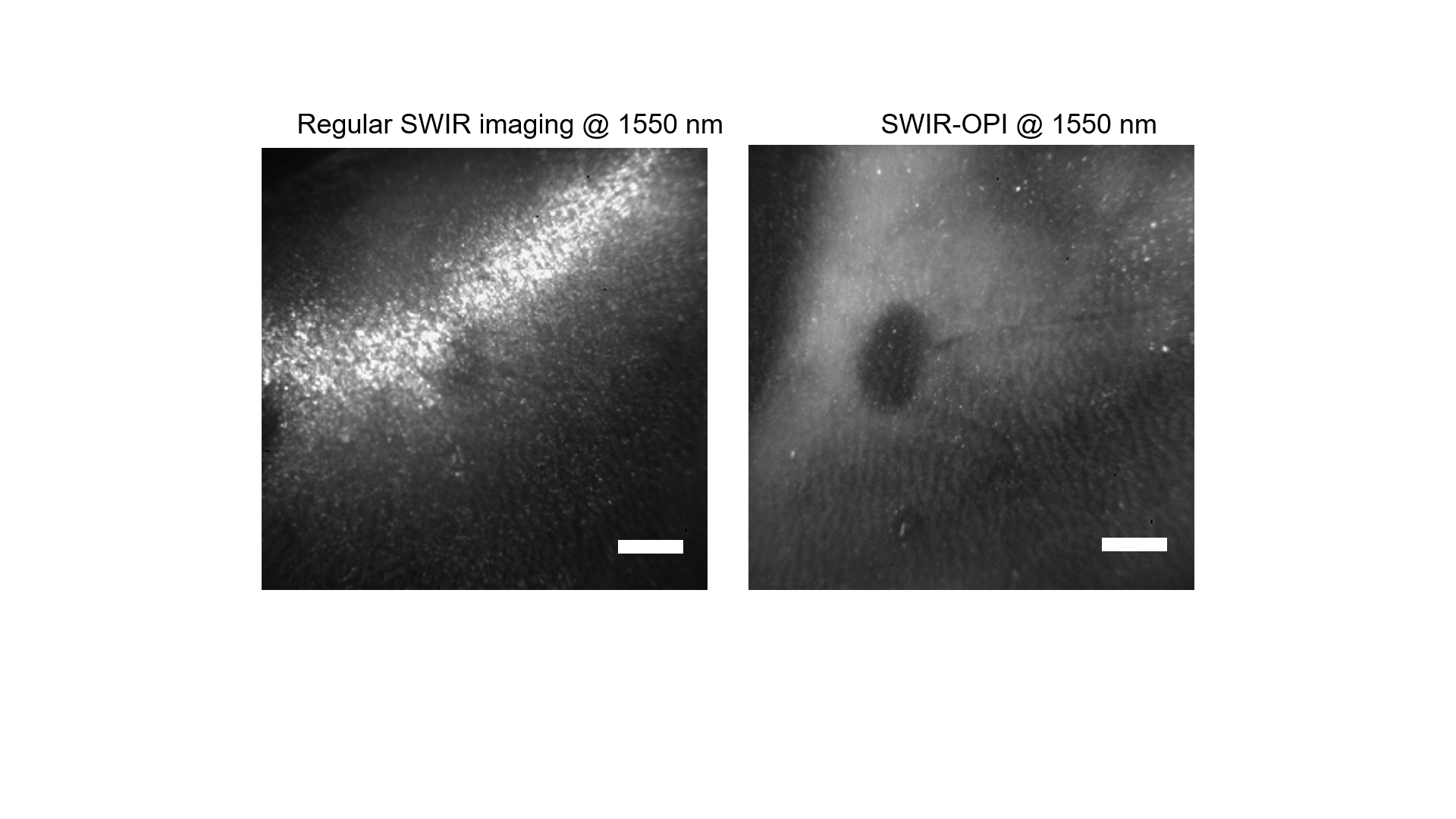

A

B

Figure S10. Comparison between regular SWIR imaging using unpolarized light at 1550 nm and SWIR-OPI at 1550 nm.

### 2. 4T1 cancer study results

#### 2.1 Size measurement results

Using the algorithms mentioned above, we systematically tracked changes to lymph nodes over the course of cancer progression. In the spaghetti plots shown in Figure S12, we can clearly see the trend that tumor-bearing mice that eventually developed lymph node metastasis have overall a larger than average increase in the size of tumor draining lymph nodes than the tumor-bearing mice that did not develop lymph node metastasis. The full reports on the size measurement results are listed in Tables S3 to S8.

Contralateral lymph node size changes

|  | Mouse-M1(283) | Mouse-M2(292) | Mouse-M3(285) | Mouse-M4(286) | Mouse-M5(295) | Mouse-M6(256) | Mouse-M7(257) | Mouse-N1(281) | Mouse-N2(284) | Mouse-N3(287) | Mouse-N4(289) | Mouse-N5(290) | Mouse-N6(291) | Mouse-N7(293) | Mouse-N8(195) | Mouse-N9(258) | Mouse-N10(260) | Mouse-N11(266) | Mouse-N12(269) | Mouse-C1(187) |
| --- | --- | --- | --- | --- | --- | --- | --- | --- | --- | --- | --- | --- | --- | --- | --- | --- | --- | --- | --- | --- |
| 0 | 0% | 0% | 0% | 0% | 0% | 0% | 0% | 0% | 0% | 0% | 0% | 0% | 0% | 0% | 0% | 0% | 0% | 0% | 0% | 0% |
| 1 | 0.0596139% | 11.656% | 29.6374% | -0.351055% | 36.6648% | -22.9779% | -3.53179% | 2.44019% | 42.9627% | 9.59001% | 29.1698% | 1.19257% | 3.92576% | 21.32% | 19% | 3.00429% | -1.38462% | 0.447227% | -26.2419% | -1.42506% |
| 2 | 16.7335% | 8.20186% | 28.7763% | 16.3126% | 32.7306% | -16.2684% | 5.14632% | 13.3937% | 43.2256% | 13.4402% | 29.1698% | 7.29051% | 12.8147% | 21.1897% | -9% | 39.485% | 8.61538% | 12.4329% | -13.2829% | 4.43567% |
| 3 | 37.4557% | 31.0903% | 28.7763% | 14.1929% | 48.4042% | 34.6507% | 9.28355% | 20.2795% | 41.706% | 24.4431% | 37.9072% | 10.6904% | 21.3657% | 27.0361% | 60.4444% | 49.8927% | -5.46154% | 3.39893% | 2.69978% | 1.77555% |
| 4 | 46.49% | 25.2704% | 51.918% | 16.34% | 56.1388% | 49.4485% | 41.776% | 26.205% | 53.2782% | 27.316% | 31.1418% | 17.8708% | 17.5887% | 27.0251% | 70.1111% | 93.6695% | 23.0769% | 33.8998% | 25.486% | 19.9892% |
| 5 | 45.4436% | 25.4396% | 36.9656% | 15.2281% | 55.0147% | 70.0368% | 37.5378% | 24.6999% | 35.9913% | 28.4132% | 39.4206% | 15.4336% | 41.8754% | 28.2551% | 89.2222% | 82.4034% | 39.6154% | 44.2755% | 26.7819% | 17.2865% |
| 6 | 59.0064% | 30.2863% | 46.2228% | 10.6822% | 46.7443% | 42.9228% | 58.5267% | 30.9724% | 72.3063% | 32.0313% | 29.6603% | 23.3769% | 53.8316% | 30.5059% | 63.4444% | 81.867% | 48.5385% | 24.5975% | 22.1382% | 29.1582% |
| 7 | 52.7598% | 26.2378% | 39.2125% | 17.3403% | 46.8798% | 30.6985% | 75.1766% | 23.0268% | 67.2247% | 28.5264% | 26.6959% | 31.9168% | 71.0124% | 31.883% | 54.8889% | 75.2146% | 23% | 14.6691% | 19.2225% | 25.2234% |
| 8 | 44.4605% | 19.9654% | 43.696% | 8.01856% | 39.3868% | 46.4154% | 50.0505% | 22.1277% | 53.3442% | 15.6043% | 35.7507% | 23.618% | 69.8409% | 24.1472% | 50.4444% | 103.541% | 23.0769% | 12.4329% | 31.3175% | 19.1232% |
| 9 | 56.0863% | 24.6925% | 47.5804% | 8.62582% | 44.6776% | 70.4044% | 62.0585% | 24.0617% | 70.3225% | 25.019% | 42.4381% | 24.16% | 80.4437% | 36.7054% | 71.1111% | 81.5451% | 44.6154% | 5.54562% | 27.9698% | 7.10887% |
| 10 | 56.8481% | 36.6304% | 45.4489% | 12.188% | 49.1377% | 46.875% | 69.0212% | 25.4533% | 79.0666% | 34.2648% | 33.7737% | 35.1177% | 67.5542% | 45.4246% | 56.3333% | 115.451% | 39.0769% | 17.8891% | 12.095% | 22.0546% |
| 11 | 44.3884% | 42.5211% | 44.9295% | 12.5563% | 47.802% | 34.0074% | 66.8012% | 22.417% | 73.0219% | 29.5221% | 36.4915% | 39.4484% | 79.169% | 60.9338% | 59.8889% | 95.8155% | 20.6923% | 9.66011% | 22.3542% | 27.5151% |
| 12 | 50.3084% | 24.9786% | 34.3458% | 5.64471% | 41.3056% | 65.2574% | 87.891% | 17.3443% | 74.5199% | 28.5039% | 32.2231% | 39.6715% | 70.8839% | 33.6714% | 66.1111% | 108.906% | 9.38462% | -0.178891% | 11.3391% | 37.5412% |
| 13 | 47.6232% | 35.3478% | 57.391% | 6.75537% | 21.53% | 20.2206% | 80.9284% | 20.0182% | 69.8502% | 23.5787% | 34.6882% | 31.4294% | 74.502% | 32.0718% | 44.4444% | 71.7811% | 12.1538% | -5.99284% | -3.02376% | 16.7834% |

Tumor draining lymph node size changes

|  | Mouse-M1(283) | Mouse-M2(292) | Mouse-M3(285) | Mouse-M4(286) | Mouse-M5(295) | Mouse-M6(256) | Mouse-M7(257) | Mouse-N1(281) | Mouse-N2(284) | Mouse-N3(287) | Mouse-N4(289) | Mouse-N5(290) | Mouse-N6(291) | Mouse-N7(293) | Mouse-N8(195) | Mouse-N9(258) | Mouse-N10(260) | Mouse-N11(266) | Mouse-N12(269) | Mouse-C1(187) |
| --- | --- | --- | --- | --- | --- | --- | --- | --- | --- | --- | --- | --- | --- | --- | --- | --- | --- | --- | --- | --- |
| 0 | 0% | 0% | 0% | 0% | 0% | 0% | 0% | 0% | 0% | 0% | 0% | 0% | 0% | 0% | 0% | 0% | 0% | 0% | 0% | 0% |
| 1 | 16.2385% | 32.4514% | 32.8709% | 16.1961% | 22.2113% | 29.8165% | 17.4579% | 16.5723% | 43.2687% | 16.6468% | 11.9764% | 3.92576% | 12.9998% | 3.87127% | 36.8137% | -3.47734% | -0.733945% | -7.07364% | 4.87552% | -4.24959% |
| 2 | 33.973% | 34.4023% | 48.9348% | 21.2198% | 33.6458% | 21.7125% | 49.1577% | 28.7552% | 54.06% | 35.2868% | 11.9764% | 12.8147% | 10.2775% | 8.55162% | 23.9424% | -0.737619% | 15.7798% | -10.0775% | -4.56432% | 0.0449926% |
| 3 | 59.7428% | 54.0153% | 83.5789% | 20.7878% | 38.7475% | 92.5076% | 37.5191% | 36.911% | 77.9232% | 44.8008% | 41.136% | 21.3657% | 41.7343% | 36.6808% | 60.036% | 17.3867% | 1.10092% | 38.9535% | 25.4149% | -6.1075% |
| 4 | 61.1777% | 66.9565% | 90.2112% | 48.1325% | 70.4692% | 109.786% | 123.124% | 48.6156% | 80.9516% | 44.6073% | 54.9551% | 17.5887% | 27.7708% | 47.8401% | 68.4068% | 57.9557% | 40.0917% | 48.8372% | 17.4274% | -10.1536% |
| 5 | 63.9679% | 72.1713% | 90.1043% | 54.8003% | 75.9242% | 131.651% | 91.5773% | 41.8077% | 97.307% | 54.3616% | 59.5261% | 41.8754% | 48.3193% | 26.3339% | 45.5446% | 98.6301% | 80.2752% | 45.9302% | 35.2697% | -1.35896% |
| 6 | 83.6481% | 72.6619% | 99.3645% | 93.4195% | 88.6271% | 175.688% | 101.378% | 55.1153% | 100.246% | 68.1218% | 62.8014% | 53.8316% | 57.5552% | 61.5805% | 82.1782% | 70.1791% | 79.0826% | 24.3217% | 27.2822% | 4.66543% |
| 7 | 104.458% | 101.467% | 133.245% | 70.0516% | 116.486% | 194.954% | 148.086% | 30.2118% | 110.564% | 51.6025% | 85.8517% | 71.0124% | 59.5499% | 47.1944% | 85.1485% | 92.0969% | 66.6055% | 58.3333% | 80.083% | 5.88659% |
| 8 | 119.491% | 124.693% | 123.157% | 91.1145% | 127.451% | 239.755% | 128.79% | 59.09% | 127.358% | 50.1147% | 72.3748% | 69.8409% | 78.0322% | 60.5644% | 78.3078% | 109.168% | 89.4495% | 82.2674% | 60.9959% | 0.0582258% |
| 9 | 108.982% | 83.4818% | 141.39% | 91.5367% | 118.232% | 183.18% | 145.329% | 39.2744% | 124.123% | 57.3615% | 94.8883% | 80.4437% | 85.5541% | 75.7656% | 95.1395% | 155.848% | 68.7156% | 46.8023% | 66.39% | -1.2684% |
| 10 | 123.586% | 108.716% | 161.427% | 75.4349% | 141.076% | 239.297% | 168.147% | 35.4579% | 119.082% | 32.3801% | 95.6662% | 67.5542% | 74.7396% | 70.3923% | 128.353% | 146.47% | 98.1651% | 46.6085% | 88.6929% | -0.122366% |
| 11 | 123.491% | 112.723% | 155.888% | 87.3978% | 99.1339% | 148.165% | 119.602% | 37.7579% | 105.817% | 12.2233% | 33.6495% | 79.169% | 79.5606% | 61.3531% | 69.577% | 83.667% | 73.0275% | 61.1434% | 91.4938% | 16.7687% |
| 12 | 88.3183% | 82.3865% | 149.376% | 89.8003% | 115.167% | 265.291% | 118.224% | 30.1847% | 73.0241% | 22.4527% | 72.1105% | 70.8839% | 90.4736% | 67.0336% | 101.44% | 139.41% | 92.7523% | 98.4496% | 72.9253% | 3.28843% |
| 13 | 90.3083% | 60.0091% | 108.887% | 95.4253% | 83.6334% | 147.248% | 58.0398% | 34.0014% | 84.8686% | 4.30086% | 59.8919% | 74.502% | 70.203% | 66.2365% | 36.8137% | 60.9062% | 48.6239% | 69.8643% | 72.7178% | 10.368% |

Figure S12. Spaghetti plots of cancer study data on changes in lymph node size. Orange lines represent tumor-bearing mice with metastasis in the pathology report, black lines represent tumor-bearing mice with no metastasis in the pathology report, and green lines are the control mice injected with PBS. It can be clearly seen that tumor-bearing mice with metastatic lymph nodes have overall a larger than average increase in the size of tumor draining lymph nodes.

| # of days | Mouse-M1(283) | Mouse-M2(292) | Mouse-M3(285) | Mouse-M4(286) | Mouse-M5(295) | Mouse-M6(256) | Mouse-M7(257) |
| --- | --- | --- | --- | --- | --- | --- | --- |
| 0 | 0.00% | 0.00% | 0.00% | 0.00% | 0.00% | 0.00% | 0.00% |
| 1 | 16.24% | 32.45% | 32.87% | 16.20% | 22.21% | 29.82% | 17.46% |
| 2 | 33.97% | 34.40% | 48.93% | 21.22% | 33.65% | 21.71% | 49.16% |
| 3 | 59.74% | 54.02% | 83.58% | 20.79% | 38.75% | 92.51% | 37.52% |
| 4 | 61.18% | 66.96% | 90.21% | 48.13% | 70.47% | 109.79% | 123.12% |
| 5 | 63.97% | 72.17% | 90.10% | 54.80% | 75.92% | 131.65% | 91.58% |
| 6 | 83.65% | 72.66% | 99.36% | 93.42% | 88.63% | 175.69% | 101.38% |
| 7 | 104.46% | 101.47% | 133.25% | 70.05% | 116.49% | 194.95% | 148.09% |
| 8 | 119.49% | 124.69% | 123.16% | 91.11% | 127.45% | 239.76% | 128.79% |
| 9 | 108.98% | 83.48% | 141.39% | 91.54% | 118.23% | 183.18% | 145.33% |
| 10 | 123.59% | 108.72% | 161.43% | 75.43% | 141.08% | 239.30% | 168.15% |
| 11 | 123.49% | 112.72% | 155.89% | 87.40% | 99.13% | 148.17% | 119.60% |
| 12 | 88.32% | 82.39% | 149.38% | 89.80% | 115.17% | 265.29% | 118.22% |
| 13 | 90.31% | 60.01% | 108.89% | 95.43% | 83.63% | 147.25% | 58.04% |

Table S3. Lymph node size measurements for tumor draining lymph nodes from tumor-bearing mice with lymph node metastasis in the pathology report.

| # of days | Mouse-M1(283) | Mouse-M2(292) | Mouse-M3(285) | Mouse-M4(286) | Mouse-M5(295) | Mouse-M6(256) | Mouse-M7(257) |
| --- | --- | --- | --- | --- | --- | --- | --- |
| 0 | 0.00% | 0.00% | 0.00% | 0.00% | 0.00% | 0.00% | 0.00% |
| 1 | 0.06% | 11.66% | 29.64% | -0.35% | 36.66% | -22.98% | -3.53% |
| 2 | 16.73% | 8.20% | 28.78% | 16.31% | 32.73% | -16.27% | 5.15% |
| 3 | 37.46% | 31.09% | 28.78% | 14.19% | 48.40% | 34.65% | 9.28% |
| 4 | 46.49% | 25.27% | 51.92% | 16.34% | 56.14% | 49.45% | 41.78% |
| 5 | 45.44% | 25.44% | 36.97% | 15.23% | 55.01% | 70.04% | 37.54% |
| 6 | 59.01% | 30.29% | 46.22% | 10.68% | 46.74% | 42.92% | 58.53% |
| 7 | 52.76% | 26.24% | 39.21% | 17.34% | 46.88% | 30.70% | 75.18% |
| 8 | 44.46% | 19.97% | 43.70% | 8.02% | 39.39% | 46.42% | 50.05% |
| 9 | 56.09% | 24.69% | 47.58% | 8.63% | 44.68% | 70.40% | 62.06% |
| 10 | 56.85% | 36.63% | 45.45% | 12.19% | 49.14% | 46.88% | 69.02% |
| 11 | 44.39% | 42.52% | 44.93% | 12.56% | 47.80% | 34.01% | 66.80% |
| 12 | 50.31% | 24.98% | 34.35% | 5.64% | 41.31% | 65.26% | 87.89% |
| 13 | 47.62% | 35.35% | 57.39% | 6.76% | 21.53% | 20.22% | 80.93% |

Table S4. Lymph node size measurements for contralateral lymph nodes from tumor-bearing mice with lymph node metastasis in the pathology report.

| # of days | Mouse-N1(281) | Mouse-N2(284) | Mouse-N3(287) | Mouse-N4(289) | Mouse-N5(290) | Mouse-N6(291) | Mouse-N7(293) | Mouse-N8(195) | Mouse-N9(258) | Mouse-N10(260) | Mouse-N11(266) | Mouse-N12(269) |
| --- | --- | --- | --- | --- | --- | --- | --- | --- | --- | --- | --- | --- |
| 0 | 0.00% | 0.00% | 0.00% | 0.00% | 0.00% | 0.00% | 0.00% | 0.00% | 0.00% | 0.00% | 0.00% | 0.00% |
| 1 | 16.57% | 43.27% | 16.65% | 11.98% | 3.93% | 13.00% | 3.87% | 36.81% | -3.48% | -0.73% | -7.07% | 4.88% |
| 2 | 28.76% | 54.06% | 35.29% | 11.98% | 12.81% | 10.28% | 8.55% | 23.94% | -0.74% | 15.78% | -10.08% | -4.56% |
| 3 | 36.91% | 77.92% | 44.80% | 41.14% | 21.37% | 41.73% | 36.68% | 60.04% | 17.39% | 1.10% | 38.95% | 25.41% |
| 4 | 48.62% | 80.95% | 44.61% | 54.96% | 17.59% | 27.77% | 47.84% | 68.41% | 57.96% | 40.09% | 48.84% | 17.43% |
| 5 | 41.81% | 97.31% | 54.36% | 59.53% | 41.88% | 48.32% | 26.33% | 45.54% | 98.63% | 80.28% | 45.93% | 35.27% |
| 6 | 55.12% | 100.25% | 68.12% | 62.80% | 53.83% | 57.56% | 61.58% | 82.18% | 70.18% | 79.08% | 24.32% | 27.28% |
| 7 | 30.21% | 110.56% | 51.60% | 85.85% | 71.01% | 59.55% | 47.19% | 85.15% | 92.10% | 66.61% | 58.33% | 80.08% |
| 8 | 59.09% | 127.36% | 50.11% | 72.37% | 69.84% | 78.03% | 60.56% | 78.31% | 109.17% | 89.45% | 82.27% | 61.00% |
| 9 | 39.27% | 124.12% | 57.36% | 94.89% | 80.44% | 85.55% | 75.77% | 95.14% | 155.85% | 68.72% | 46.80% | 66.39% |
| 10 | 35.46% | 119.08% | 32.38% | 95.67% | 67.55% | 74.74% | 70.39% | 128.35% | 146.47% | 98.17% | 46.61% | 88.69% |
| 11 | 37.76% | 105.82% | 12.22% | 33.65% | 79.17% | 79.56% | 61.35% | 69.58% | 83.67% | 73.03% | 61.14% | 91.49% |
| 12 | 30.18% | 73.02% | 22.45% | 72.11% | 70.88% | 90.47% | 67.03% | 101.44% | 139.41% | 92.75% | 98.45% | 72.93% |
| 13 | 34.00% | 84.87% | 4.30% | 59.89% | 74.50% | 70.20% | 66.24% | 36.81% | 60.91% | 48.62% | 69.86% | 72.72% |

Table S5. Lymph node size measurements for tumor draining lymph nodes from tumor-bearing mice without lymph node metastasis in the pathology report.

| # of days | Mouse-N1(281) | Mouse-N2(284) | Mouse-N3(287) | Mouse-N4(289) | Mouse-N5(290) | Mouse-N6(291) | Mouse-N7(293) | Mouse-N8(195) | Mouse-N9(258) | Mouse-N10(260) | Mouse-N11(266) | Mouse-N12(269) |
| --- | --- | --- | --- | --- | --- | --- | --- | --- | --- | --- | --- | --- |
| 0 | 0.00% | 0.00% | 0.00% | 0.00% | 0.00% | 0.00% | 0.00% | 0.00% | 0.00% | 0.00% | 0.00% | 0.00% |
| 1 | 2.44% | 42.96% | 9.59% | 29.17% | 1.19% | 3.93% | 21.32% | 19.00% | 3.00% | -1.38% | 0.45% | -26.24% |
| 2 | 13.39% | 43.23% | 13.44% | 29.17% | 7.29% | 12.81% | 21.19% | -9.00% | 39.48% | 8.62% | 12.43% | -13.28% |
| 3 | 20.28% | 41.71% | 24.44% | 37.91% | 10.69% | 21.37% | 27.04% | 60.44% | 49.89% | -5.46% | 3.40% | 2.70% |
| 4 | 26.20% | 53.28% | 27.32% | 31.14% | 17.87% | 17.59% | 27.03% | 70.11% | 93.67% | 23.08% | 33.90% | 25.49% |
| 5 | 24.70% | 35.99% | 28.41% | 39.42% | 15.43% | 41.88% | 28.26% | 89.22% | 82.40% | 39.62% | 44.28% | 26.78% |
| 6 | 30.97% | 72.31% | 32.03% | 29.66% | 23.38% | 53.83% | 30.51% | 63.44% | 81.87% | 48.54% | 24.60% | 22.14% |
| 7 | 23.03% | 67.22% | 28.53% | 26.70% | 31.92% | 71.01% | 31.88% | 54.89% | 75.21% | 23.00% | 14.67% | 19.22% |
| 8 | 22.13% | 53.34% | 15.60% | 35.75% | 23.62% | 69.84% | 24.15% | 50.44% | 103.54% | 23.08% | 12.43% | 31.32% |
| 9 | 24.06% | 70.32% | 25.02% | 42.44% | 24.16% | 80.44% | 36.71% | 71.11% | 81.55% | 44.62% | 5.55% | 27.97% |
| 10 | 25.45% | 79.07% | 34.26% | 33.77% | 35.12% | 67.55% | 45.42% | 56.33% | 115.45% | 39.08% | 17.89% | 12.10% |
| 11 | 22.42% | 73.02% | 29.52% | 36.49% | 39.45% | 79.17% | 60.93% | 59.89% | 95.82% | 20.69% | 9.66% | 22.35% |
| 12 | 17.34% | 74.52% | 28.50% | 32.22% | 39.67% | 70.88% | 33.67% | 66.11% | 108.91% | 9.38% | -0.18% | 11.34% |
| 13 | 20.02% | 69.85% | 23.58% | 34.69% | 31.43% | 74.50% | 32.07% | 44.44% | 71.78% | 12.15% | -5.99% | -3.02% |

Table S6. Lymph node size measurements for contralateral lymph nodes from tumor-bearing mice without lymph node metastasis in the pathology report.

| # of days | Mouse-C1(187) | Mouse-C2(197) |
| --- | --- | --- |
| 0 | 0.00% | 0.00% |
| 1 | -4.25% | -2.66% |
| 2 | 0.04% | -2.14% |
| 3 | -6.11% | 4.63% |
| 4 | -10.15% | 2.15% |
| 5 | -1.36% | 26.43% |
| 6 | 4.67% | 18.47% |
| 7 | 5.89% | 8.51% |
| 8 | 0.06% | 13.45% |
| 9 | -1.27% | 15.71% |
| 10 | -0.12% | 10.62% |
| 11 | 16.77% | 16.49% |
| 12 | 3.29% | 3.07% |
| 13 | 10.37% | 18.89% |

Table S7. Lymph node size measurements for injection-draining lymph nodes from tumor-bearing mice without lymph node metastasis in the pathology report.

| # of days | Mouse-C1(187) | Mouse-C2(197) |
| --- | --- | --- |
| 0 | 0.00% | 0.00% |
| 1 | -4.25% | -2.66% |
| 2 | 0.04% | -2.14% |
| 3 | -6.11% | 4.63% |
| 4 | -10.15% | 2.15% |
| 5 | -1.36% | 26.43% |
| 6 | 4.67% | 18.47% |
| 7 | 5.89% | 8.51% |
| 8 | 0.06% | 13.45% |
| 9 | -1.27% | 15.71% |
| 10 | -0.12% | 10.62% |
| 11 | 16.77% | 16.49% |
| 12 | 3.29% | 3.07% |
| 13 | 10.37% | 18.89% |

Table S8. Lymph node size measurements for contralateral lymph nodes from control mice.

There are a few limitations to this technique that would contribute to the variability in the measurements:

(1) **Imaging position and angle:** Inguinal lymph nodes are located in the groin area of mice. Since it is a creased area with loose skin on top of it, the inguinal lymph nodes need to be gently pulled to the surface of the abdominal cavity in order to be imaged clearly. Therefore there will be inconsistencies associated with this slight difference in imaging positions from day to day operations. The imaging angle is also another factor that could also give rise to some inconsistencies because SWIR-OPI provides a 2D en-face view of the lymph nodes while the lymph nodes are 3D objects that are asymmetric in shape. SWIR-OPI can only provide 2D snapshots from specific imaging angles. This problem is mitigated by taking a video of the lymph nodes while the operator gently massages the skin in different directions and therefore obtains images from slightly different angles. The results are therefore an average of multiple imaging angles and more representative of the true lymph node sizes.
(2) **Visual obstruction from tumors:** The orthotopically implanted 4T1 tumors became very large in size within two weeks. Starting from around Day 12, the primary tumors grew so large that they were usually only approximately 2 mm away from the tumor draining lymph nodes. Therefore, the operators’ ability to pull the loose skin around the mice lymph node area became very limited, and the measurements became less accurate. It is far more difficult to pull the lymph nodes from the crease in the groin area to the surface of the mouse body for imaging when the tumor is that close to a lymph node. This phenomenon could have contributed to the downward trend in size towards the end of the two-week measurement period, shown in Figure 5 in the main text.

#### 2.2 Statistical analysis

Here, we performed statistical analysis on the results that we obtained above. First, we used ANOVA analysis to prove that the tumor draining lymph nodes from metastatic mice (Table S3) are statistically different from the tumor draining lymph nodes from non-metastatic mice (Table S5). ANOVA results showed that the p value is 0.02, lower than 0.05 and that F is 6.55, greater than 4.23. Therefore, tumor draining lymph nodes from metastatic mice are statistically different in size from the tumor draining lymph nodes from non-metastatic mice.

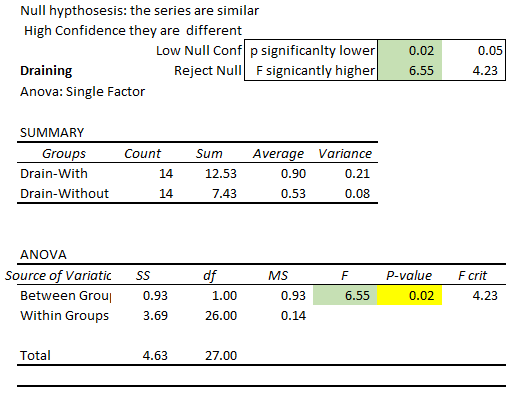

Table S9. ANOVA analysis of tumor draining lymph nodes from metastatic and non-metastatic groups

Second, to prove that the statistical difference mentioned above is not due to reasons other than lymph node metastasis, we performed another ANOVA analysis on the contralateral lymph nodes from metastatic mice versus non-metastatic mice. Results showed that the p value is 0.94, greater than 0.05 and that F is 0.01, lower than 4.23. Therefore, the sizes of contralateral lymph nodes are not statistically different in metastatic mice versus non-metastatic mice.

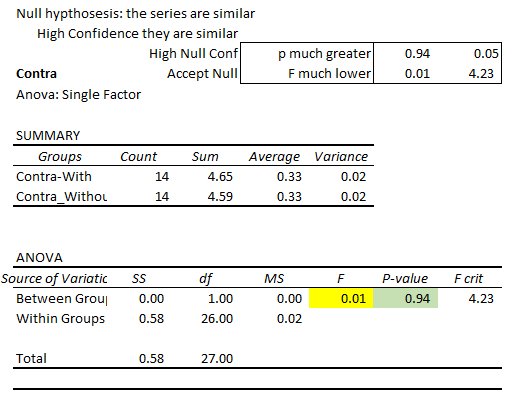

Table S10. ANOVA analysis of contralateral lymph nodes from metastatic and non-metastatic groups

Third, we investigated how early we could differentiate the tumor draining lymph nodes from metastatic mice compared to those from non-metastatic mice. As shown in Table S11, starting from Day 4, the sizes of lymph nodes are consistently significantly different, as the p values are smaller than 0.05, and the F values are greater than 4.23.

| Days | F | p-value |
| --- | --- | --- |
| 1 | 3.85 | 0.066441 |
| 2 | 6.45 | 0.021195 |
| 3 | 3.02 | 0.100249 |
| 4 | 11.02 | 0.00405 |
| 5 | 5.37 | 0.033272 |
| 6 | 10.22 | 0.033272 |
| 7 | 14.78 | 0.0013 |
| 8 | 13.51 | 0.001874 |
| 9 | 7.03 | 0.0168 |
| 10 | 9.32 | 0.007187 |
| 11 | 20.12 | 0.000326 |
| 12 | 5.79 | 0.027817 |
| 13 | 8.26 | 0.010521 |

Table S11. Day to day comparison of tumor draining lymph nodes from mice with metastatic lymph nodes or non-metastatic lymph nodes. Starting from Day 4, metastatic lymph nodes are consistently larger than non-metastatic lymph nodes.

#### 2.3 Pathology report

Here, we have included H&E histology images of lymph nodes in which we discovered metastasis (Figure S13). Two H&E slides were analyzed for every lymph node specimen. Black arrows point to tumor cells and yellow arrows point to normal cells in lymph nodes, as marked by professional rodent pathologists.

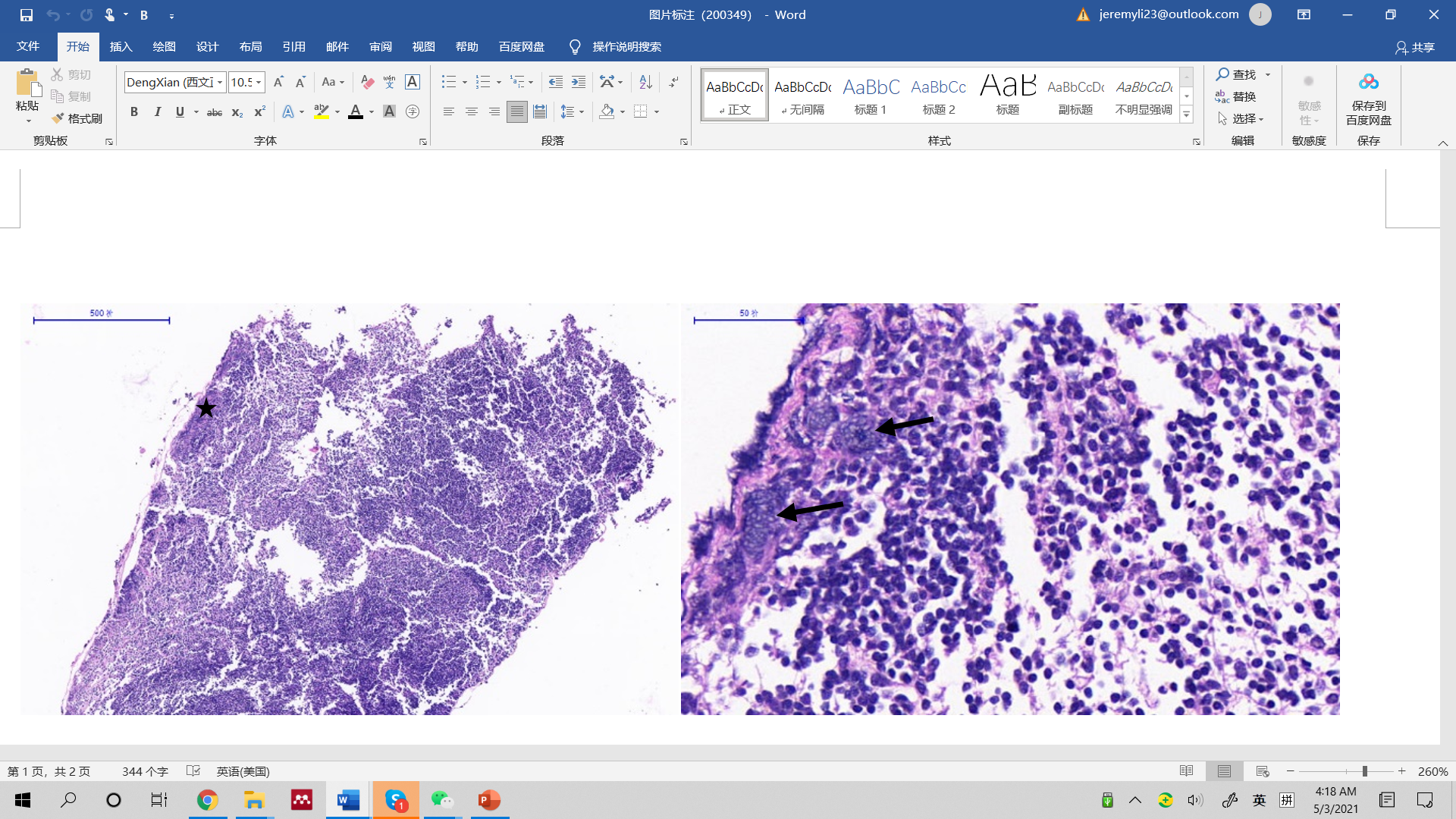

500 μm

50 μm

50 μm

500 μm

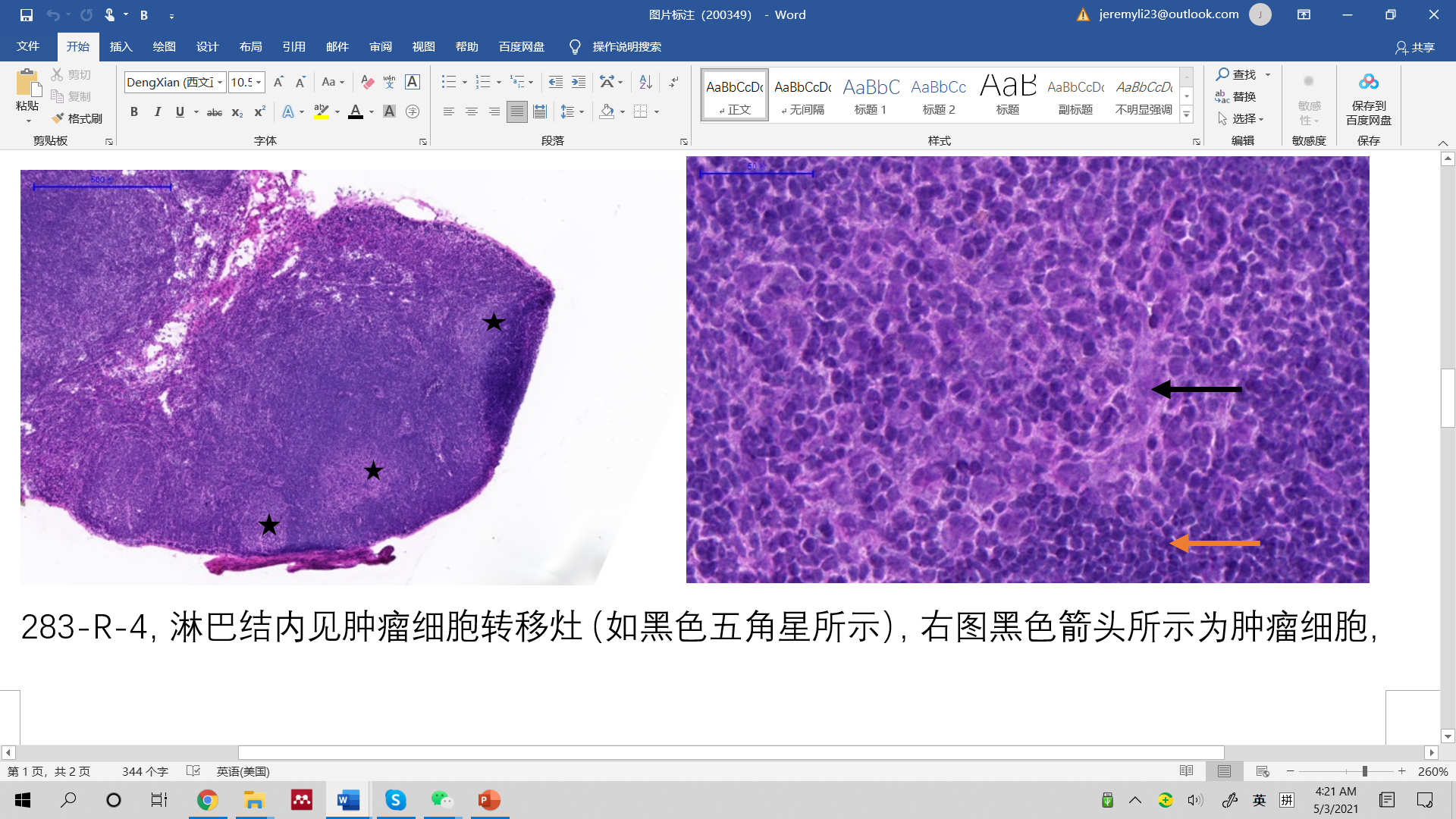

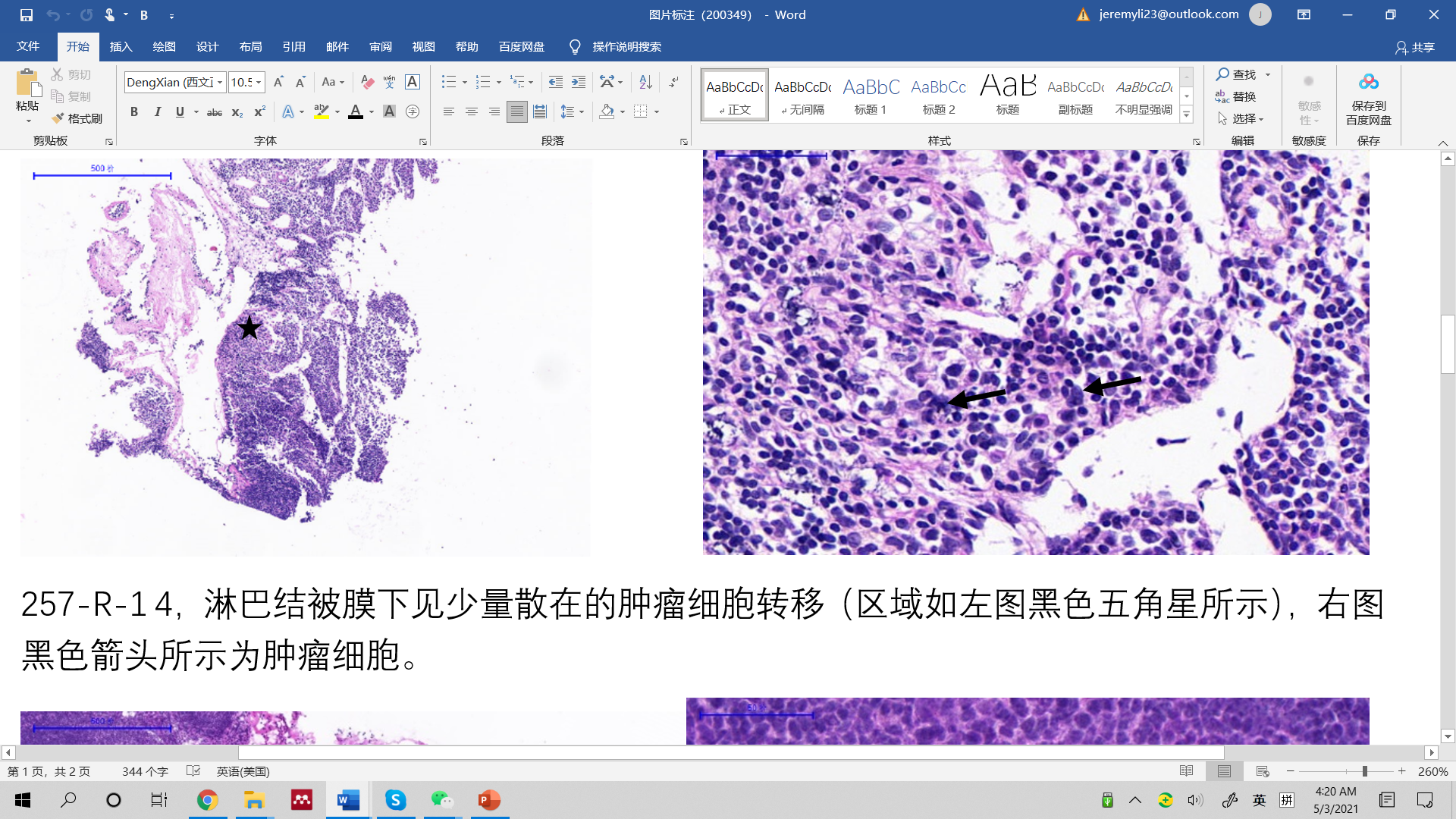

500 μm

500 μm

50 μm

50 μm

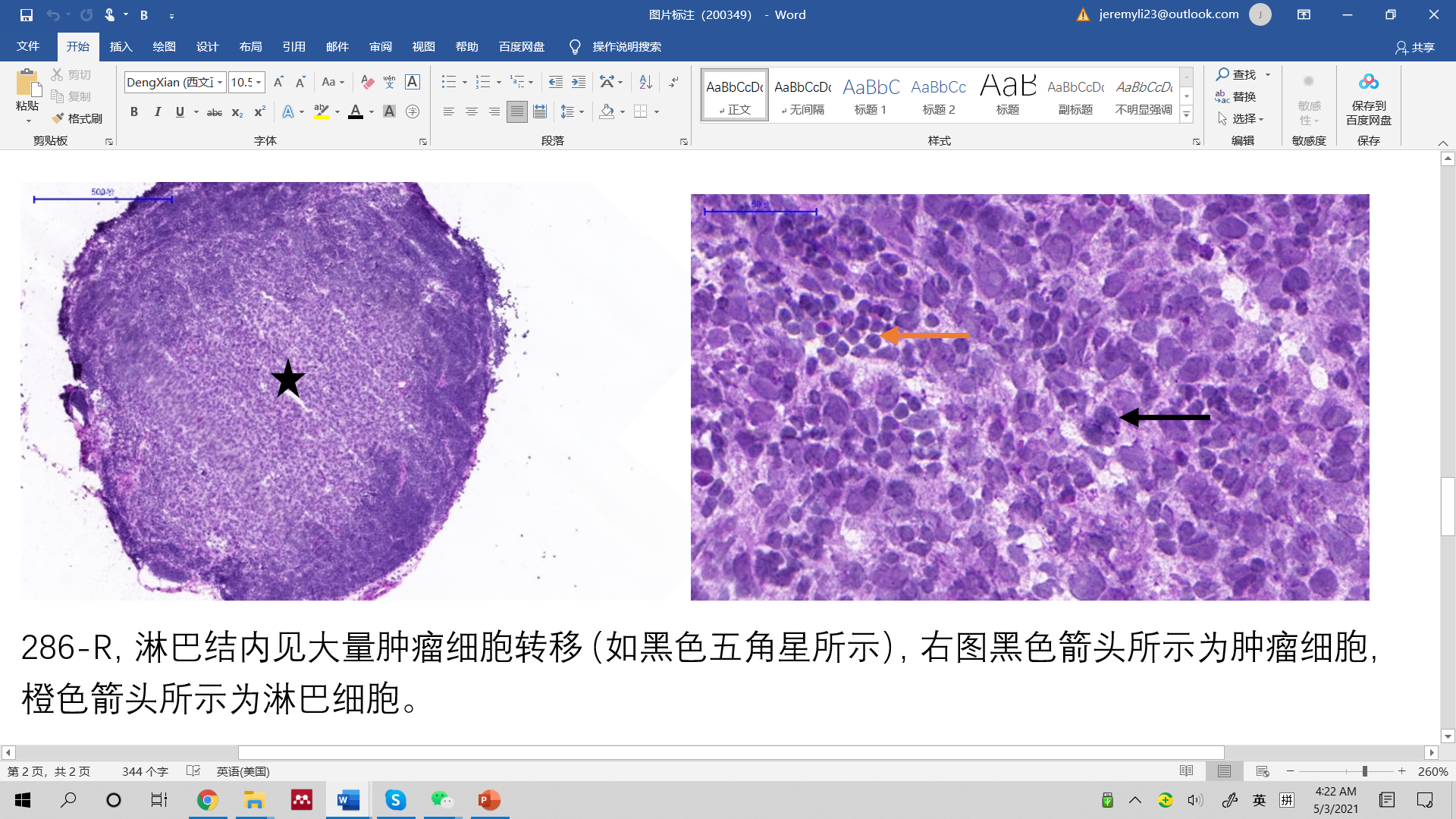

500 μm

50 μm

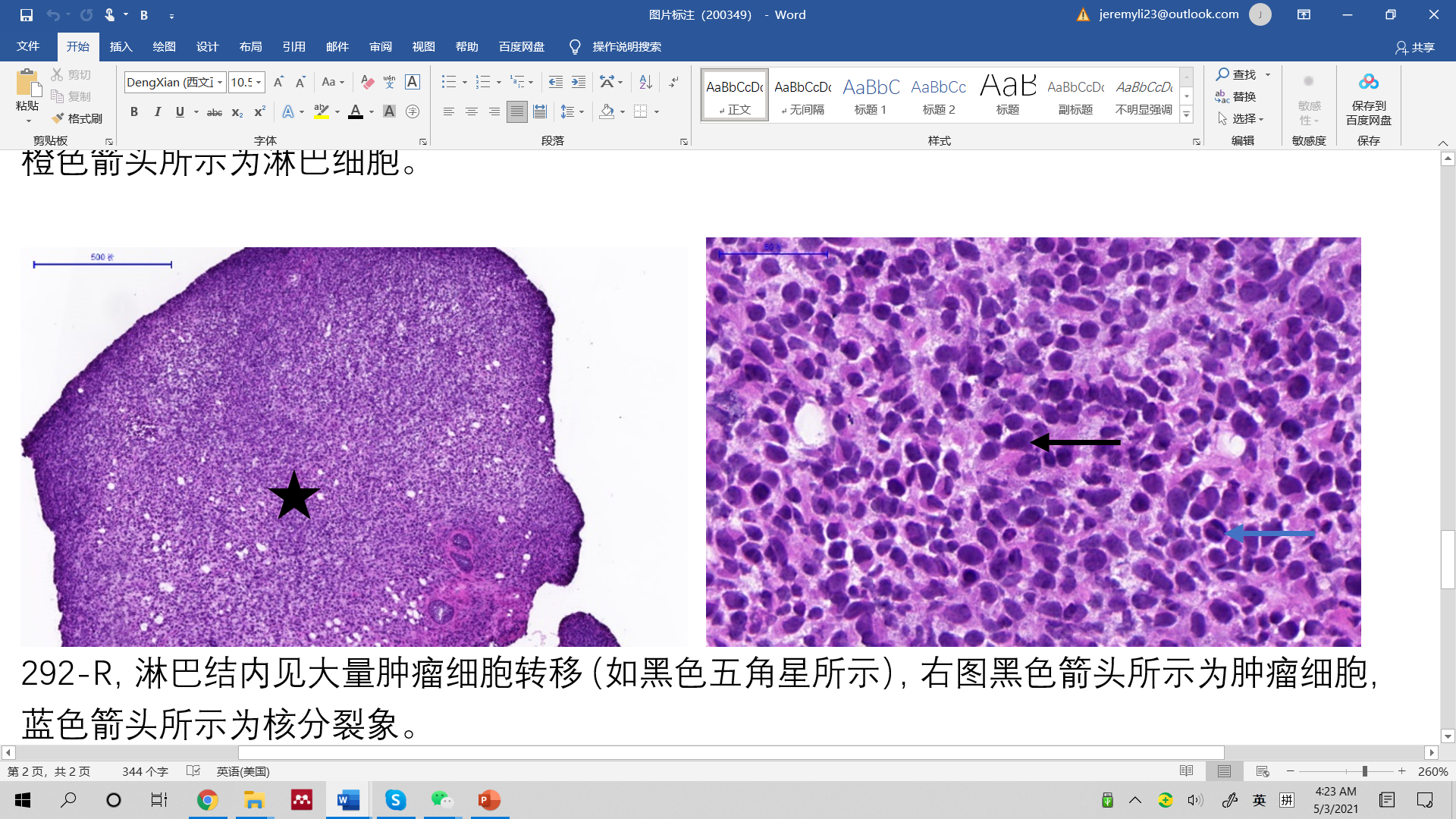

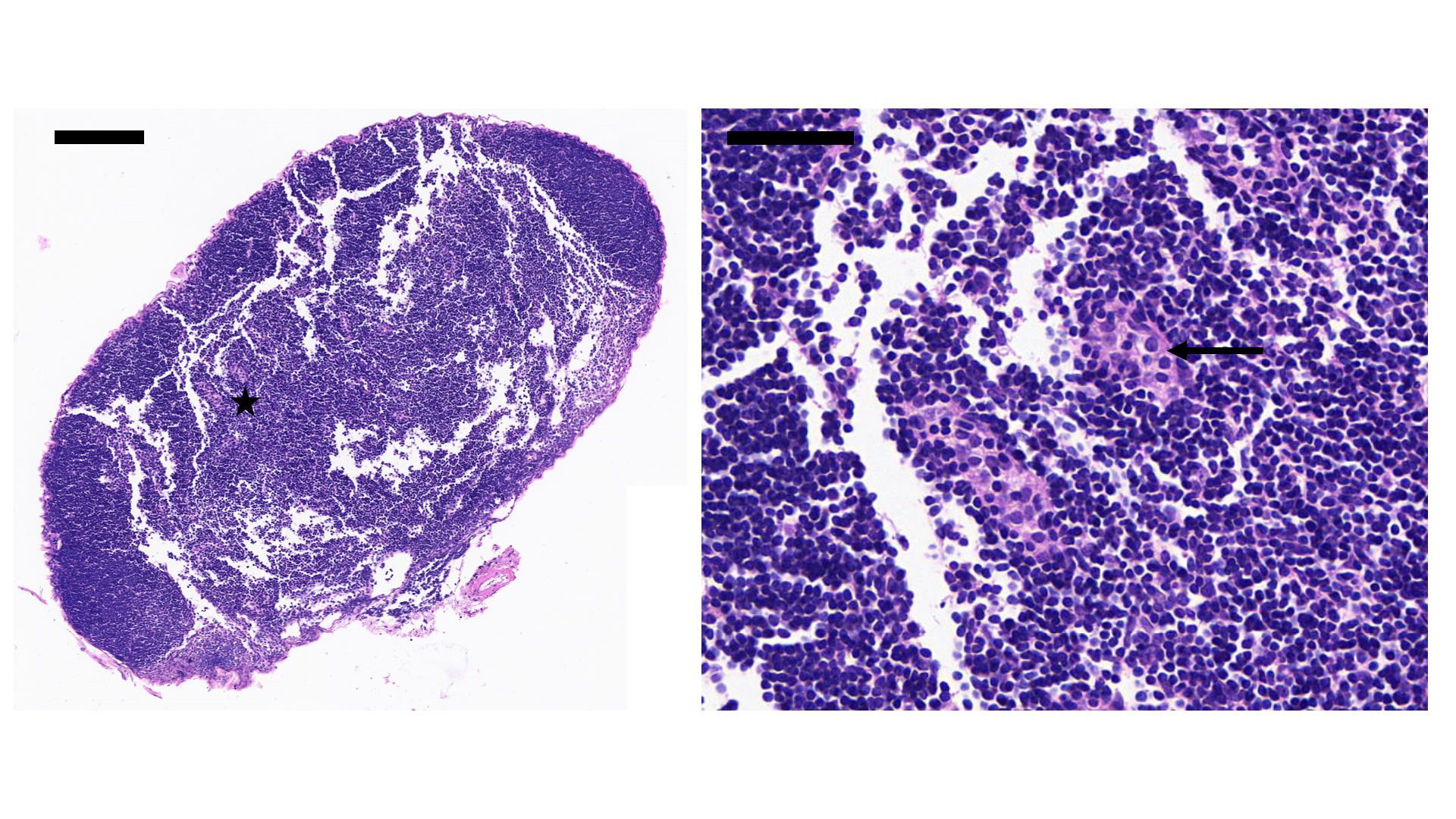

200 μm

50 μm

500 μm

50 μm

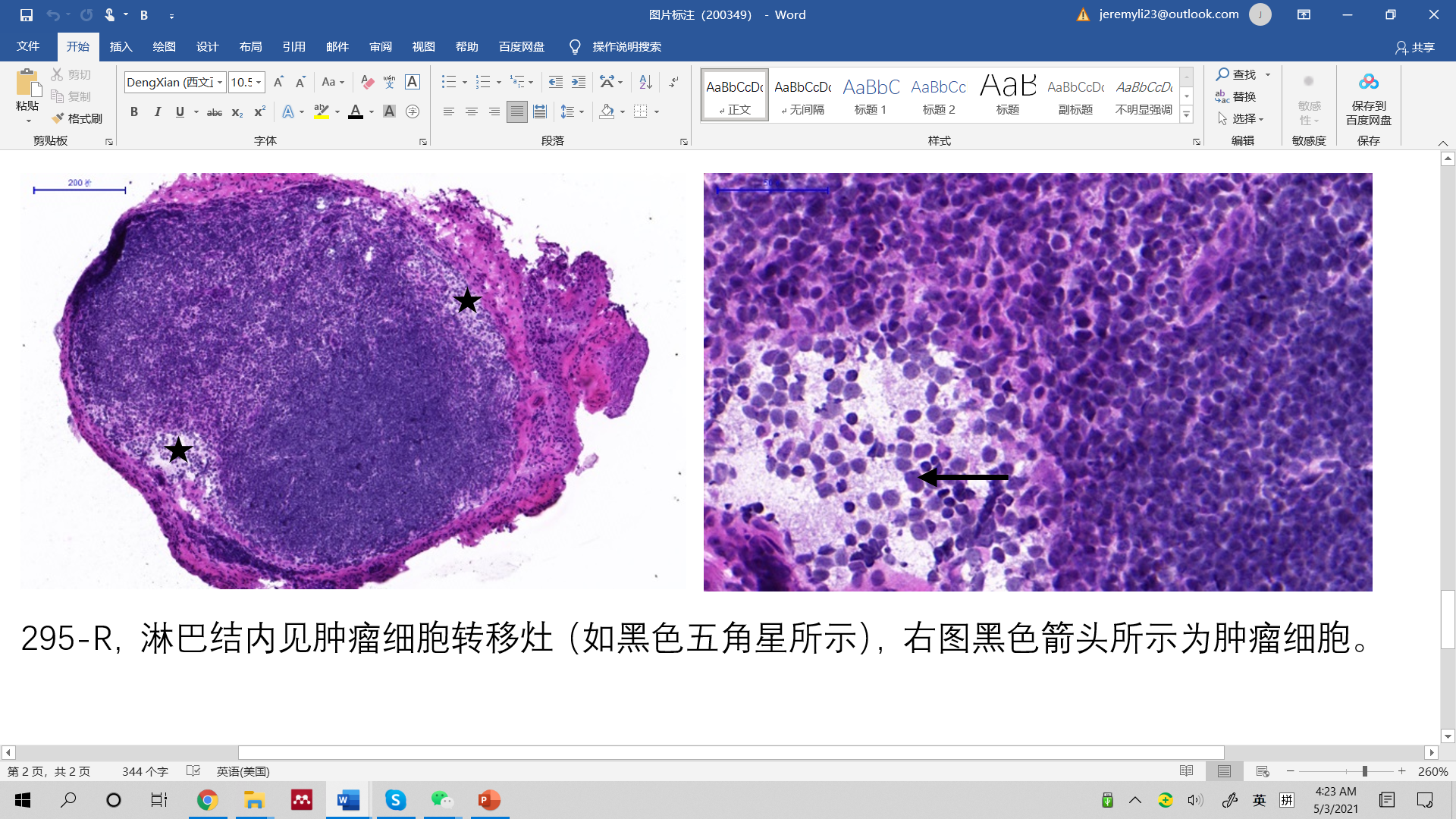

200 μm

50 μm

Figure S13. H&E scans of metastatic lymph nodes.

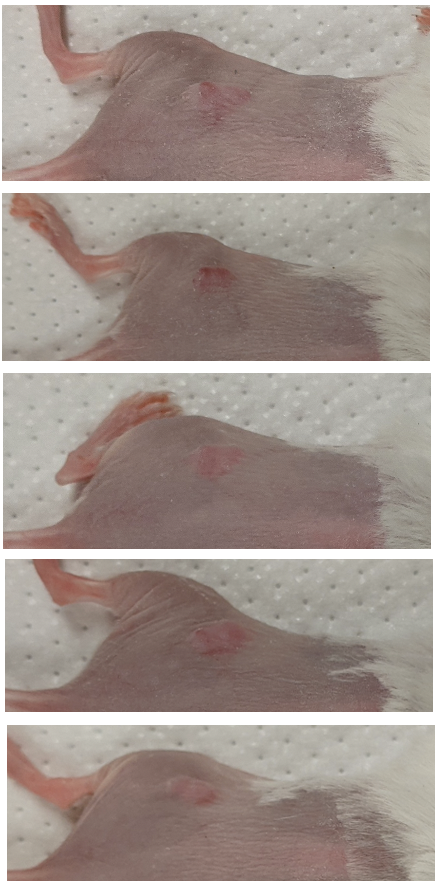

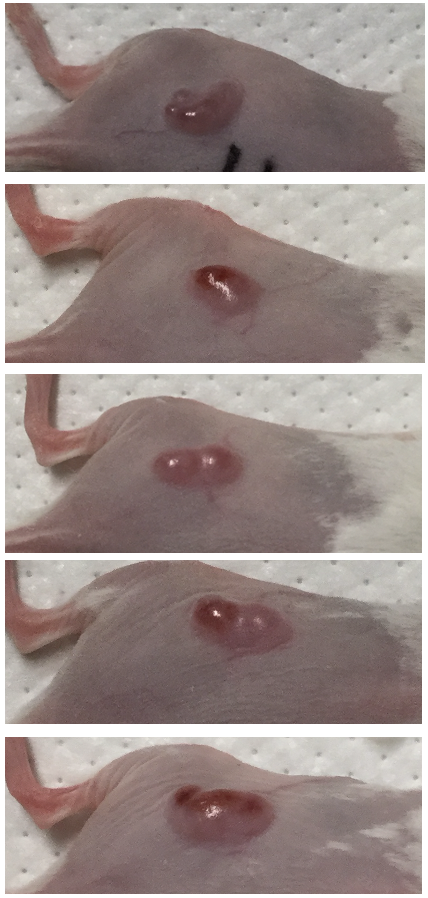

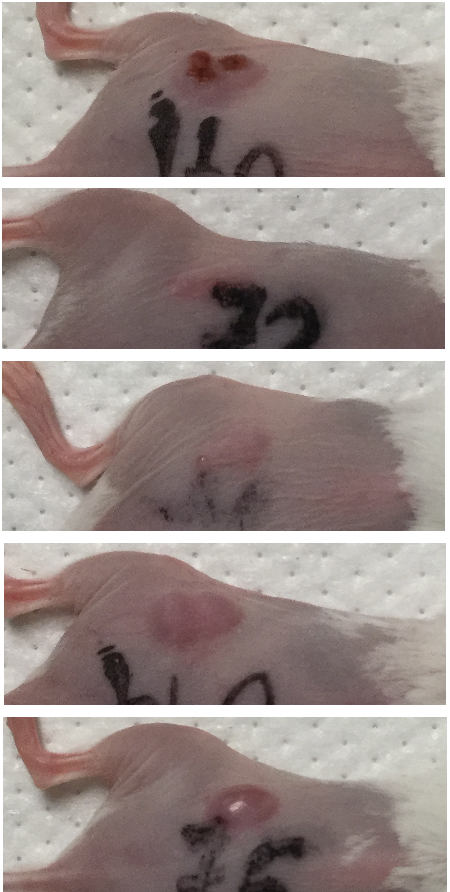

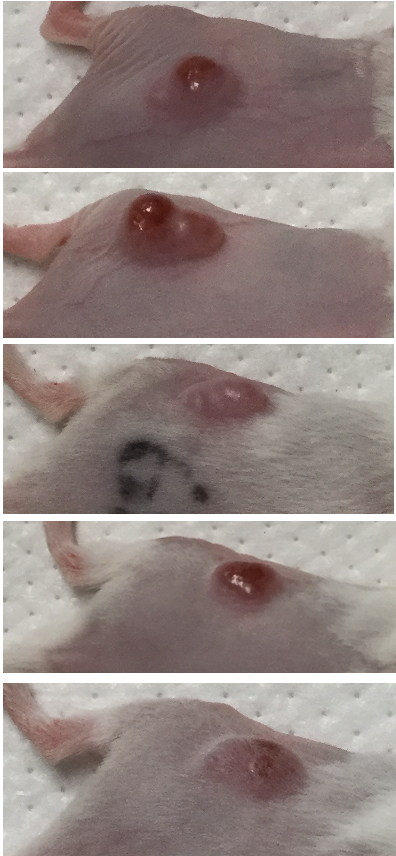

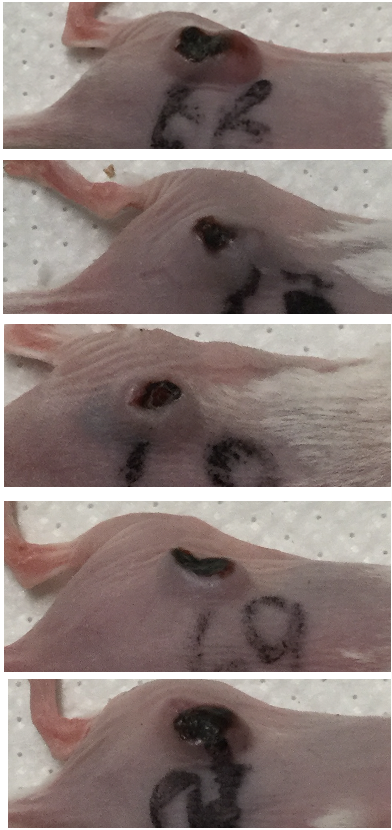

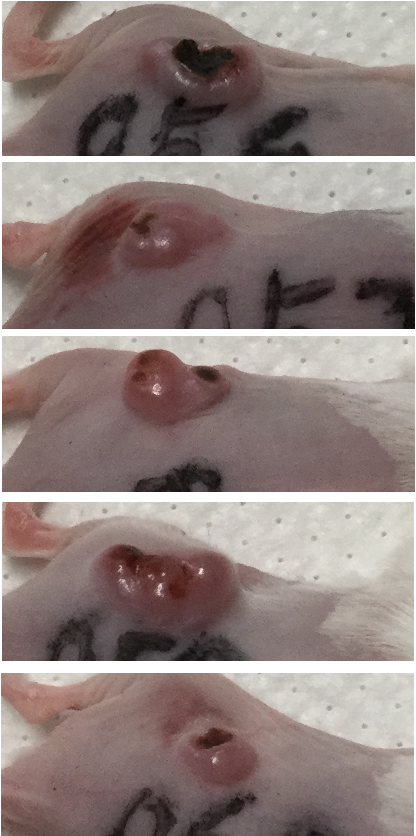

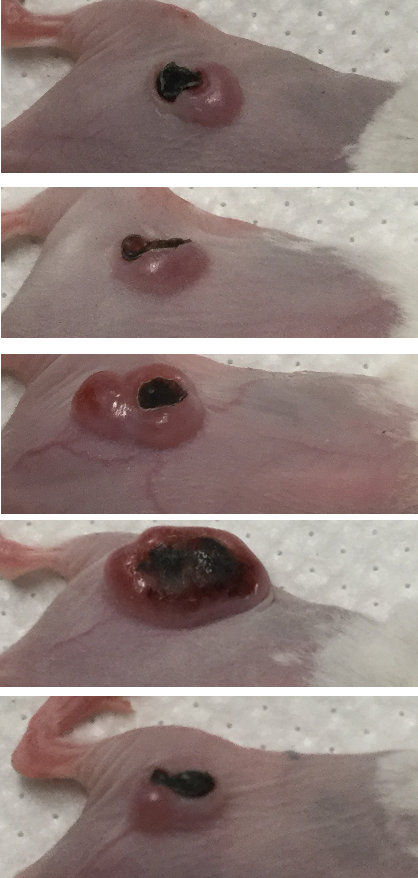

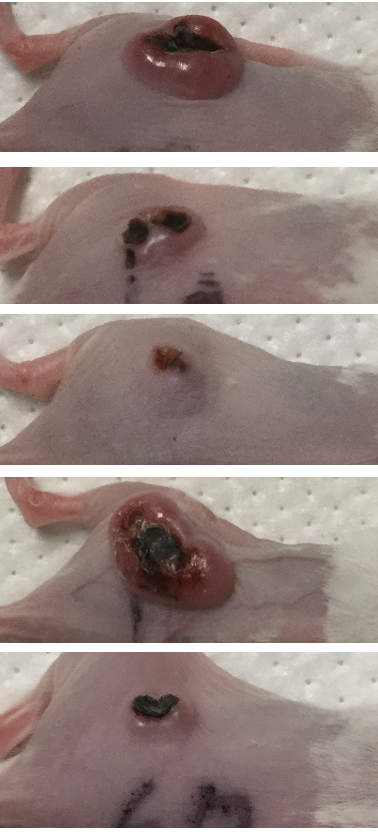

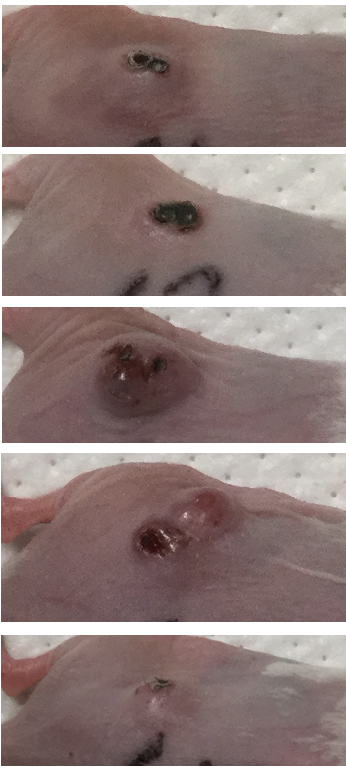

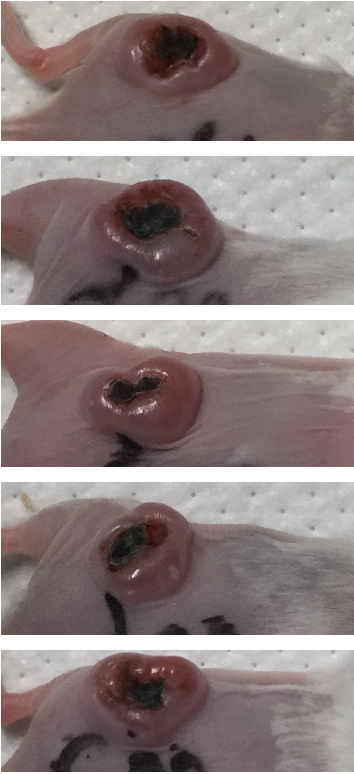

### 3. MC38 cancer cytokine correlation study

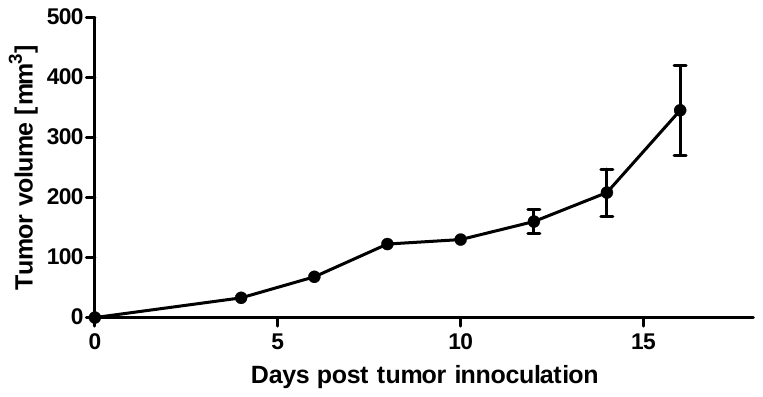

(b)

(a)

Day 5

Day 6

Day 7

Day 8

Day 9

Day 12

Day 13

Day 14

Day 16

Day 17

Here we show the growth curve along with photos of all MC38 tumors before cytokine analysis.

Figure S14. (a) MC38 tumor growth curve of Figure 5, syringe symbols represent the day of intratumoral treatment. (b) Photos of all mice (N = 5 per group) before sacrifice for tumor and lymph nodes sampling, labeled with number of days post tumor inoculation. Scale bar 1 cm.

### 4. Clinical study with human mesenteric lymph nodes

A pilot clinical study was carried out at the School of Health Science at Quinnipiac University. All study protocols were reviewed and approved by the Institutional Review Board (IRB) of Quinnipiac University.

Gross examination of lymph nodes, i.e., lymph node grossing, is one of the most common yet difficult and time-consuming procedures in anatomical pathology. Among the different cancer types, finding mesenteric lymph nodes in colectomy specimens is the most common scenario for lymph node grossing. The standard of care for lymph node grossing is manual palpation. Pathologists’ assistants, resident pathologists, or junior pathologists usually spend on average approximately 1.5 hours per sample, and inconsistencies due to human error are prevalent in hospitals around the world. As mentioned in the main text, the American Joint Cancer Committee (AJCC) requires a specific number of lymph nodes to be analyzed for specific cancer types, because the number of lymph nodes examined has been demonstrated to be correlated with patient survival. Unfortunately, human error during lymph node grossing is common and clinically consequential.^2^ Human error in this area also makes this process extremely wasteful at the moment. A large amount of adipose tissue is commonly submitted when the required numbers of lymph nodes cannot be found during lymph node grossing, creating unnecessary costs for histology materials and histo-tech and pathologist time.

Here, we have demonstrated that SWIR-OPI can be a powerful tool to improve the efficiency and accuracy of lymph node grossing. As shown in Figure 6 in the main text and in Figure S14 below, lymph nodes appear with significantly higher contrast in SWIR-OPI images than in regular images.

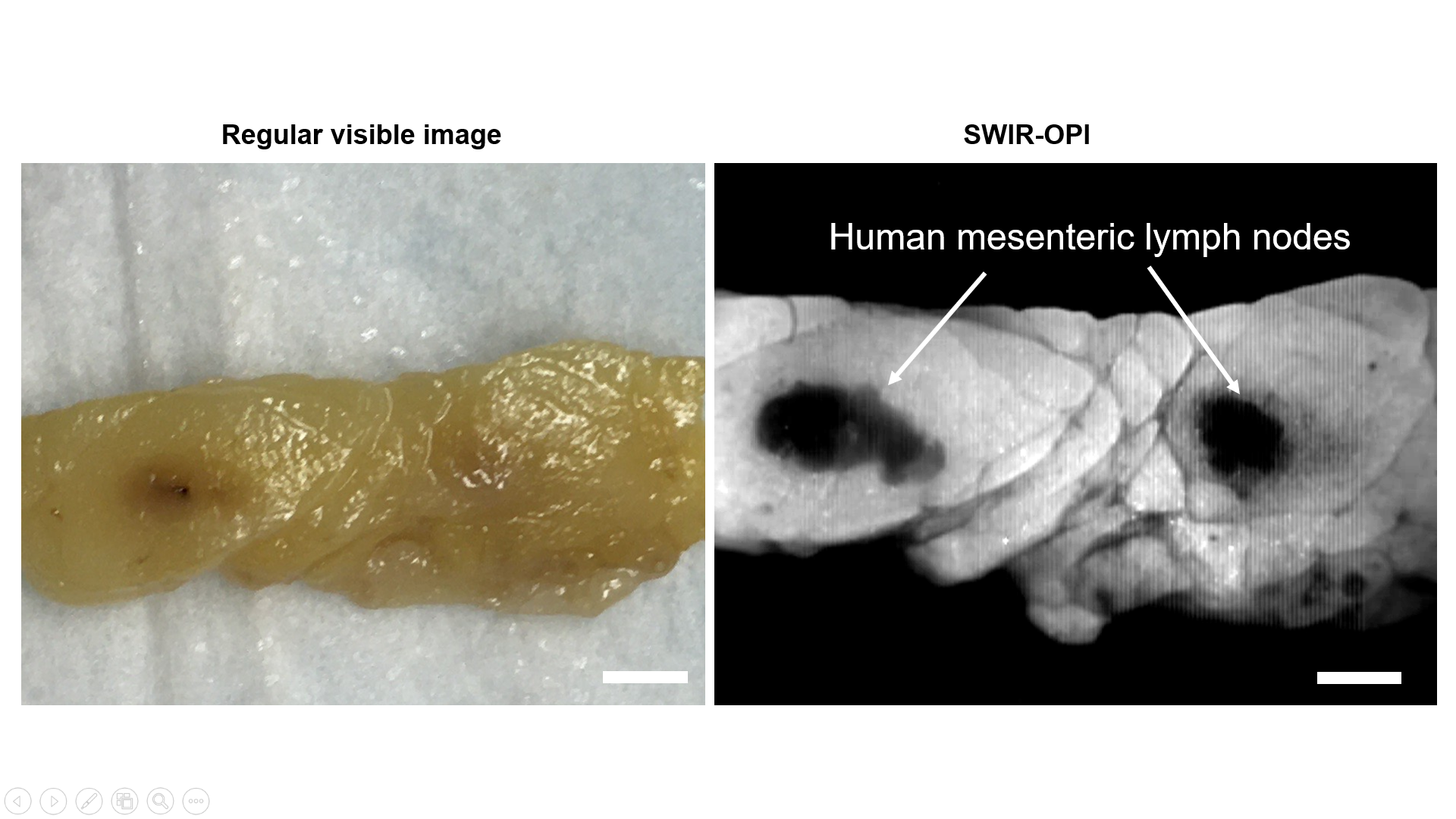

Figure S14. Human mesenteric lymph nodes under SWIR-OPI (*right*) and regular (*left*) imaging systems. Scale bars are 5 mm.

Compared to regular images, SWIR-OPI demonstrated another advantage during lymph node grossing over regular light imaging – hemorrhagic regions, which are a common distraction during lymph node grossing, do not appear in SWIR-OPI images, as shown in Figure S15. Hemorrhagic regions are dark in color and prevalent in mesentery specimens; they make the lymph nodes even more difficult to see. The reason for such contrast in regular images is not completely understood, but it is speculated to originate from scattering. Detailed analysis regarding the origin of the SWIR signal is included in Section 7 of the Supplementary Information.

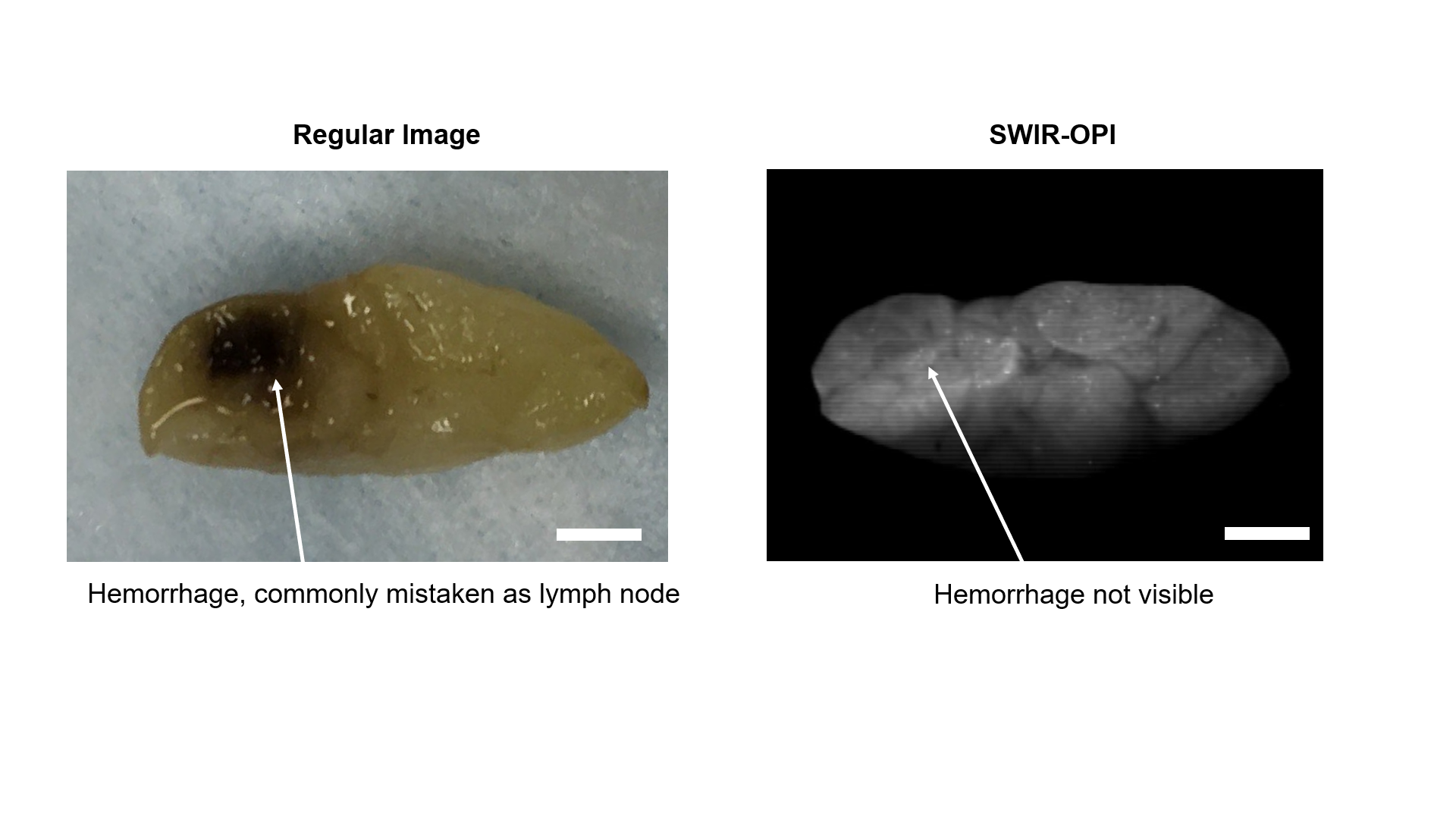

Figure S15. Hemorrhagic region in a human mesentery specimen under SWIR-OPI (*right*) and a regular imaging system (*left*). Scale bars are 5 mm.

### 5. Investigation of SWIR-OPI signal origin

SWIR-OPI is a label-free imaging technique that is based on the natural chemical contrast between different types of tissue. Regarding the origin of the SWIR-OPI signal, it can be attributed to absorption and scattering effects of SWIR light inside biological tissue. We speculate that water content is the main reason for the high contrast of lymph nodes and lymphatic vessels under SWIR-OPI.

As shown in Figure 2 in the main text, the contrast of the lymph nodes is greatest around 1500 nm. The natural optical absorption of water in the SWIR range is a broad peak between 1400 nm – 1600 nm with a maximum at 1453 nm. Water absorbs far more light in this region compared to any other wavelength in the visible and near-infrared regions. Quantitatively, the absorption coefficient of water at 1550 nm is approximately 20 cm^-1^, whereas the absorption coefficient of water at 970 nm is approximately 0.5 cm^-1^.^3^

In order to demonstrate that water content is the most important contributing factor to lymph node contrast in SWIR-OPI, we compared images of lymph nodes before and after dehydration. As shown in Figure S16(A), we excised mouse inguinal lymph nodes and first imaged under SWIR-OPI. We then dehydrated the tissue specimen by placing the specimen inside a small (1 cm × 1 cm) Ziploc^TM^ bag with silicone silica gel packets (Interteck Packaging) for 48 hours. As shown in Figure S16(B), SWIR-OPI can no longer detect a contrast between lymph nodes and the surrounding fat tissue after drying.

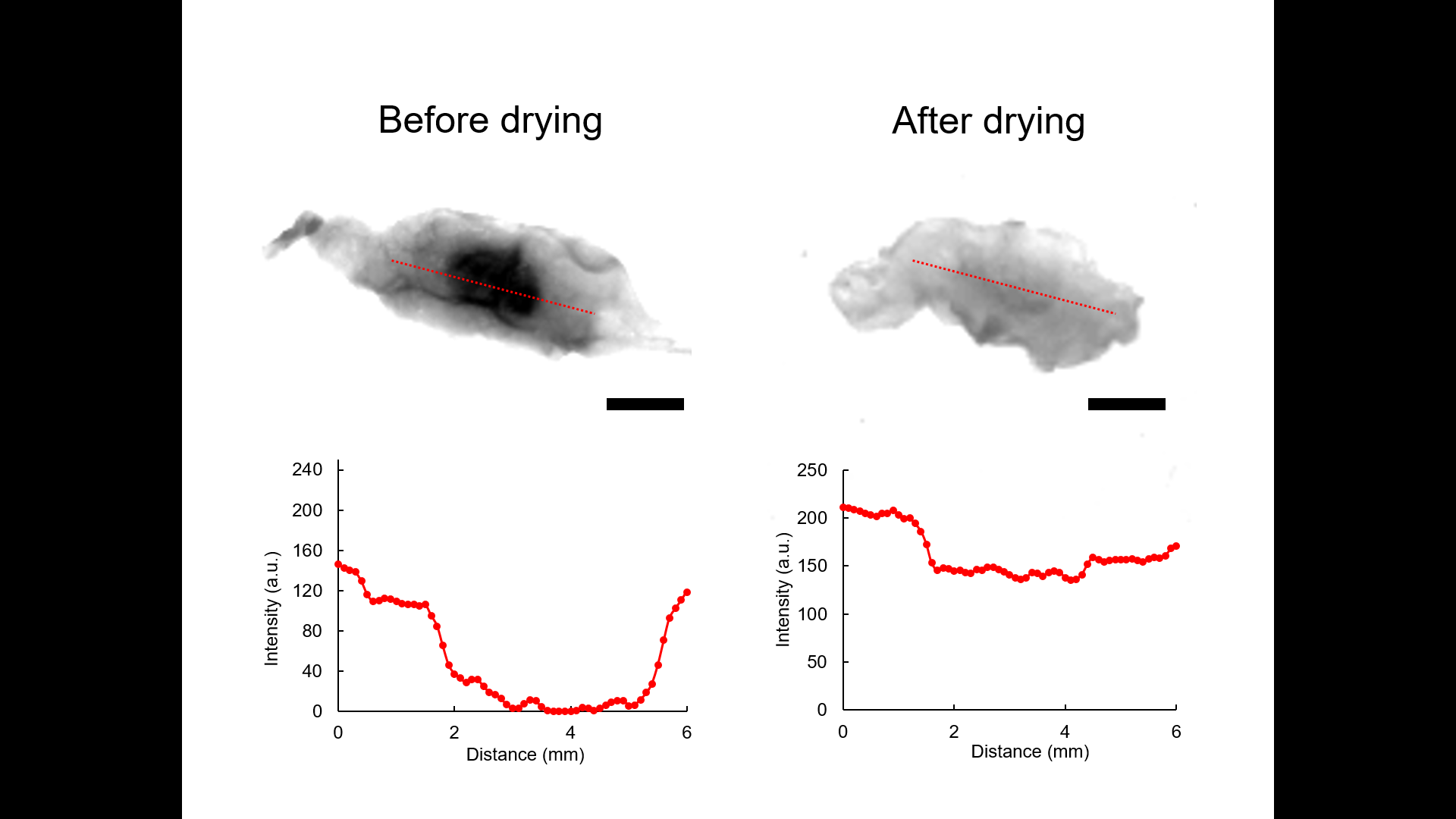

B

A

Figure S16. SWIR-OPI imaging of an *ex vivo* mouse inguinal lymph node specimen before and after drying.

The strength of SWIR-OPI signal, i.e., contrast between features of interest and the surrounding background tissue, is not a simple function of differences in water content. Blood and blood vessels, which are notably high in water content, do not appear dark under SWIR-OPI compared to the surrounding fat tissue, as shown Figure S18. This interesting observation of lack of contrast can be explained by the scattering effects from red blood cells, which constitute 40% of the whole blood volume. To test this hypothesis, we imaged capillary tubes filled with whole blood, blood serum, and water side by side. Blood serum was obtained by centrifuging a whole blood sample at 4000 rpm for 15 minutes. Blood serum does not contain blood cells that are in whole blood. As shown in Figure S17, water appears completely dark under SWIR-OPI at 1550 nm, as expected. Whole blood appears significantly lighter under the same conditions, whereas blood serum appears significantly darker compared to whole blood and very comparable to water. This experiment demonstrates that blood cells in whole blood are the reason for the lack of contrast of blood vessels compared to the surrounding fat tissue, as shown in Figure S19.

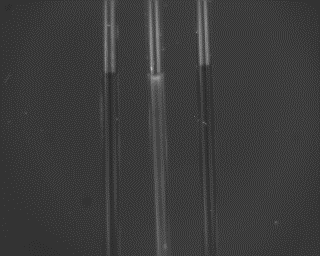

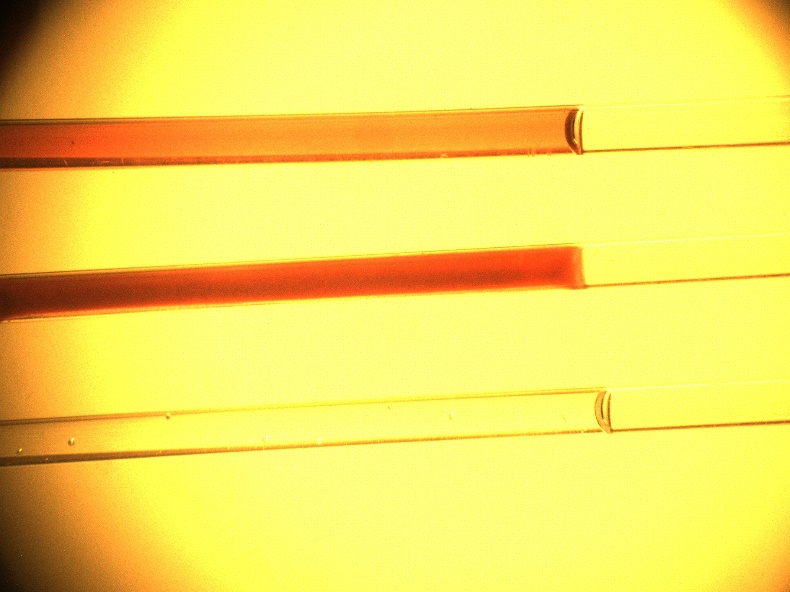

(A)

(B)

water

blood

serum

water

blood

serum

Figure S17. Comparison of water, blood, and serum under regular visible light illumination and SWIR-OPI. Scale bars: 2 mm

Similar to the natural contrast between lymph nodes and surrounding fat, we also observed other natural sources of contrast that are highly visible under SWIR-OPI. For example, skeletal muscles (Figure S18) and mammary ducts (Figure S19) appear dark under SWIR-OPI. The reasons for this contrast are mainly attributed to their high water content. There could be many other natural anatomical sources of contrast under SWIR-OPI that are yet to be explored; however, this possibility does also present the challenge of how to distinguish different tissues that appear similar under SWIR-OPI. For example, mammary ducts in Figure S19 could be easily mistaken as lymphatic vessels. Therefore, SWIR-OPI should only be used for visualizing and measuring lymph nodes and lymphatic vessels when the anatomical locations and structural features are already known to the users. It should not be used as a lymph node or lymphatic vessel specific tool to explore anatomical locations that are not well-defined or well-known to the users. In other words, SWIR-OPI as a label-free imaging modality should be used in a similar manner compared to ultrasound, CT, or MRI, where certain level of familiarity of the anatomical locations is expected while using the technique. It is not a lymphatic structure targeted technique where one could expect all the signals to be originated from lymphatic structures.

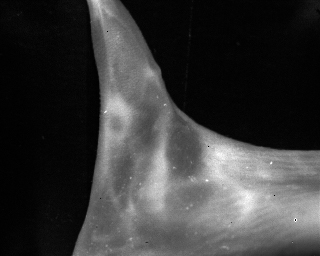

Popliteal lymph node

Skeletal muscles

Figure S18. SWIR-OPI image of mouse lower limb. Scale bar is 2 cm.

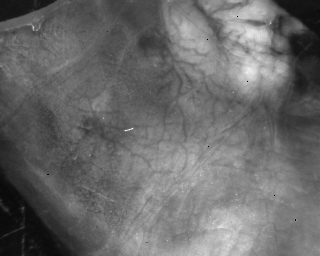

blood vessel

mammary duct

(A)

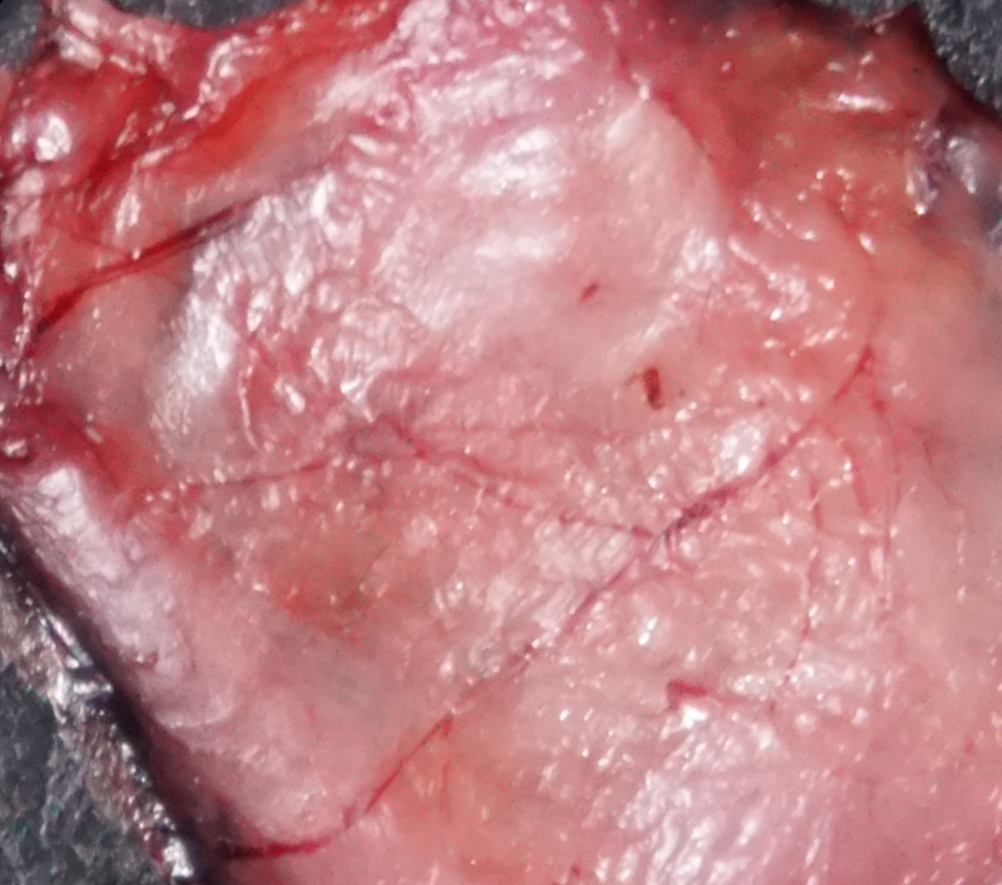

blood vessel

(B)

nipple

nipple

Figure S19. SWIR-OPI image (*left*) and regular image (*right*) of the inside of a mouse skin flap in the lower abdomen. Nipples and mammary ducts can be clearly seen under SWIR-OPI, but are not visible in the regular image. Capillary blood vessels, on the other hand, are not visible under SWIR-OPI, but can be clearly seen in the regular image. Scale bars are 2 cm.

### Appendix: Deep Learning Source Code

#### 1. U-Net architecture

Figure S20. The U-Net architecture for segmenting lymph node structures from SWIR-OPI images. Scale bars are 2 mm.

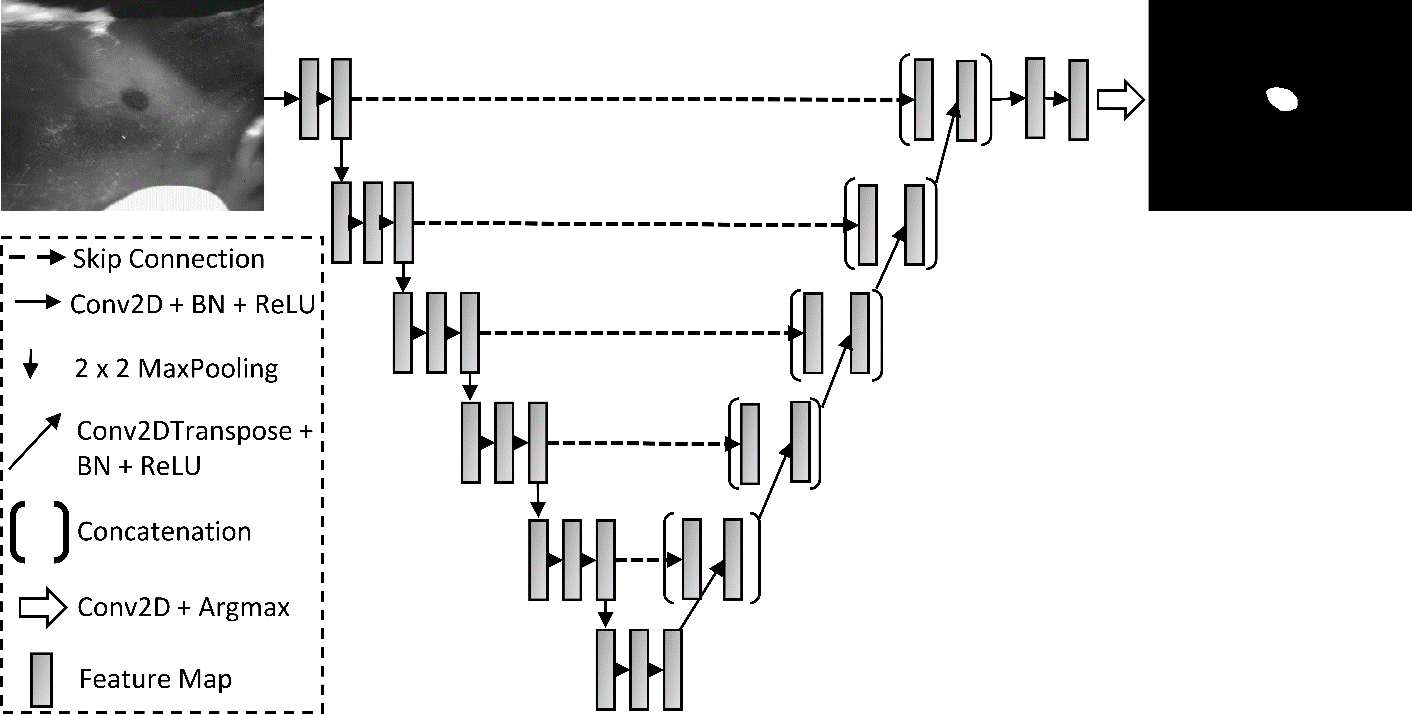

#### 2. Source code for training:

from __future__ import absolute_import

from __future__ import division

from __future__ import print_function

import argparse

import sys

import math

import numpy as np

import os

import random

import time

import tensorflow as tf

import tensorflow.contrib.slim as slim

from tensorflow.contrib.slim.python.slim.nets import resnet_v2

from tensorflow.python.ops import control_flow_ops

import skimage

from skimage.io import imread

from skimage.io import imsave

from skimage.transform import rotate

from skimage.transform import rescale

from operator import itemgetter

from skimage.morphology import label

from scipy.ndimage.morphology import binary_dilation

FLAGS = None

imageType = 1

ous = 192

ins = 192

interv = 255

nclass =2

nclass4= 2

batch_size = 8

test_batch_size = 0

#homedir = '/home/lyang5/'

homedir = '/afs/crc.nd.edu/user/y/yzhang29/Mouse_lymph_node/'

logdir = homedir + 'tftrain'

resultdir = homedir + 'tfsave_unet'

imgdir = '/afs/crc.nd.edu/user/y/yzhang29/Mouse_lymph_node/'

def load_data(file_dir):

train_imgs = []

label_imgs = []

count = 0

for file in os.listdir(os.path.join(file_dir,'data/unlabeled/')):

if file.endswith(".png"):

tI=imread(os.path.join(file_dir,'data/unlabeled/',file))

#if (len(tI.shape)==3):

\# tI=tI[:,:,0]

tI=skimage.img_as_float(tI)

#tI=rescale(tI,0.5,order=0)

if (len(tI.shape)==2):

tI=np.expand_dims(tI,axis=2)

train_imgs.append(tI)

lI=imread(os.path.join(file_dir,'data/mask727/',file))

#lI=skimage.img_as_float(lI)

print (np.unique(lI))

if (len(lI.shape)==3):

lI=lI[:,:,0]

label_imgs.append(lI)

for file in os.listdir(os.path.join(file_dir,'data/train/')):

if file.endswith(".png"):

tI=imread(os.path.join(file_dir,'data/train/',file))

#if (len(tI.shape)==3):

\# tI=tI[:,:,0]

tI=skimage.img_as_float(tI)

#tI=rescale(tI,0.5,order=0)

if (len(tI.shape)==2):

tI=np.expand_dims(tI,axis=2)

train_imgs.append(tI)

lI=imread(os.path.join(file_dir,'data/mask/',file))

#lI=skimage.img_as_float(lI)

print (np.unique(lI))

if (len(lI.shape)==3):

lI=lI[:,:,0]

label_imgs.append(lI)

return train_imgs, label_imgs

def load_test(file_dir):

test_imgs = []

for file in os.listdir(os.path.join(file_dir,'raw/')):

if file.endswith(".png"):

tI=imread(os.path.join(file_dir,'raw/',file))

tI=skimage.img_as_float(tI)

#tI=rescale(tI,0.5,order=0)

if (len(tI.shape)==2):

tI=np.expand_dims(tI,axis=2)

test_imgs.append(tI)

return test_imgs

def get_image(train_imgs,label_imgs):

i_idx = random.randint(0, len(train_imgs)-1)

big_tI = train_imgs[i_idx]

big_lI = label_imgs[i_idx]

xs = big_tI.shape[0]

ys = big_tI.shape[1]

stx = random.randint(0,xs-ins)

sty = random.randint(0,ys-ins)

train_sample = np.zeros((ins,ins,imageType))

label_sample = np.zeros((ins,ins))

train_sample = big_tI[stx:stx+ins,sty:sty+ins,:]

label_sample = big_lI[stx:stx+ins,sty:sty+ins]

nrotate = random.randint(0, 3)

train_sample = np.rot90(train_sample, nrotate)

#print (np.unique(label_sample))

label_sample = np.rot90(label_sample, nrotate).astype('uint8')

#print (np.unique(label_sample))

#asd

nflip = random.randint(0, 1)

if nflip:

#print('flip')

train_sample = np.fliplr(train_sample)

label_sample = np.fliplr(label_sample)

return train_sample, label_sample

def get_image_balanced_pos(train_imgs,label_imgs):

i_idx = random.randint(0, len(train_imgs)-1)

big_tI = train_imgs[i_idx]

big_lI = label_imgs[i_idx]

xs = big_tI.shape[0]

ys = big_tI.shape[1]

flag=1

print ('get_pos:')

while flag==1:

stx = random.randint(0,xs-ins)

sty = random.randint(0,ys-ins)

train_sample = np.zeros((ins,ins,imageType))

label_sample = np.zeros((ins,ins))

train_sample = big_tI[stx:stx+ins,sty:sty+ins,:]

label_sample = big_lI[stx:stx+ins,sty:sty+ins]

#print (label_sample[int(ins/2),int(ins/2)])

if label_sample[int(ins/2),int(ins/2)]==40.0:

flag==0

nrotate = random.randint(0, 3)

train_sample = np.rot90(train_sample, nrotate)

#print (np.unique(label_sample))

label_sample = np.rot90(label_sample, nrotate).astype('uint8')

#print (np.unique(label_sample))

#asd

nflip = random.randint(0, 1)

if nflip:

#print('flip')

train_sample = np.fliplr(train_sample)

label_sample = np.fliplr(label_sample)

return train_sample, label_sample

def get_image_balanced_neg(train_imgs,label_imgs):

i_idx = random.randint(0, len(train_imgs)-1)

big_tI = train_imgs[i_idx]

big_lI = label_imgs[i_idx]

xs = big_tI.shape[0]

ys = big_tI.shape[1]

flag=1

print ('get_neg:')

while flag==1:

stx = random.randint(0,xs-ins)

sty = random.randint(0,ys-ins)

train_sample = np.zeros((ins,ins,imageType))

label_sample = np.zeros((ins,ins))

train_sample = big_tI[stx:stx+ins,sty:sty+ins,:]

label_sample = big_lI[stx:stx+ins,sty:sty+ins]

if label_sample[int(ins/2),int(ins/2)]!=40.0:

flag==0

nrotate = random.randint(0, 3)

train_sample = np.rot90(train_sample, nrotate)

#print (np.unique(label_sample))

label_sample = np.rot90(label_sample, nrotate).astype('uint8')

#print (np.unique(label_sample))

#asd

nflip = random.randint(0, 1)

if nflip:

#print('flip')

train_sample = np.fliplr(train_sample)

label_sample = np.fliplr(label_sample)

return train_sample, label_sample

def get_batch(train_imgs,label_imgs,batch_size):

train_samples = np.zeros((batch_size,ins,ins,imageType))

label_samples = np.zeros((batch_size,ins,ins))

for i in range(int(batch_size)):

train_sample, label_sample = get_image(train_imgs,label_imgs)

train_samples[i,:,:,:] = train_sample

label_samples[i,:,:] = label_sample/interv

return train_samples, label_samples.astype('int32')

def get_test_batch(test_imgs,batch_size):

test_samples = np.zeros((batch_size,ins,ins,imageType))

label_samples = np.zeros((batch_size,ins,ins))

for i in range(batch_size):

test_sample, label_sample = get_test_image(test_imgs)

test_samples[i,:,:,:] = test_sample

label_samples[i,:,:] = 2

return test_samples, label_samples.astype('int32')

import tensorflow as tf

import math

def weight_variable(shape,stdv):

initial = tf.get_variable("weights", shape=shape, initializer=tf.random_normal_initializer(stddev=stdv))

return initial

def bias_variable(shape):

initial = tf.get_variable("bias", shape=shape, initializer=tf.constant_initializer(value=0.0))

return initial

def conv_(x,k,nin,nout,phase,s=1,d=1):

stdv = math.sqrt(2/(nin*k*k))

return tf.layers.conv2d(x, nout, k, strides=[s,s], dilation_rate=[d,d], padding='same',

kernel_initializer = tf.random_normal_initializer(stddev=stdv),

use_bias=False)

def convnn_(x,k,s,nin,nout,phase):

print('convnn')

stdv = math.sqrt(1/(nin*k*k))

return tf.layers.conv2d(x, nout, k, strides=[s,s], padding='same',

kernel_initializer = tf.random_uniform_initializer(minval=-stdv, maxval=stdv,dtype=tf.float32),

use_bias=False)

def bn_(x,phase):

bn_result = tf.layers.batch_normalization(x, momentum=0.9, epsilon = 1e-5,

gamma_initializer = tf.random_uniform_initializer(minval=0.0, maxval=1.0,dtype=tf.float32),

training = phase)

return bn_result

def relu_(x):

return tf.nn.relu(x)

def cbr_(x,k,nin,nout,phase,s=1,d=1):

x_conv = conv_(x,k,nin,nout,phase,s,d)

x_bn = bn_(x_conv,phase)

x_relu = relu_(x_bn)

return x_relu

def bottleneck(x, nin, nout, phase, s=1, d=1):

if (nin != nout) or (s != 1):

print('conv_skip')

with tf.variable_scope('skip'):

skip_conv = conv_(x,1,nin,nout,phase,s=s)

skip = bn_(skip_conv,phase)

else:

skip = x

with tf.variable_scope('conv1'):

c1_conv = conv_(x,3,nin,nout,phase,s=s,d=d)

c1_bn = bn_(c1_conv,phase)

c1_relu = relu_(c1_bn)

with tf.variable_scope('conv2'):

c2_conv = conv_(c1_relu,3,nout,nout,phase,d=d)

c2_bn = bn_(c2_conv,phase)

out = skip + c2_bn

out_relu = relu_(out)

return out_relu

def stack(x, nin, nout, nblock, phase, s=1, d=1, new_level=True):

for i in range(nblock):

with tf.variable_scope('block%d' % (i)):

if i==0:

x = bottleneck(x,nin,nout,phase,s=s,d=d)

else:

x = bottleneck(x,nout,nout,phase,d=d)

return x

def max_pool_2x2(x):

print('new pool')

return tf.layers.max_pooling2d(x,[2,2],[2,2])

def avg_pool_2x2(x):

print('avg')

return tf.nn.avg_pool(x, ksize=[1, 2, 2, 1],

strides=[1, 2, 2, 1], padding='SAME')

def show(x,k):

x_sm = tf.nn.softmax(x)

x_sl = tf.unstack(x_sm,num=nclass,axis=-1)[k]

return tf.expand_dims(x_sl,-1)

def deconv_(x, nin, nout, phase):

stdv = math.sqrt(2/(nin*2*2))

conv_r = tf.layers.conv2d_transpose(x, nout, 4, strides=[2,2], padding='same',

kernel_initializer = tf.random_normal_initializer(stddev=stdv),

use_bias=False)

bn_r = bn_(conv_r,phase)

relu_r = relu_(bn_r)

return relu_r

def deconv1_(x, nin, num_up, phase):

cur_in = nin

cur_out=(2**(num_up-1))

for i in range(num_up):

with tf.variable_scope('up%d' % (i)):

x = deconv_(x,cur_in,cur_out,phase)

cur_in = cur_out

cur_out = math.floor(cur_out/2)

return x

def deconvn_(x, nin, nout, num_up, phase):

cur_in = nin

cur_out=((2**(num_up-1))*nout)

for i in range(num_up):

with tf.variable_scope('up%d' % (i)):

x = deconv_(x,cur_in,cur_out,phase)

cur_in = cur_out

cur_out = math.floor(cur_out/2)

return x

def deconv2_(x, nin, num_up, phase):

cur_in = nin

cur_out=(2**(num_up))

for i in range(num_up):

with tf.variable_scope('up%d' % (i)):

x = deconv_(x,cur_in,cur_out,phase)

cur_in = cur_out

cur_out = math.floor(cur_out/2)

return x

def model(input_img,phase,nc,id,gpu):

with tf.device('/gpu:%d' % (gpu)):

with tf.variable_scope('model%d' % (id)):

with tf.variable_scope('scale1'):

with tf.variable_scope('conv1'):

c1_1_conv = conv_(input_img, 3,imageType,nc,phase)

c1_1_bn = bn_(c1_1_conv, phase)

c1_1_relu = relu_(c1_1_bn)

with tf.variable_scope('conv2'):

c1_2_conv = conv_(c1_1_relu,3,nc,2*nc,phase)

c1_2_bn = bn_(c1_2_conv, phase)

c1_2_relu = relu_(c1_2_bn)

c1_pool = max_pool_2x2(c1_2_relu)

with tf.variable_scope('scale2'):

with tf.variable_scope('conv1'):

c2_1_conv = conv_(c1_pool,3,2*nc,2*nc,phase)

c2_1_bn = bn_(c2_1_conv, phase)

c2_1_relu = relu_(c2_1_bn)

with tf.variable_scope('conv2'):

c2_2_conv = conv_(c2_1_relu,3,2*nc,4*nc,phase)

c2_2_bn = bn_(c2_2_conv, phase)

c2_2_relu = relu_(c2_2_bn)

c2_pool = max_pool_2x2(c2_2_relu)

with tf.variable_scope('scale3'):

with tf.variable_scope('conv1'):

c3_1_conv = conv_(c2_pool,3,4*nc,4*nc,phase)

c3_1_bn = bn_(c3_1_conv, phase)

c3_1_relu = relu_(c3_1_bn)

with tf.variable_scope('conv2'):

c3_2_conv = conv_(c3_1_relu,3,4*nc,8*nc,phase)

c3_2_bn = bn_(c3_2_conv, phase)

c3_2_relu = relu_(c3_2_bn)

c3_pool = max_pool_2x2(c3_2_relu)

with tf.variable_scope('scale4'):

with tf.variable_scope('conv1'):

c4_1_conv = conv_(c3_pool,3,8*nc,8*nc,phase)

c4_1_bn = bn_(c4_1_conv, phase)

c4_1_relu = relu_(c4_1_bn)

with tf.variable_scope('conv2'):

c4_2_conv = conv_(c4_1_relu,3,8*nc,8*nc,phase)

c4_2_bn = bn_(c4_2_conv, phase)

c4_2_relu = relu_(c4_2_bn)

c4_pool = max_pool_2x2(c4_2_relu)

with tf.variable_scope('scale5'):

with tf.variable_scope('conv1'):

c5_1_conv = conv_(c4_pool,3,8*nc,8*nc,phase)

c5_1_bn = bn_(c5_1_conv, phase)

c5_1_relu = relu_(c5_1_bn)

with tf.variable_scope('conv2'):

c5_2_conv = conv_(c5_1_relu,3,8*nc,8*nc,phase)

c5_2_bn = bn_(c5_2_conv, phase)

c5_2_relu = relu_(c5_2_bn)

c5_pool = max_pool_2x2(c5_2_relu)

with tf.variable_scope('scale6'):

with tf.variable_scope('conv1'):

c6_1_conv = conv_(c5_pool,3,8*nc,8*nc,phase)

c6_1_bn = bn_(c6_1_conv, phase)

c6_1_relu = relu_(c6_1_bn)

with tf.variable_scope('conv2'):

c6_2_conv = conv_(c6_1_relu,3,8*nc,8*nc,phase)

c6_2_bn = bn_(c6_2_conv, phase)

c6_2_relu = relu_(c6_2_bn)

with tf.variable_scope('up_minus1'):

with tf.variable_scope('deconv1'):

up_minus1ss = deconv_(c6_2_relu, 8*nc, 8*nc, phase)

with tf.variable_scope('up0'):

up0_1 = tf.concat([up_minus1ss, c5_2_relu], axis=3)

with tf.variable_scope('conv1'):

up0_1_conv = conv_(up0_1,3,16*nc,8*nc,phase)

up0_1_2 = bn_(up0_1_conv,phase)

up0_1_relu = relu_(up0_1_2)

with tf.variable_scope('conv2'):

up0_conv = conv_(up0_1_relu,3,8*nc,8*nc,phase)

up0_2 = bn_(up0_conv,phase)

up0_relu = relu_(up0_2)

with tf.variable_scope('deconv'):

up0 = deconv_(up0_relu, 8*nc, 8*nc, phase)

with tf.variable_scope('up1'):

up1_1 = tf.concat([up0, c4_2_relu], axis=3)

with tf.variable_scope('conv1'):

up1_1_conv = conv_(up1_1,3,16*nc,8*nc,phase)

up1_1_2 = bn_(up1_1_conv,phase)

up1_1_relu = relu_(up1_1_2)

with tf.variable_scope('conv2'):

up1_conv = conv_(up1_1_relu,3,8*nc,8*nc,phase)

up1_2 = bn_(up1_conv,phase)

up1_relu = relu_(up1_2)

with tf.variable_scope('deconv'):

up1 = deconv_(up1_relu, 8*nc, 8*nc, phase)

with tf.variable_scope('up2'):

up2_1 = tf.concat([up1, c3_2_relu], axis=3)

with tf.variable_scope('conv1'):

up2_1_conv = conv_(up2_1,3,16*nc,8*nc,phase)

up2_1_2 = bn_(up2_1_conv,phase)

up2_1_relu = relu_(up2_1_2)

with tf.variable_scope('conv2'):

up2_conv = conv_(up2_1_relu,3,8*nc,8*nc,phase)

up2_2 = bn_(up2_conv,phase)

up2_relu = relu_(up2_2)

with tf.variable_scope('deconv'):

up2 = deconv_(up2_relu, 8*nc, 8*nc, phase)

with tf.variable_scope('up3'):

up3_1 = tf.concat([up2, c2_2_relu], axis=3)

with tf.variable_scope('conv1'):

up3_1_conv = conv_(up3_1,3,12*nc,4*nc,phase)

up3_1_2 = bn_(up3_1_conv,phase)

up3_1_relu = relu_(up3_1_2)

with tf.variable_scope('conv2'):

up3_conv = conv_(up3_1_relu,3,4*nc,4*nc,phase)

up3_2 = bn_(up3_conv,phase)

up3_relu = relu_(up3_2)

with tf.variable_scope('deconv'):

up3 = deconv_(up3_relu, 4*nc, 4*nc, phase)

with tf.variable_scope('up4'):

up4_1 = tf.concat([up3, c1_2_relu], axis=3)

with tf.variable_scope('conv1'):

up4_1_conv = conv_(up4_1,3,6*nc,2*nc,phase)

up4_1_2 = bn_(up4_1_conv,phase)

up4_1_relu = relu_(up4_1_2)

with tf.variable_scope('conv2'):

up4_conv = conv_(up4_1_relu,3,2*nc,2*nc,phase)

up4_2 = bn_(up4_conv,phase)

up4_relu = relu_(up4_2)

with tf.variable_scope('conv3'):

up4 = conv_(up4_1_relu,3,2*nc,nclass,phase)

resultnew=tf.nn.softmax(up4)

return up4, resultnew

def model_apply(sess,result,input_img,phase,tI,ins,num_gpu):

wI=np.zeros([ins,ins])

pmap=np.zeros([tI.shape[0],tI.shape[1],nclass-1])

avI=np.zeros([tI.shape[0],tI.shape[1],nclass-1])

for i in range(ins):

for j in range(ins):

dx=min(i,ins-1-i)

dy=min(j,ins-1-j)

d=min(dx,dy)+1

wI[i,j]=d;

wI = wI/wI.max()

avk = 16

nrotate = 4

for i1 in range(math.ceil(float(avk)*(float(tI.shape[0])-float(ins))/float(ins))+1):

for j1 in range(math.ceil(float(avk)*(float(tI.shape[1])-float(ins))/float(ins))+1):

insti=math.floor(float(i1)*float(ins)/float(avk))

instj=math.floor(float(j1)*float(ins)/float(avk))

inedi=insti+ins

inedj=instj+ins

if inedi>tI.shape[0]:

inedi=tI.shape[0]

insti=inedi-ins

if inedj>tI.shape[1]:

inedj=tI.shape[1]

instj=inedj-ins

print(insti,inedi,instj,inedj)

small_pmap=np.zeros([1,ins,ins,nclass])

feed_image=np.zeros([num_gpu*nrotate,ins,ins,imageType])

for i in range(num_gpu):

for j in range(nrotate):

small_in = tI[insti:inedi,instj:inedj]

feed_image[i*nrotate+j,:,:,:] = np.rot90(small_in, j)

small_out = sess.run(result,feed_dict={input_img:feed_image, phase: False})

for i in range(num_gpu):

for j in range(nrotate):

small_pmap = small_pmap + np.rot90(small_out[i,j,:,:,:],-j)

small_pmap = small_pmap / nrotate / num_gpu

for i in range(nclass-1):

pmap[insti:inedi,instj:inedj,i] += np.multiply(small_pmap[0,:,:,i+1],wI)

avI[insti:inedi,instj:inedj,i] += wI

return np.divide(pmap,np.max(pmap))

def spatial_dropout(x, phase):

d = tf.shape(x)

x_drop = tf.layers.dropout(x, noise_shape=[d[0],1,1,d[3]], training=phase)

return x_drop

def main(_):

for file in os.listdir(logdir):

print('removing '+os.path.join(logdir,file))

os.remove(os.path.join(logdir,file))

np.random.seed(1)

tf.set_random_seed(1)

train_imgs, label_imgs = load_data(imgdir)

#print (np.unique(label_imgs))

#asd

#test_imgs = load_test(imgdir)

#print('test:',len(test_imgs))

count = [0.]*nclass

for lI in label_imgs:

for i in range(nclass):

ok = np.equal(lI,float(i)*interv/255)

count[i] += ok.sum()

count = count/sum(count)

model_loc = FLAGS.model

testing_dir = FLAGS.test_dir

nc = 64

input_img = tf.placeholder(tf.float32, [None,ins,ins,imageType])

phase = tf.placeholder(tf.bool, name='phase')

output_gt = tf.placeholder(tf.int32, [None, ins, ins])

lr = tf.placeholder(tf.float32, name='lr')

tf.summary.image('input_img',input_img)

tf.summary.image('output_gt',tf.cast(tf.expand_dims(output_gt,axis=-1),tf.float32))

use_gpus = FLAGS.gpu.split(',')

print(use_gpus)

errors = []

results = []

input_img_list = tf.split(input_img,num_or_size_splits=int(len(use_gpus)),axis=0)

output_gt_list = tf.split(output_gt,num_or_size_splits=int(len(use_gpus)),axis=0)

for model_id in range(len(use_gpus)):

print(model_id)

cur_score, cur_results = model(input_img_list[model_id],phase,nc,model_id,int(use_gpus[model_id]))

losses = []

loss = (1.0)*tf.reduce_mean(tf.nn.sparse_softmax_cross_entropy_with_logits(logits = cur_score,labels =output_gt))

losses.append(loss)

print('tf.add')

cur_error = tf.add_n(losses)

tf.summary.scalar('error%d' % (model_id), cur_error)

with tf.variable_scope('result%d' % (model_id)):

result = cur_results

for i in range(1,nclass):

tf.summary.image('result_%d_%d' % (model_id, i), tf.expand_dims(result[:,:,:,i-1],axis=-1))

errors.append(cur_error)

results.append(result)

update_ops = tf.get_collection(tf.GraphKeys.UPDATE_OPS)

with tf.variable_scope('result%d' % (model_id)):

result = tf.stack(results,axis=0)

with tf.control_dependencies(update_ops):

bn_result = tf.identity(result)

train_ops = []

for model_id in range(len(use_gpus)):

with tf.device('/gpu:%d' % int(use_gpus[model_id])):

if (model_id==0):

train_step = tf.train.AdamOptimizer(lr).minimize(errors[model_id])

train_ops.append(train_step)

else:

train_step = tf.train.AdamOptimizer(lr).minimize(errors[model_id])

train_ops.append(train_step)

config = tf.ConfigProto()

config.gpu_options.allow_growth = True

config.allow_soft_placement=True

sess = tf.Session(config=config)

merged = tf.summary.merge_all()

train_writer = tf.summary.FileWriter(logdir,

sess.graph)

saver = tf.train.Saver(max_to_keep=100000000)

if (len(model_loc)==0):

print('Init model from scratch')

sess.run(tf.global_variables_initializer())

else:

print('Load model: ' + model_loc)

saver.restore(sess, model_loc)

sess.run(tf.variables_initializer([x for x in tf.global_variables() if 'Adam' in x.name]))

total_parameters = 0

for variable in tf.trainable_variables():

shape = variable.get_shape()

variable_parametes = 1

for dim in shape:

variable_parametes *= dim.value

total_parameters += variable_parametes

print(total_parameters)

if (len(testing_dir)>0):

print('Applying on: ' + testing_dir)

for file in os.listdir(testing_dir):

if file.endswith(".png"):

print(file)

testI=imread(os.path.join(testing_dir,file))

testI=skimage.img_as_float(testI)

pmap = model_apply(sess,result,input_img,phase,testI,ins,len(use_gpus))

for k in range(1,nclass):

pmap_img = np.zeros([testI.shape[0],testI.shape[1]],dtype='uint8')

pmap_img[:,:] = pmap[:,:,k]*255

img_save_dir = '/home/lyang5/tfsave/c' + str(k) + '_' + file

imsave(img_save_dir,pmap_img)

else:

cur_lr = 5e-4

for i in range(50010):

#print (np.unique(label_imgs))

#asd

train_batch, label_batch = get_batch(train_imgs,label_imgs,batch_size*len(use_gpus))

print(np.unique(label_batch))

if i==30000:

cur_lr = 5e-5

info = sess.run([merged,lr,bn_result]+errors+train_ops, feed_dict={

input_img:train_batch, output_gt: label_batch, phase: True, lr: cur_lr})

summary = info[0]

lr_val = info[1]

error_val = info[3:3+len(use_gpus)]

print(i,lr_val)

print(error_val)

train_writer.add_summary(summary, i)

if (i>0) and (i%10000 == 0):

checkpoint_name = os.path.join(resultdir, 'model_' + str(i) + '.ckpt')

print('Saving model: ' + checkpoint_name)

saver.save(sess,checkpoint_name)

for file in os.listdir(os.path.join(imgdir,'data/test/')):

if file.endswith(".png"):

print(file)

testI=imread(os.path.join(imgdir,'data/test/',file))

#testI=testI[:,:,0]

#if (len(testI.shape)==3):

\# testI=testI[:,:,0]

if (len(testI.shape)==2):

testI=np.expand_dims(testI,axis=2)

testI=skimage.img_as_float(testI)

#testI=rescale(testI,0.5,order=0)

print(testI.max())

pmap = model_apply(sess,result,input_img,phase,testI,ins,len(use_gpus))

for k in range(nclass-1):

pmap_img = np.zeros([testI.shape[0],testI.shape[1]],dtype='uint8')

pmap_img[:,:] = pmap[:,:,k]*255

print(pmap_img.max())

print(pmap_img.min())

img_save_dir = resultdir + '/iter' + str(i) +'_c' + str(k+1) + '_' + file

imsave(img_save_dir,pmap_img)

if __name__ == '__main__':

parser = argparse.ArgumentParser()

parser.add_argument('-model', type=str, default='',

help='model location')

parser.add_argument('-test_dir', type=str, default='',

help='model location')

parser.add_argument('-gpu', type=str, default='0',

help='use gpu')

FLAGS, unparsed = parser.parse_known_args()

tf.app.run(main=main, argv=[sys.argv[0]] + unparsed)

#### 3. Source code for segmenting using the trained model:

from __future__ import absolute_import

from __future__ import division

from __future__ import print_function

import argparse

import sys

import math

import numpy as np

import os

import random

import time

import tensorflow as tf

import tensorflow.contrib.slim as slim

from tensorflow.contrib.slim.python.slim.nets import resnet_v2

from tensorflow.python.ops import control_flow_ops

import skimage

from skimage.io import imread

from skimage.io import imsave

from skimage.transform import rotate

from skimage.transform import rescale

from operator import itemgetter

from skimage.morphology import label

from scipy.ndimage.morphology import binary_dilation

FLAGS = None

imageType = 1

ous = 192

ins = 192

interv = 255

nclass =2

nclass4= 2

batch_size = 8

test_batch_size = 0

#homedir = '/home/lyang5/'

homedir = '/afs/crc.nd.edu/user/y/yzhang29/Mouse_lymph_node/'

logdir = homedir + 'tftrain'

resultdir = homedir + 'RESULTS'

imgdir = '/afs/crc.nd.edu/user/y/yzhang29/Mouse_lymph_node/'

def load_data(file_dir):

train_imgs = []

label_imgs = []

count = 0

for file in os.listdir(os.path.join(file_dir,'data/unlabeled/')):

if file.endswith(".png"):

tI=imread(os.path.join(file_dir,'data/unlabeled/',file))

#if (len(tI.shape)==3):

\# tI=tI[:,:,0]

tI=skimage.img_as_float(tI)

#tI=rescale(tI,0.5,order=0)

if (len(tI.shape)==2):

tI=np.expand_dims(tI,axis=2)

train_imgs.append(tI)

lI=imread(os.path.join(file_dir,'data/mask727/',file))

#lI=skimage.img_as_float(lI)

print (np.unique(lI))

if (len(lI.shape)==3):

lI=lI[:,:,0]

label_imgs.append(lI)

for file in os.listdir(os.path.join(file_dir,'data/train/')):

if file.endswith(".png"):

tI=imread(os.path.join(file_dir,'data/train/',file))

#if (len(tI.shape)==3):

\# tI=tI[:,:,0]

tI=skimage.img_as_float(tI)

#tI=rescale(tI,0.5,order=0)

if (len(tI.shape)==2):

tI=np.expand_dims(tI,axis=2)

train_imgs.append(tI)

lI=imread(os.path.join(file_dir,'data/mask/',file))

#lI=skimage.img_as_float(lI)

print (np.unique(lI))

if (len(lI.shape)==3):

lI=lI[:,:,0]

label_imgs.append(lI)

return train_imgs, label_imgs

def load_test(file_dir):

test_imgs = []

for file in os.listdir(os.path.join(file_dir,'raw/')):

if file.endswith(".png"):

tI=imread(os.path.join(file_dir,'raw/',file))

tI=skimage.img_as_float(tI)

#tI=rescale(tI,0.5,order=0)

if (len(tI.shape)==2):

tI=np.expand_dims(tI,axis=2)

test_imgs.append(tI)

return test_imgs

def get_image(train_imgs,label_imgs):

i_idx = random.randint(0, len(train_imgs)-1)

big_tI = train_imgs[i_idx]

big_lI = label_imgs[i_idx]

xs = big_tI.shape[0]

ys = big_tI.shape[1]

stx = random.randint(0,xs-ins)

sty = random.randint(0,ys-ins)

train_sample = np.zeros((ins,ins,imageType))

label_sample = np.zeros((ins,ins))

train_sample = big_tI[stx:stx+ins,sty:sty+ins,:]

label_sample = big_lI[stx:stx+ins,sty:sty+ins]

nrotate = random.randint(0, 3)

train_sample = np.rot90(train_sample, nrotate)

#print (np.unique(label_sample))

label_sample = np.rot90(label_sample, nrotate).astype('uint8')

#print (np.unique(label_sample))

#asd

nflip = random.randint(0, 1)

if nflip:

#print('flip')

train_sample = np.fliplr(train_sample)

label_sample = np.fliplr(label_sample)

return train_sample, label_sample

def get_image_balanced_pos(train_imgs,label_imgs):

i_idx = random.randint(0, len(train_imgs)-1)

big_tI = train_imgs[i_idx]

big_lI = label_imgs[i_idx]

xs = big_tI.shape[0]

ys = big_tI.shape[1]

flag=1

print ('get_pos:')

while flag==1:

stx = random.randint(0,xs-ins)

sty = random.randint(0,ys-ins)

train_sample = np.zeros((ins,ins,imageType))

label_sample = np.zeros((ins,ins))

train_sample = big_tI[stx:stx+ins,sty:sty+ins,:]

label_sample = big_lI[stx:stx+ins,sty:sty+ins]

#print (label_sample[int(ins/2),int(ins/2)])

if label_sample[int(ins/2),int(ins/2)]==40.0:

flag==0

nrotate = random.randint(0, 3)

train_sample = np.rot90(train_sample, nrotate)

#print (np.unique(label_sample))

label_sample = np.rot90(label_sample, nrotate).astype('uint8')

#print (np.unique(label_sample))

#asd

nflip = random.randint(0, 1)

if nflip:

#print('flip')

train_sample = np.fliplr(train_sample)

label_sample = np.fliplr(label_sample)

return train_sample, label_sample

def get_image_balanced_neg(train_imgs,label_imgs):

i_idx = random.randint(0, len(train_imgs)-1)

big_tI = train_imgs[i_idx]

big_lI = label_imgs[i_idx]

xs = big_tI.shape[0]

ys = big_tI.shape[1]

flag=1

print ('get_neg:')

while flag==1:

stx = random.randint(0,xs-ins)

sty = random.randint(0,ys-ins)

train_sample = np.zeros((ins,ins,imageType))

label_sample = np.zeros((ins,ins))

train_sample = big_tI[stx:stx+ins,sty:sty+ins,:]

label_sample = big_lI[stx:stx+ins,sty:sty+ins]

if label_sample[int(ins/2),int(ins/2)]!=40.0:

flag==0

nrotate = random.randint(0, 3)

train_sample = np.rot90(train_sample, nrotate)

#print (np.unique(label_sample))

label_sample = np.rot90(label_sample, nrotate).astype('uint8')

#print (np.unique(label_sample))

#asd

nflip = random.randint(0, 1)

if nflip:

#print('flip')

train_sample = np.fliplr(train_sample)

label_sample = np.fliplr(label_sample)

return train_sample, label_sample

def get_batch(train_imgs,label_imgs,batch_size):

train_samples = np.zeros((batch_size,ins,ins,imageType))

label_samples = np.zeros((batch_size,ins,ins))

for i in range(int(batch_size)):

train_sample, label_sample = get_image(train_imgs,label_imgs)

train_samples[i,:,:,:] = train_sample

label_samples[i,:,:] = label_sample/interv

return train_samples, label_samples.astype('int32')

def get_test_batch(test_imgs,batch_size):

test_samples = np.zeros((batch_size,ins,ins,imageType))

label_samples = np.zeros((batch_size,ins,ins))

for i in range(batch_size):

test_sample, label_sample = get_test_image(test_imgs)

test_samples[i,:,:,:] = test_sample

label_samples[i,:,:] = 2

return test_samples, label_samples.astype('int32')

import tensorflow as tf

import math

def weight_variable(shape,stdv):

initial = tf.get_variable("weights", shape=shape, initializer=tf.random_normal_initializer(stddev=stdv))

return initial

def bias_variable(shape):

initial = tf.get_variable("bias", shape=shape, initializer=tf.constant_initializer(value=0.0))

return initial

def conv_(x,k,nin,nout,phase,s=1,d=1):

stdv = math.sqrt(2/(nin*k*k))

return tf.layers.conv2d(x, nout, k, strides=[s,s], dilation_rate=[d,d], padding='same',

kernel_initializer = tf.random_normal_initializer(stddev=stdv),

use_bias=False)

def convnn_(x,k,s,nin,nout,phase):

print('convnn')

stdv = math.sqrt(1/(nin*k*k))

return tf.layers.conv2d(x, nout, k, strides=[s,s], padding='same',

kernel_initializer = tf.random_uniform_initializer(minval=-stdv, maxval=stdv,dtype=tf.float32),

use_bias=False)

def bn_(x,phase):

bn_result = tf.layers.batch_normalization(x, momentum=0.9, epsilon = 1e-5,

gamma_initializer = tf.random_uniform_initializer(minval=0.0, maxval=1.0,dtype=tf.float32),

training = phase)

return bn_result

def relu_(x):

return tf.nn.relu(x)

def cbr_(x,k,nin,nout,phase,s=1,d=1):

x_conv = conv_(x,k,nin,nout,phase,s,d)

x_bn = bn_(x_conv,phase)

x_relu = relu_(x_bn)

return x_relu

def bottleneck(x, nin, nout, phase, s=1, d=1):

if (nin != nout) or (s != 1):

print('conv_skip')

with tf.variable_scope('skip'):

skip_conv = conv_(x,1,nin,nout,phase,s=s)

skip = bn_(skip_conv,phase)

else:

skip = x

with tf.variable_scope('conv1'):

c1_conv = conv_(x,3,nin,nout,phase,s=s,d=d)

c1_bn = bn_(c1_conv,phase)

c1_relu = relu_(c1_bn)

with tf.variable_scope('conv2'):

c2_conv = conv_(c1_relu,3,nout,nout,phase,d=d)

c2_bn = bn_(c2_conv,phase)

out = skip + c2_bn

out_relu = relu_(out)

return out_relu

def stack(x, nin, nout, nblock, phase, s=1, d=1, new_level=True):

for i in range(nblock):

with tf.variable_scope('block%d' % (i)):

if i==0:

x = bottleneck(x,nin,nout,phase,s=s,d=d)

else:

x = bottleneck(x,nout,nout,phase,d=d)

return x

def max_pool_2x2(x):

print('new pool')

return tf.layers.max_pooling2d(x,[2,2],[2,2])

def avg_pool_2x2(x):

print('avg')

return tf.nn.avg_pool(x, ksize=[1, 2, 2, 1],

strides=[1, 2, 2, 1], padding='SAME')

def show(x,k):

x_sm = tf.nn.softmax(x)

x_sl = tf.unstack(x_sm,num=nclass,axis=-1)[k]

return tf.expand_dims(x_sl,-1)

def deconv_(x, nin, nout, phase):

stdv = math.sqrt(2/(nin*2*2))

conv_r = tf.layers.conv2d_transpose(x, nout, 4, strides=[2,2], padding='same',

kernel_initializer = tf.random_normal_initializer(stddev=stdv),

use_bias=False)

bn_r = bn_(conv_r,phase)

relu_r = relu_(bn_r)

return relu_r

def deconv1_(x, nin, num_up, phase):

cur_in = nin

cur_out=(2**(num_up-1))

for i in range(num_up):

with tf.variable_scope('up%d' % (i)):

x = deconv_(x,cur_in,cur_out,phase)

cur_in = cur_out

cur_out = math.floor(cur_out/2)

return x

def deconvn_(x, nin, nout, num_up, phase):

cur_in = nin

cur_out=((2**(num_up-1))*nout)

for i in range(num_up):

with tf.variable_scope('up%d' % (i)):

x = deconv_(x,cur_in,cur_out,phase)

cur_in = cur_out

cur_out = math.floor(cur_out/2)

return x

def deconv2_(x, nin, num_up, phase):

cur_in = nin

cur_out=(2**(num_up))

for i in range(num_up):

with tf.variable_scope('up%d' % (i)):

x = deconv_(x,cur_in,cur_out,phase)

cur_in = cur_out

cur_out = math.floor(cur_out/2)

return x

def model(input_img,phase,nc,id,gpu):

with tf.device('/gpu:%d' % (gpu)):

with tf.variable_scope('model%d' % (id)):

with tf.variable_scope('scale1'):

with tf.variable_scope('conv1'):

c1_1_conv = conv_(input_img, 3,imageType,nc,phase)

c1_1_bn = bn_(c1_1_conv, phase)

c1_1_relu = relu_(c1_1_bn)

with tf.variable_scope('conv2'):

c1_2_conv = conv_(c1_1_relu,3,nc,2*nc,phase)

c1_2_bn = bn_(c1_2_conv, phase)

c1_2_relu = relu_(c1_2_bn)

c1_pool = max_pool_2x2(c1_2_relu)

with tf.variable_scope('scale2'):

with tf.variable_scope('conv1'):

c2_1_conv = conv_(c1_pool,3,2*nc,2*nc,phase)

c2_1_bn = bn_(c2_1_conv, phase)

c2_1_relu = relu_(c2_1_bn)

with tf.variable_scope('conv2'):

c2_2_conv = conv_(c2_1_relu,3,2*nc,4*nc,phase)

c2_2_bn = bn_(c2_2_conv, phase)

c2_2_relu = relu_(c2_2_bn)

c2_pool = max_pool_2x2(c2_2_relu)

with tf.variable_scope('scale3'):

with tf.variable_scope('conv1'):

c3_1_conv = conv_(c2_pool,3,4*nc,4*nc,phase)

c3_1_bn = bn_(c3_1_conv, phase)

c3_1_relu = relu_(c3_1_bn)

with tf.variable_scope('conv2'):

c3_2_conv = conv_(c3_1_relu,3,4*nc,8*nc,phase)

c3_2_bn = bn_(c3_2_conv, phase)

c3_2_relu = relu_(c3_2_bn)

c3_pool = max_pool_2x2(c3_2_relu)

with tf.variable_scope('scale4'):

with tf.variable_scope('conv1'):

c4_1_conv = conv_(c3_pool,3,8*nc,8*nc,phase)

c4_1_bn = bn_(c4_1_conv, phase)

c4_1_relu = relu_(c4_1_bn)

with tf.variable_scope('conv2'):

c4_2_conv = conv_(c4_1_relu,3,8*nc,8*nc,phase)

c4_2_bn = bn_(c4_2_conv, phase)

c4_2_relu = relu_(c4_2_bn)

c4_pool = max_pool_2x2(c4_2_relu)

with tf.variable_scope('scale5'):

with tf.variable_scope('conv1'):

c5_1_conv = conv_(c4_pool,3,8*nc,8*nc,phase)

c5_1_bn = bn_(c5_1_conv, phase)

c5_1_relu = relu_(c5_1_bn)

with tf.variable_scope('conv2'):

c5_2_conv = conv_(c5_1_relu,3,8*nc,8*nc,phase)

c5_2_bn = bn_(c5_2_conv, phase)

c5_2_relu = relu_(c5_2_bn)

c5_pool = max_pool_2x2(c5_2_relu)

with tf.variable_scope('scale6'):

with tf.variable_scope('conv1'):

c6_1_conv = conv_(c5_pool,3,8*nc,8*nc,phase)

c6_1_bn = bn_(c6_1_conv, phase)

c6_1_relu = relu_(c6_1_bn)

with tf.variable_scope('conv2'):

c6_2_conv = conv_(c6_1_relu,3,8*nc,8*nc,phase)

c6_2_bn = bn_(c6_2_conv, phase)

c6_2_relu = relu_(c6_2_bn)

with tf.variable_scope('up_minus1'):

with tf.variable_scope('deconv1'):

up_minus1ss = deconv_(c6_2_relu, 8*nc, 8*nc, phase)

with tf.variable_scope('up0'):

up0_1 = tf.concat([up_minus1ss, c5_2_relu], axis=3)

with tf.variable_scope('conv1'):

up0_1_conv = conv_(up0_1,3,16*nc,8*nc,phase)

up0_1_2 = bn_(up0_1_conv,phase)

up0_1_relu = relu_(up0_1_2)

with tf.variable_scope('conv2'):

up0_conv = conv_(up0_1_relu,3,8*nc,8*nc,phase)

up0_2 = bn_(up0_conv,phase)

up0_relu = relu_(up0_2)

with tf.variable_scope('deconv'):

up0 = deconv_(up0_relu, 8*nc, 8*nc, phase)

with tf.variable_scope('up1'):

up1_1 = tf.concat([up0, c4_2_relu], axis=3)

with tf.variable_scope('conv1'):

up1_1_conv = conv_(up1_1,3,16*nc,8*nc,phase)

up1_1_2 = bn_(up1_1_conv,phase)

up1_1_relu = relu_(up1_1_2)

with tf.variable_scope('conv2'):

up1_conv = conv_(up1_1_relu,3,8*nc,8*nc,phase)

up1_2 = bn_(up1_conv,phase)

up1_relu = relu_(up1_2)

with tf.variable_scope('deconv'):

up1 = deconv_(up1_relu, 8*nc, 8*nc, phase)

with tf.variable_scope('up2'):

up2_1 = tf.concat([up1, c3_2_relu], axis=3)

with tf.variable_scope('conv1'):

up2_1_conv = conv_(up2_1,3,16*nc,8*nc,phase)

up2_1_2 = bn_(up2_1_conv,phase)

up2_1_relu = relu_(up2_1_2)

with tf.variable_scope('conv2'):

up2_conv = conv_(up2_1_relu,3,8*nc,8*nc,phase)

up2_2 = bn_(up2_conv,phase)

up2_relu = relu_(up2_2)

with tf.variable_scope('deconv'):

up2 = deconv_(up2_relu, 8*nc, 8*nc, phase)

with tf.variable_scope('up3'):

up3_1 = tf.concat([up2, c2_2_relu], axis=3)

with tf.variable_scope('conv1'):

up3_1_conv = conv_(up3_1,3,12*nc,4*nc,phase)

up3_1_2 = bn_(up3_1_conv,phase)

up3_1_relu = relu_(up3_1_2)

with tf.variable_scope('conv2'):

up3_conv = conv_(up3_1_relu,3,4*nc,4*nc,phase)

up3_2 = bn_(up3_conv,phase)

up3_relu = relu_(up3_2)

with tf.variable_scope('deconv'):

up3 = deconv_(up3_relu, 4*nc, 4*nc, phase)

with tf.variable_scope('up4'):

up4_1 = tf.concat([up3, c1_2_relu], axis=3)

with tf.variable_scope('conv1'):

up4_1_conv = conv_(up4_1,3,6*nc,2*nc,phase)

up4_1_2 = bn_(up4_1_conv,phase)

up4_1_relu = relu_(up4_1_2)

with tf.variable_scope('conv2'):

up4_conv = conv_(up4_1_relu,3,2*nc,2*nc,phase)

up4_2 = bn_(up4_conv,phase)

up4_relu = relu_(up4_2)

with tf.variable_scope('conv3'):

up4 = conv_(up4_1_relu,3,2*nc,nclass,phase)

resultnew=tf.nn.softmax(up4)

return up4, resultnew

def model_apply(sess,result,input_img,phase,tI,ins,num_gpu):

wI=np.zeros([ins,ins])

pmap=np.zeros([tI.shape[0],tI.shape[1],nclass-1])

avI=np.zeros([tI.shape[0],tI.shape[1],nclass-1])

for i in range(ins):

for j in range(ins):

dx=min(i,ins-1-i)

dy=min(j,ins-1-j)

d=min(dx,dy)+1

wI[i,j]=d;

wI = wI/wI.max()

avk = 16

nrotate = 4

for i1 in range(math.ceil(float(avk)*(float(tI.shape[0])-float(ins))/float(ins))+1):

for j1 in range(math.ceil(float(avk)*(float(tI.shape[1])-float(ins))/float(ins))+1):

insti=math.floor(float(i1)*float(ins)/float(avk))

instj=math.floor(float(j1)*float(ins)/float(avk))

inedi=insti+ins

inedj=instj+ins

if inedi>tI.shape[0]:

inedi=tI.shape[0]

insti=inedi-ins

if inedj>tI.shape[1]:

inedj=tI.shape[1]

instj=inedj-ins

print(insti,inedi,instj,inedj)

small_pmap=np.zeros([1,ins,ins,nclass])

feed_image=np.zeros([num_gpu*nrotate,ins,ins,imageType])

for i in range(num_gpu):

for j in range(nrotate):

small_in = tI[insti:inedi,instj:inedj]

feed_image[i*nrotate+j,:,:,:] = np.rot90(small_in, j)

small_out = sess.run(result,feed_dict={input_img:feed_image, phase: False})

for i in range(num_gpu):

for j in range(nrotate):

small_pmap = small_pmap + np.rot90(small_out[i,j,:,:,:],-j)

small_pmap = small_pmap / nrotate / num_gpu

for i in range(nclass-1):

pmap[insti:inedi,instj:inedj,i] += np.multiply(small_pmap[0,:,:,i+1],wI)

avI[insti:inedi,instj:inedj,i] += wI

return np.divide(pmap,np.max(pmap))

def spatial_dropout(x, phase):

d = tf.shape(x)

x_drop = tf.layers.dropout(x, noise_shape=[d[0],1,1,d[3]], training=phase)

return x_drop

def main(_):

for file in os.listdir(logdir):

print('removing '+os.path.join(logdir,file))

os.remove(os.path.join(logdir,file))

np.random.seed(1)

tf.set_random_seed(1)

train_imgs, label_imgs = load_data(imgdir)

#print (np.unique(label_imgs))

#asd

#test_imgs = load_test(imgdir)

#print('test:',len(test_imgs))

count = [0.]*nclass

for lI in label_imgs:

for i in range(nclass):

ok = np.equal(lI,float(i)*interv/255)

count[i] += ok.sum()

count = count/sum(count)

model_loc = FLAGS.model

testing_dir = FLAGS.test_dir

nc = 64

input_img = tf.placeholder(tf.float32, [None,ins,ins,imageType])

phase = tf.placeholder(tf.bool, name='phase')

output_gt = tf.placeholder(tf.int32, [None, ins, ins])

lr = tf.placeholder(tf.float32, name='lr')

tf.summary.image('input_img',input_img)

tf.summary.image('output_gt',tf.cast(tf.expand_dims(output_gt,axis=-1),tf.float32))

use_gpus = FLAGS.gpu.split(',')

print(use_gpus)

errors = []

results = []

input_img_list = tf.split(input_img,num_or_size_splits=int(len(use_gpus)),axis=0)

output_gt_list = tf.split(output_gt,num_or_size_splits=int(len(use_gpus)),axis=0)

for model_id in range(len(use_gpus)):

print(model_id)

cur_score, cur_results = model(input_img_list[model_id],phase,nc,model_id,int(use_gpus[model_id]))

losses = []

loss = (1.0)*tf.reduce_mean(tf.nn.sparse_softmax_cross_entropy_with_logits(logits = cur_score,labels =output_gt))

losses.append(loss)

print('tf.add')

cur_error = tf.add_n(losses)

tf.summary.scalar('error%d' % (model_id), cur_error)

with tf.variable_scope('result%d' % (model_id)):

result = cur_results

for i in range(1,nclass):

tf.summary.image('result_%d_%d' % (model_id, i), tf.expand_dims(result[:,:,:,i-1],axis=-1))

errors.append(cur_error)

results.append(result)

update_ops = tf.get_collection(tf.GraphKeys.UPDATE_OPS)

with tf.variable_scope('result%d' % (model_id)):

result = tf.stack(results,axis=0)

with tf.control_dependencies(update_ops):

bn_result = tf.identity(result)

train_ops = []

for model_id in range(len(use_gpus)):

with tf.device('/gpu:%d' % int(use_gpus[model_id])):

if (model_id==0):

train_step = tf.train.AdamOptimizer(lr).minimize(errors[model_id])

train_ops.append(train_step)

else:

train_step = tf.train.AdamOptimizer(lr).minimize(errors[model_id])

train_ops.append(train_step)

config = tf.ConfigProto()

config.gpu_options.allow_growth = True

config.allow_soft_placement=True

sess = tf.Session(config=config)

merged = tf.summary.merge_all()

train_writer = tf.summary.FileWriter(logdir,

sess.graph)

saver = tf.train.Saver(max_to_keep=100000000)

#if (len(model_loc)==0):

\# print('Init model from scratch')

\# sess.run(tf.global_variables_initializer())

#else:

#print('Load model: ' + model_loc)

saver.restore(sess, '/afs/crc.nd.edu/user/y/yzhang29/Mouse_lymph_node/tfsave_unet/model_30000.ckpt')

sess.run(tf.variables_initializer([x for x in tf.global_variables() if 'Adam' in x.name]))

total_parameters = 0

for variable in tf.trainable_variables():

shape = variable.get_shape()

variable_parametes = 1

for dim in shape:

variable_parametes *= dim.value

total_parameters += variable_parametes

print(total_parameters)

if (len(testing_dir)>0):

print('Applying on: ' + testing_dir)

for file in os.listdir(testing_dir):

if file.endswith(".png"):

print(file)

testI=imread(os.path.join(testing_dir,file))

testI=skimage.img_as_float(testI)

pmap = model_apply(sess,result,input_img,phase,testI,ins,len(use_gpus))

for k in range(1,nclass):

pmap_img = np.zeros([testI.shape[0],testI.shape[1]],dtype='uint8')

pmap_img[:,:] = pmap[:,:,k]*255

img_save_dir = '/home/lyang5/tfsave/c' + str(k) + '_' + file

imsave(img_save_dir,pmap_img)

else:

cur_lr = 5e-4

for i in range(1):

#print (np.unique(label_imgs))

#asd

if (i==0) and (i%10000 == 0):

#checkpoint_name = os.path.join(resultdir, 'model_' + str(i) + '.ckpt')

#print('Saving model: ' + checkpoint_name)

#saver.save(sess,checkpoint_name)

for file in os.listdir('/afs/crc.nd.edu/user/y/yzhang29/LN_w_tumor_right/'):

counter=0

for filefile in (os.listdir('/afs/crc.nd.edu/user/y/yzhang29/LN_w_tumor_right/' + file)):

if filefile.endswith(".png"):

print ('~')

else:

img_save_dir = resultdir + '/' + file + '_' + filefile

if os.path.isfile(img_save_dir):

counter=counter+1

if (counter%10)==0:

print(file)

testI=imread('/afs/crc.nd.edu/user/y/yzhang29/LN_w_tumor_right/'+file+'/'+filefile)

#testI=testI[:,:,0]

#if (len(testI.shape)==3):

\# testI=testI[:,:,0]

if (len(testI.shape)==2):

testI=np.expand_dims(testI,axis=2)

testI=skimage.img_as_float(testI)

#testI=rescale(testI,0.5,order=0)

print(testI.max())

pmap = model_apply(sess,result,input_img,phase,testI,ins,len(use_gpus))

for k in range(nclass-1):

pmap_img = np.zeros([testI.shape[0],testI.shape[1]],dtype='uint8')

pmap_img[:,:] = pmap[:,:,k]*255

print(pmap_img.max())

print(pmap_img.min())

img_save_dir = resultdir + '/' + file + '_' + filefile

imsave(img_save_dir,pmap_img)

if __name__ == '__main__':

parser = argparse.ArgumentParser()

parser.add_argument('-model', type=str, default='',

help='model location')

parser.add_argument('-test_dir', type=str, default='',

help='model location')

parser.add_argument('-gpu', type=str, default='0',

help='use gpu')

FLAGS, unparsed = parser.parse_known_args()

tf.app.run(main=main, argv=[sys.argv[0]] + unparsed)
